# Improving the Robustness of Phylogenetic Independent Contrasts: Addressing Abrupt Evolutionary Shifts with Outlier-and Distribution-Guided Correlation

**DOI:** 10.1101/2024.06.16.599156

**Authors:** Zheng-Lin Chen, Rui Huang, Hong-Ji Guo, Deng-Ke Niu

## Abstract

Traditional phylogenetically aware correlation methods perform well under gradual evolutionary processes. However, abrupt evolutionary shifts—or macroevolutionary jumps, characteristic of punctuated evolution—can produce extreme phylogenetically independent contrasts (PIC), leading to inflated false positives or increased false negatives in trait correlation analyses. We introduce O(D)GC (Outlier-and Distribution-Guided Correlation), a flexible workflow that identifies outliers in PICs using a distribution-free boxplot criterion and applies Spearman correlation whenever influential outliers are detected. If no outliers are detected, Pearson correlation is used—automatically for large datasets (*n* ≥ 30), or guided by normality testing in smaller samples. We systematically compared PIC-O(D)GC with five widely applied phylogenetic correlation methods—PIC-Pearson, PIC-MM, PGLS (phylogenetic generalized least squares), MR-PMM (multi-response phylogenetic mixed model), and Corphylo—on 322,000 simulated datasets spanning five evolutionary scenarios (two shift settings: single-trait shifts and dual-trait co-directional jumps; and three no-shift gradual evolution settings), including both fixed-depth and randomly located shifts, tested across 11 shift or noise gradients, three tree sizes (16, 128, 256 tips), and both balanced and random topologies. Overall, PIC-O(D)GC achieved error rates comparable to—or noticeably higher than—those of PIC-MM, while yielding substantially lower error rates than most alternative methods. Under no-shift conditions, it retained power similar to other methods.

Analyses of three empirical datasets likewise showed that PIC-O(D)GC and PIC-MM corrected shift-induced distortions that misled conventional methods. Moreover, PIC-O(D)GC offers a conceptually simple framework and incurs markedly lower computational cost. By design, its correlation-only output provides less mechanistic detail than regression-based approaches like PGLS. However, when paired with PIC diagnostics, this outlier-guided strategy highlights evolutionary jumps, distinguishes coupled from decoupled shifts, and—via clade partitioning or tip pruning—recovers background correlations, offering biologically informative insights into how punctuated events interact with gradual trends in trait evolution.

## Introduction

Phylogenetically closer species tend to exhibit more similar traits due to their shared ancestry, resulting in statistical dependencies among samples (Felsenstein 1985, Garamszegi 2014, Cornwallis and Griffin 2024). Ignoring these statistical dependencies and using conventional statistical methods can lead to erroneous conclusions. To address this challenge, evolutionary biologists have developed a range of phylogenetic comparative methods that explicitly account for these dependencies (Grafen 1989, Lynch 1991, Garamszegi 2014, O’Meara 2016, Cornwallis and Griffin 2024, Dewar et al. 2025).

The earliest method was phylogenetically independent contrasts (PIC) (Felsenstein 1985). The core principle behind PIC is that, while species at the tips of a phylogenetic tree are interdependent due to shared ancestry, the evolutionary changes along each branch can be treated as statistically independent events. In this context, PIC refers to the contrast values calculated between species pairs along the phylogenetic tree, inferring evolutionary changes along pairs of branches and standardizing these changes by branch length (or time) to generate transformed data points. In traditional applications, these PIC values are analyzed using ordinary least squares (OLS) regression or, equivalently under linear assumptions, Pearson correlation, to assess associations between traits. This transformation renders evolutionary data more suitable for conventional statistical methods by removing the phylogenetic non-independence. Thus, PIC can refer both to the contrast values calculated from the phylogeny and to the subsequent OLS regression analysis applied to these values.

Applying OLS regression to PIC values assumes that traits evolve according to a Brownian motion (BM) model, under which trait variation accumulates gradually and randomly over time. While the computation of PIC values does not strictly require a BM model, the statistical justification for treating PIC values as independent and identically distributed (i.i.d.) variables in regression analyses typically relies on the BM assumption or its extensions. Under this workflow, OLS regression through the origin on PIC values yields slope estimates, standard errors, and *p*-values that are numerically identical to those obtained from phylogenetic generalized least squares (PGLS), assuming BM holds (Rohlf 2001, Blomberg et al. 2012). PGLS generalizes this approach by directly modeling the phylogenetic covariance structure through a variance–covariance matrix of the residuals in the linear model, thereby accommodating phylogenetic autocorrelation among observations (Martins and Hansen 1997, Pagel 1997, Pagel 1999). This adjustment helps reduce false positives in hypothesis testing due to shared evolutionary history. Although some PGLS models, such as Pagel’s λ, Ornstein-Uhlenbeck (OU), and Early-burst (EB) models, can handle heterogeneous evolutionary rates (Hansen 1997, Freckleton et al. 2002, Harmon et al. 2010, Ho and Ane 2014), they fundamentally assume gradual trait changes (Landis et al. 2013). This assumption can be problematic when dealing with evolutionary jumps caused by singular events.

During biological evolution, certain branches may experience rare and dramatic trait changes (Uyeda et al. 2011, Landis et al. 2013). Increasingly, studies and data have shown that abrupt evolutionary shifts, or evolutionary jumps, have a significant impact on biodiversity evolution (Landis and Schraiber 2017, Gao and Wu 2022, Sumner et al. 2023). A typical example of this is Felsenstein’s worst-case scenario (Felsenstein 1985, Uyeda et al. 2018), where two traits undergo a significant jump near the root of the phylogenetic tree while evolving independently on other branches. Even if this dramatic change occurred only once, it could still lead to erroneous conclusions of significant correlation when using traditional regression-based PIC analyses or PGLS models (Uyeda et al. 2018).

Recently, more studies have focused on identifying jump events and providing more reasonable explanations for the processes underlying trait evolution. Uyeda et al. (2018) underscore the need for models like the singular events model, which can accommodate the complexities and unpredictabilities inherent in evolutionary biology. Some studies suggest that jump events cause trait changes to exhibit a “fat-tails” distribution, using Levy processes to handle this distribution (Landis et al. 2013, Duchen et al. 2017). Additionally, LASSO-based methods within models like OU have been applied to pinpoint shifts along specific branches of phylogenetic trees (Khabbazian et al. 2016, Bastide et al. 2018, Zhang et al. 2024). Others propose that abrupt shift events lead to outliers, using robust regression to reduce model sensitivity to violations (Slater and Pennell 2014, Adams et al. 2024). A variety of mixed-model approaches have also been explored to address model deviations and trait interdependence (Ives and Helmus 2011, Westoby et al. 2023, Halliwell et al. 2025).

Evolutionary shifts, especially those occurring in branches close to the root of the phylogenetic tree, typically do not manifest directly as outliers in the trait data but instead appear as outliers in the PIC values. As a result, robust statistical methods designed to handle outliers in the original trait data may not be as effective in addressing evolutionary shifts.

Adams et al. (2024) demonstrated that when four different estimators (L1, S, M, and MM) were integrated into both PGLS and PIC analysis, methods like PIC-MM and PIC-S performed significantly better in detecting evolutionary relationships, especially when accounting for outliers. Thus, computing PIC values is essential for effective analysis of the relationships between traits with evolutionary shifts.

Dramatic changes in some evolutionary branches can result in several outliers in the PIC dataset (Uyeda et al. 2018), distorting the statistical results of parametric tests. While nonparametric approaches are generally more robust to such outliers because they do not rely on specific distributional assumptions, they are not routinely incorporated into traditional phylogenetic correlation workflows.

To address this gap, we propose a hybrid workflow, Outlier-Guided Correlation (OGC), for analyzing pairwise trait relationships under diverse evolutionary conditions. The core idea of OGC is to integrate the PIC framework with a data-driven mechanism that evaluates the presence of outliers in trait contrasts and selects an appropriate correlation measure accordingly. For larger datasets, OGC selects Spearman or Pearson correlation depending on whether outliers are detected. For smaller datasets, where non-normality can significantly impact results, the workflow incorporates both normality testing and outlier detection, referred to as Outlier-and Distribution-Guided Correlation (ODGC). Together, these approaches form a unified workflow collectively referred to as O(D)GC, designed to address the challenges posed by varying data conditions.

It should be noted that, whereas regression-based methods such as PGLS are commonly used for both parameter estimation and hypothesis testing, the O(D)GC workflow is intentionally restricted to null hypothesis testing, focusing on whether significant pairwise trait correlations are present. Readers should interpret its results within this scope.

## Materials and Methods

### Phylogenetic Tree Construction and Overview of Simulation Strategy

We categorize the construction of phylogenetic trees into two types. The first is the fixed-balanced tree scenario, where all non-leaf nodes have left and right subtrees with identical topology and depth, forming a fully symmetric structure. In this setting, a single fixed-balanced phylogenetic tree with 128 species is used as the basis for all trait simulations.

Specifically, traits are simulated under combinations of evolutionary scenarios and experimental conditions reflecting different levels of stochastic variation or evolutionary shifts, with 1,000 replicates per condition, using different random seeds while keeping the phylogenetic tree fixed. Consequently, for each trait evolutionary scenario-gradient combination, 1,000 datasets are generated based on the same phylogenetic structure. To assess the influence of tree size, additional analyses were conducted using 16-and 256-species fixed-balanced trees, following the same simulation protocol.

The second category is a constrained random-tree scenario (hereafter referred to as the random-tree scenario), in which a new phylogenetic tree is generated prior to each trait simulation. In this setting, the root ancestor is forced to bifurcate into two subtrees of equal size that share an identical topology. This topology is randomly generated under a coalescent model for each simulation replicate, such that randomness is introduced at the level of subtree topology. For each replicate, a phylogenetic tree is first constructed, followed by trait simulation under a given evolutionary scenario and gradient condition. This pair of operations—tree construction and trait simulation—is repeated 1,000 times using different random seeds, ensuring variation in both tree topology and simulated trait values.

Subsequently, we evaluated the performance of each trait-correlation inference method across all evolutionary scenario–gradient combinations using two complementary benchmarks for defining reference correlation outcomes. For each simulated dataset, the inferred correlation result produced by a given method was compared with the corresponding benchmark outcome, and method performance was summarized across the 1,000 replicates for each scenario–gradient combination.

Depending on the benchmark definition, performance of methods was quantified using appropriate aggregate metrics, including overall correctness and error-based measures.

Detailed definitions of the benchmarks and the corresponding evaluation metrics are provided in the following sections.

### Felsenstein’s Worst-Case Scenario: Deep-root Directional Shifts

Following previous simulation studies of directional trait shifts (Uyeda et al. 2018), we simulated abrupt shifts occurring on a single deep branch near the root of the phylogeny, representing a classical worst-case scenario for PIC-based methods. In this setting, trait evolution follows a BM process on all other branches, but an abrupt directional change is introduced on one of the branches closest to the root, a configuration known to generate clusters of extreme contrasts and to challenge standard phylogenetic comparative approaches.

Trait evolution under BM was simulated with unit evolutionary rate (σ^2^ = 1). Directional shifts were implemented as multiplicative scaling factors applied to trait values rather than additive offsets. Shift magnitudes ranged from 4^−3^ to 4^7^, where a magnitude of 4^0^ = 1 corresponds to the baseline condition with no effective shift. Positive exponents indicate trait upsizing, whereas negative exponents indicate downsizing. Each shift magnitude was simulated 1,000 times. Random seeds were controlled to ensure reproducibility and, where appropriate, independence between traits.

To implement the deep-root shift, the phylogenetic tree was split at the root into two subtrees—a background subtree and a shift subtree—using the extract.clade() function in the **ape** package (version 5.8) (Paradis and Schliep 2019). The background subtree was assigned a baseline root value of 1, whereas the shift subtree was assigned a root value scaled by the specified shift magnitude (1 ×shift magnitude). Trait evolution was then simulated independently on each subtree under BM using the rTraitCont() function from the **ape** package.

#### Two-trait shared shift (independent traits)

To evaluate false positive behavior under shared directional shifts, we simulated two independent traits (*X*_1_ and *X*_2_) on the same phylogeny within a regression framework, *X*_2_ = *βX*_1_ + *ε*, with *β* = 0. Under this setting, *ε* itself constitutes the second trait *X*_2_. Both traits evolved independently under a Brownian motion model with identical variance (σ^2^ = 1) and the same root value (1), and were therefore uncorrelated by construction. An abrupt shift of equal magnitude was introduced on the same deep branch in both traits, while the remainder of the tree followed background BM evolution. Independent random seeds were used for *X*_1_ and *X*_2_ to ensure the absence of any true association. This design allowed us to assess whether phylogenetic methods spuriously detect correlations when none exist.

#### Single-trait shift in the predictor (correlated traits)

To evaluate false negative behavior, we simulated two positively correlated traits with *X*_2_ = *βX*_1_ + *ε*, where *β* = 1 and *ε* followed a normal distribution with mean 0 and variance 0.01, following the phylogenetic regression framework of Revell (2010) and Mazel et al. (2016). An abrupt directional shift was introduced only in the predictor trait *X*_1_ on the same deep branch using the same set of shift magnitudes as described above, while *X*_2_ remained unchanged. To preserve the predefined correlation structure, *X*_1_ was first simulated across the entire tree, *X*_2_ was constructed as a function of *X*_1_, and *X*_1_ was then re-simulated only within the shift subtree using the same random seed, with the only change being that the subtree root value was shifted by the specified magnitude (i.e., the shift magnitude was added at the subtree root). The final *X*_1_ was assembled by combining background and shift-subtree values, whereas *X*_2_ remained unchanged. This design allowed us to quantify how often true correlations were masked by directional shifts affecting only one trait.

### Simulation with Randomly Located Trait Shifts

To evaluate method performance under more biologically realistic conditions, we extended the simulation framework by allowing the branch experiencing a directional shift to be randomly selected across the phylogeny. In contrast to shifts fixed at deep branches, this design captures scenarios in which shifts arise on shallow internal branches or terminal lineages.

For each replicate, a single branch was sampled uniformly at random from the phylogeny and assigned a directional shift. Except for the location of the shift, all simulation settings—including trait evolutionary models, shift implementation, shift magnitudes, and replication scheme—were identical to those used in the deep-root simulations described above.

As in the deep-root scenario, we considered both two-trait shared shifts in independent traits (to assess false positive behavior) and single-trait shifts in the predictor for correlated traits (to assess false negative behavior), with shifts applied to the randomly selected branch following the same procedures outlined above.

### Trait Shifts Introduced at Different Phylogenetic Depths

To evaluate whether method performance depends on the phylogenetic depth at which a directional shift occurs, we conducted an additional set of simulations on the 128-species fixed-balanced tree, in which shifts were introduced systematically at predefined phylogenetic depths. In contrast to the worst-case and random-position-shift simulations, this analysis was designed as a controlled diagnostic experiment to isolate the effect of shift depth while holding all other factors constant. Based on preliminary results from the deep-root (worst-case) simulations (see Results), which indicated that all methods are largely insensitive to downsizing shifts, we focused exclusively on a representative large-magnitude upsizing shift that poses the greatest challenge for phylogenetic correlation inference.

Specifically, we examined a shift magnitude of 4^5^, which provides a strong but non-extreme perturbation suitable for isolating depth-dependent effects. For each phylogenetic depth, a single shift was introduced on one randomly selected branch at that depth. At each shift depth, simulations were conducted under both the dual-trait shared-shift and the single-trait (*X*_1_-only) shift scenarios, with 1,000 independent replicates for each scenario.

### Simulated Traits Without Shifts: Fixed Correlation with Varying Noise

Although the primary focus of this study is on evaluating PIC-O(D)GC under shift-driven evolutionary scenarios, we also considered trait simulations without abrupt shifts to assess whether the workflow behaves comparably to established methods under gradual evolutionary dynamics. To provide such a baseline, we followed the core simulation framework used by Revell (2010) and Mazel et al. (2016), in which paired traits *X*_1_ and *X*_2_ were generated under a linear generative model,

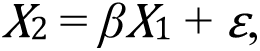

with *β* = 1, such that a structural dependence between the two traits is biologically encoded. The noise term *ε* was assigned varying variance to modulate the strength of the realized association. Under this design, these simulations primarily function as power analyses, assessing how effectively different methods recover an intended association as stochastic variation increases, while acknowledging that sufficiently large noise can render some replicates statistically indistinguishable from uncorrelated data.

To reflect different sources of phylogenetic signal under gradual evolutionary dynamics (i.e., without abrupt shifts), we considered three classes of simulation scenarios that differ in whether phylogenetic structure arises from the focal trait *X*_1_, from the noise term *ε*, or from both.

In the BM–BM scenario, both *X*_1_ and *ε* evolve under BM, with *X*_1_ having a variance of 1 and *ε* having variance *d*. Accordingly, both the signal and noise components carry phylogenetic structure, and the realized association between *X*_1_ and *X*_2_ can vary substantially across replicates as noise variance increases.

In the BM–Norm scenario, *X*_1_ evolves under BM with variance 1, whereas *ε* is drawn from an independent and identically distributed normal distribution with mean 0 and variance *d*. Because the noise term is phylogenetically unstructured, this scenario satisfies the assumptions of ordinary least squares (OLS) regression and therefore represents the only setting in which OLS is theoretically appropriate and included for comparison (Revell 2010).

In the Norm–BM scenario, *X*_1_ follows a normal distribution with mean 0 and variance 1, while *ε* evolves under BM with variance *d*. Here, phylogenetic signal arises solely from the noise term, allowing us to examine how phylogenetically structured error can obscure an otherwise simple linear relationship.

Across all three scenarios, the noise variance parameter *d* took values of 4^-3^, 4^-2^, 4^-1^, 4^0^, 4^1^, 4^2^, 4^3^, 4^4^, 4^5^, 4^6^, and 4^7^. For each scenario and value of *d*, we conducted 1,000 replicate simulations, resulting in a total of 33,000 datasets per phylogenetic tree.

We did not include the fourth scenario described by Revell (2010), in which both *X*_1_ and *ε* lack phylogenetic structure. Because this study focuses on evaluating method behavior in settings where phylogenetic signal is present in either the traits or the residual variation, explicitly simulating a fully non-phylogenetic scenario was not necessary for our purposes.

### Empirical Datasets

To complement the simulation analyses and evaluate the relative performance of PIC-O(D)GC against other phylogenetic comparative methods under real biological conditions, we applied these methods to three empirical datasets that may contain abrupt evolutionary shifts. Two of them were previously analyzed by Adams et al. (2024) in a study of phylogenetically robust regression.

The first dataset was obtained from the supplementary material of Makino and Kawata (2019), and includes propagule size, genome size, and a phylogenetic tree for 44 invertebrate species. Following the original study, both traits were log_10_-transformed prior to analysis to reduce skewness and improve normality assumptions. Propagule size in this context refers to the physical size (cm) of a reproductive unit that is separated from the parent and capable of developing into a new individual. In ecological and evolutionary studies, propagule size is widely used as a key indicator of an organism’s reproductive strategy, particularly along the r/K selection continuum (Romiguier et al. 2014). The phylogenetic tree provided by the authors was used without modification.

The second dataset was originally compiled by Scales et al. (2009) and includes the relative proportions of fast glycolytic (FG) and slow oxidative (SO) muscle fibers in the limb muscles of 22 lizard species. This dataset was later used by Uyeda et al. (2018) to illustrate the limitations of phylogenetic comparative methods, particularly in cases involving abrupt shifts or convergent trait evolution. We used the phylogenetic tree provided by Uyeda et al. (2018).

The third dataset includes body mass and population density for 109 species of Artiodactyla. Trait data were obtained from the PanTHERIA database (accessed June 4, 2025) (Jones et al. 2009). Species with missing values for either trait were excluded. Both body mass (grams) and population density (individuals per km^2^) were log_10_-transformed prior to analysis to reduce skewness and better meet the assumptions of linear models and comparative methods. The phylogenetic tree used for the Artiodactyla clade was extracted from the time-scaled maximum clade credibility tree for mammals developed by Upham et al. (2019), available at VertLife.org (accessed on June 4, 2025).

### Statistical Methods Compared

To evaluate the performance of PIC-O(D)GC examining the evolutionary relationship between *X*_1_ and *X*_2_ under both gradual evolution and scenarios involving abrupt evolutionary shifts, we compared it with several previously established phylogenetic comparative methods:

**1. PIC-Pearson**: Pearson correlation applied to PIC values; because it yields the same direction and significance inferences as an OLS regression on the same PICs, running a separate PIC-OLS analysis would be redundant, so we used only PIC-Pearson.
**2. PGLS**: We fitted five alternative evolutionary covariance models: BM (Felsenstein 1985), Pagel’s λ (Pagel 1999), OU fixed, OU random (Hansen 1997), and EB (Harmon et al. 2010). For each simulated dataset, we used the Akaike information criterion (AIC) to identify the best-fitting covariance structure. PGLS results were then reported under the model with the lowest AIC.
**3. Robust phylogenetic regression with both PIC and PGLS frameworks:** For each framework, we utilized four robust estimators: L1, S, M, and MM. These were implemented as PIC-L1, PIC-S, PIC-M, and PIC-MM for PIC-based methods, and PGLS-L1, PGLS-S, PGLS-M, and PGLS-MM for PGLS-based methods (Adams et al. 2024).
**4. Corphylo**: A method that calculates Pearson correlation coefficients for multiple continuous traits, allowing users to incorporate phylogenetic signal and measurement errors into the model (Zheng et al. 2009). This method assumes an OU process for trait evolution, making it particularly suitable for traits that deviate from the BM model. Corphylo serves a dual purpose: it enables both the detection of phylogenetic signals in individual traits and the examination of correlations between multiple traits. A key feature of Corphylo is its ability to include measurement errors of trait data, potentially improving accuracy under real-world conditions.
**5. MR-PMM**: Multi-response phylogenetic mixed models (MR-PMM) extend phylogenetic regression frameworks by jointly modeling multiple traits as responses, allowing the decomposition of trait covariation into phylogenetic and residual components (Westoby et al. 2023, Halliwell et al. 2025). Unlike conventional phylogenetic regression, MR-PMM does not impose a directional predictor–response structure and is therefore well suited for assessing trait correlations and co-evolution. Although our simulations involve only two traits rather than higher-dimensional response vectors, MR-PMM is directly applicable to this bivariate setting and represents a conceptually distinct framework for evaluating phylogenetic trait associations.
**6. Raw-Pearson**: The standard Pearson correlation applied directly to the original trait values, without any phylogenetic correction. Because Pearson (or its OLS equivalent) assumes i.i.d. residuals, we report Raw-Pearson results only for the BM–Norm scenarios—the only conditions in our simulations that satisfy this assumption. Although Pearson/OLS methods are seldom used in phylogenetic comparative analyses due to the non-independence created by shared ancestry, they remain a useful theoretical benchmark when their statistical requirements are met (Revell 2010).

### Outlier-and Distribution-Guided Phylogenetic Correlation Analysis

To address the influence of evolutionary shifts and outliers in phylogenetic comparative analysis, we introduced a novel workflow: PIC-O(D)GC. In this method, the choice between Pearson and Spearman correlation analysis is guided by the identification of outliers within the PIC data, with additional consideration of the PIC value distribution in small-sample settings.

Potential outliers in PICs were identified using the boxplot rule (Tukey 1977), flagging values beyond Q1 − 1.5 × IQR or Q3 + 1.5 × IQR, where Q1 and Q3 are the first and third quartiles and IQR = Q3 – Q1 is the interquartile range. This nonparametric, distribution-free rule—corresponding to the whiskers of the boxplot—provides a robust and widely used definition of extreme observations, flagging about 0.7% of normally distributed data as outliers (Hoaglin et al. 1986). Because the boxplot rule is intentionally conservative, it may occasionally flag points spuriously due to random variation or over-flag observations in skewed distributions. However, such misclassification has little practical impact in our framework, as flagged datasets are analyzed using Spearman correlation, for which a small number of additional flagged points primarily leads to a modest loss of power while preserving monotonic associations. In contrast, failing to control even a single influential outlier in a Pearson analysis can substantially distort, or even reverse, the estimated correlation.

Outlier detection is performed at the level of individual observations: if a given sample is flagged as an outlier in any trait, that observation is treated as an outlier overall, and the corresponding dataset is analyzed using Spearman correlation; otherwise, Pearson correlation is applied. This rule ensures robustness when one or both traits—whether at the same or different phylogenetic positions—experience abrupt shifts, and, although not explored here, the underlying logic can be readily generalized to multivariate settings, including partial correlation analyses.

For larger datasets with 128 species (127 PIC values), Pearson correlation, despite its reliance on normality assumptions, is generally robust to minor deviations from normality due to the central limit theorem. In such cases, the impact of skewness and kurtosis is reduced by the large sample size, ensuring reliable correlation estimates. Therefore, for these datasets, the choice of Pearson or Spearman correlation was determined solely by the presence or absence of outliers.

For smaller datasets with 16 species (15 PIC values), both normality testing and outlier detection were conducted. This workflow, referred to as PIC-ODGC, dynamically selects the appropriate correlation method based on data characteristics. If no outliers were detected and normality assumptions were satisfied, the Pearson correlation was used. In cases where outliers were present, or normality was violated, Spearman correlation was applied to minimize the impact of extreme values on the results.

### Benchmark I: Structural-Coupling–Based Evaluation (β-based)

In simulation-based evaluations, trait associations are often benchmarked against the structure explicitly specified in the generative model (*X*_2_ = *βX*_1_ + *ε*). Under this approach, correlations are treated as present or absent according to whether the coupling parameter (*β*) is encoded during data generation.

Accordingly, we implemented a generative-model–based benchmark in which simulation replicates with a coupling parameter *β* = 1 were classified as true positives, whereas replicates with *β* = 0 were classified as true negatives. For each replicate, method performance was evaluated based on whether a statistically significant association between the two traits was detected.

This benchmark directly assesses a method’s ability to recover the structural relationship encoded by the coupling parameter *β* in the simulation design and therefore provides a clear reference for evaluating consistency with that aspect of the generative model. Performance was summarized using class-specific error rates, including the false positive rate (FPR), defined as the proportion of *β* = 0 replicates incorrectly classified as significant, and the false negative rate (FNR), defined as the proportion of *β* = 1 replicates in which the encoded relationship was not detected.

### Benchmark II: Data-Pattern–Based Evaluation

Beyond assumptions encoded in the generative model about trait coupling, a distinct inferential question concerns whether an association specified during data generation is statistically supported by the realized data. In stochastic evolutionary processes, the realized relationship between traits is jointly determined by the coupling parameter *β* and evolutionary noise *ε*, such that even strong generative coupling may yield weak or undetectable associations when noise variance is large (see Supplementary Methods S1). In empirical data, *β* and *ε* are not separately observable, and inference is therefore based on their combined effect on realized trait patterns.

To address this distinction, we implemented an additional benchmarking framework that evaluates whether association is statistically supported in the realized data. Under this framework, benchmark labels were assigned at the replicate level according to whether evidence for trait association was present in the realized data.

Specifically, for each simulation replicate, evidence for association was evaluated using correlation tests applied to normalized branch-level trait changes. Because ancestral states are recorded in the simulations, branch-level changes can be computed directly as Δ*X*_1_/*L* and Δ*X*_2_/*L*, where Δ*X* denotes trait change along each branch and *L* the corresponding branch length. Under a Brownian motion evolutionary model, these normalized changes are independently distributed across branches, providing a valid basis for conventional correlation testing (Supplementary Methods S1). Replicates exhibiting significant correlation (*p* < 0.05) were classified as benchmark-positive, whereas replicates lacking significant evidence were classified as benchmark-negative. Method performance was quantified using classification accuracy, defined as the proportion of replicates for which inferred results agreed with these data-driven benchmark labels.

This benchmark therefore targets a different inferential question than the *β*-based framework. Whereas the *β*-based benchmark evaluates method performance relative to the structural coupling specified during data generation, the data-pattern–based benchmark assesses whether the combined effects of *β*, *ε*, and finite sample size yield statistically supported association in the realized data (Ives 2022).

### Programs and Packages

All simulations and analyses were performed in the R environment (version 4.1.3) (R Core Team 2020) on a Linux system. The **phytools** package (version 2.3.0) (Revell 2024) and **ape** package (version 5.8) (Paradis and Schliep 2019) were used for tree generation. Simulations and PIC calculations were performed using the **ape** package in R. PIC values were calculated with ape∷pic() under a Brownian motion assumption, using the standard contrast-based pruning procedure rather than global ancestral-state reconstruction. Normality tests were conducted using the Shapiro-Wilk test (Royston 1992) via the shapiro.test() function from the stats package (version 4.3.1). Pearson and Spearman correlation analyses were performed using stats∷cor.test(). All PGLS regressions were conducted using the phylolm() function from the phylolm package (version 2.6.2) (Ho and Ane 2014). Corphylo was conducted using ape∷corphylo() function (Zheng et al. 2009). MR-PMM was conducted using the MCMCglmm() function from the MCMCglmm package (version 2.36) (Sorensen and Gianola 2002, Hadfield 2010, Hadfield and Nakagawa 2010). Robust phylogenetic regression were conducted using the Conduct.Robust_PhylogeneticRegression() function from ROBRT package (version 0.0.0.9000) (Adams et al. 2024).

## Results

### Impact of Abrupt Evolutionary Shifts on Data Patterns and Correlation Analysis

To illustrate PIC-ODGC’s ability to handle abrupt evolutionary shifts, we simulated two traits on a fixed-balanced phylogenetic tree with 16 species (Fig. 1a), considering two shift configurations: (1) simultaneous shifts applied to both traits, which are uncorrelated, and (2) shifts applied to *X*_1_ only, where *X*_1_ and *X*_2_ are correlated. For each configuration, we selected three shift levels—baseline, a weak shift, and a strong shift—as representative cases to illustrate how shift magnitude affects trait distributions (e.g., group separation) and contrasts (e.g., emergence of isolated outliers).

**Figure 1.**
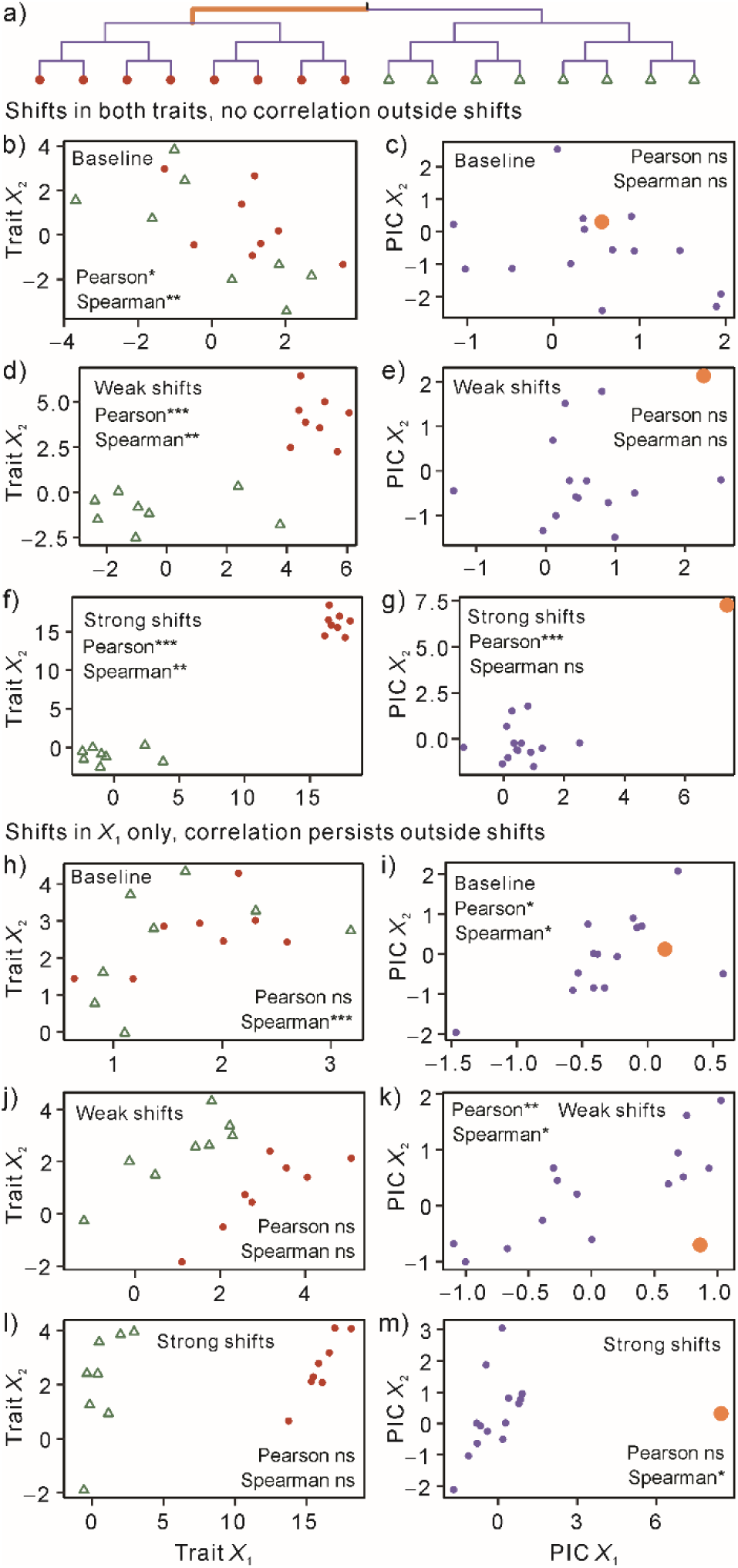
Effects of abrupt shifts on data pattern and correlation analysis. a) Schematic fixed-balanced phylogenetic tree with 16 species; the thick branch marks the location of an abrupt evolutionary shift. b–c) Baseline scenario for the dual-shift simulations shown in panels d–g. In this scenario, both traits *X*_1_ and *X*_2_ evolve independently under Brownian motion (BM) without an effective shift (baseline condition; multiplicative shift factor = 1). d–e) and f–g) Dual-shift simulations in which both traits undergo an abrupt shift on the marked branch, with increasing shift magnitudes (4-fold and 16-fold, respectively). h–i) Baseline scenario for the single-trait shift simulation series shown in panels j–m. In this scenario, *X*_1_ evolves under BM without an effective shift (multiplicative shift factor = 1), and *X*_2_ is generated from *X*_1_ via a fixed linear relationship with added noise. j–k) Moderate (4-fold) shift in *X*_1_ only. l–m) Strong (64-fold) shift in *X*_1_ only. In all phylogenetically independent contrast (PIC) plots (c, e, g, i, k, m), the contrast corresponding to the branch marked by the thick line in panel a is highlighted as an enlarged dot. In panels e, g, k, and m, this contrast reflects an evolutionary shift; in panels c and i, it serves as a baseline reference. In the weak dual-shift dataset (e), the highlighted contrast lies approximately 1.8 interquartile ranges (IQRs) above the third quartile (Q3) for PIC *X*_1_ and 2.1 IQRs above Q3 for PIC *X*_2_. In the strong dual-shift dataset (g), this deviation increases markedly, reaching approximately 8.8 and 7.8 IQRs above Q3 for PIC *X*_1_ and PIC *X*_2_, respectively. In the weak single-trait shift dataset (k), the shifted trait shows only moderate deviation in PIC, remaining below the outlier threshold (i.e., < 1.5 IQRs above Q3). In contrast, in the extreme single-trait shift case (m), PIC *X*_1_ exhibits a pronounced deviation of approximately 5.9 IQRs above Q3. Pearson and Spearman correlation tests: *** indicates *p* < 0.001; ** indicates *p* < 0.01; * indicates *p* < 0.05; ns indicates not significant (*p* ≥ 0.05). Data used to generate this figure are available in Supplementary Table S1.

As shown in Figures 1d and 1f, when both traits undergo an abrupt shift near the root, the raw trait values form two distinct clusters in the coordinate space, with the separation between these clusters increasing as the shift magnitude becomes larger. In the PIC coordinate system, the shared shift manifests as a single outlier in the PIC data, with the degree of outlier deviation becoming more pronounced as the shift magnitude increases (Figs. 1e and 1g).

When phylogenetic autocorrelation is ignored, directly applying Pearson or Spearman correlation to the raw trait data frequently results in false positives, regardless of the shift magnitude (Figs. 1b, 1d, and 1f). In contrast, controlling for phylogenetic autocorrelation by using PIC values eliminates false positives when there is no effective shift or when the shift is small (Figs. 1c and 1e). However, under large upsizing shifts, Pearson correlation becomes highly sensitive to the outlier, leading to a false positive result and erroneously inferring a significant positive correlation between *X*_1_ and *X*_2_, akin to non-phylogenetic methods.

Conversely, the Spearman correlation applied to the PIC data under large shifts avoids false positives by correctly detecting no significant correlation between the two traits (Fig. 1g).

In the second set of simulations, where only *X*_1_ experiences an abrupt shift while *X*_2_ follows unperturbed BM, the true correlation between the two traits remains intact outside the shifted lineage. As in the dual-shift scenario, the raw data exhibit a visible cluster separation as the shift magnitude increases (Figs. 1j and 1l). Despite the presence of a shift in only one trait, the PIC data again display a prominent outlier under large upsizing (Fig. 1m). While Pearson correlation applied to the PIC values is affected by this outlier and fails to detect the existing correlation—resulting in a false negative (Fig. 1m)—Spearman correlation remains robust across shift levels, consistently recovering the true positive association (Figs. 1i, 1k, and 1m).

Together, these results show that large upsizing shifts can produce outliers in PIC data, which in turn may lead to false negatives or false positives in Pearson-based correlation analyses, depending on whether the shift affects one or both traits. While we simulated a deep-branch shift to illustrate maximal distortion, equally large shifts on intermediate or terminal branches can also generate extreme contrasts and thereby mislead mislead PIC-based parametric analyses. This reflects an inherent limitation of traditional PIC approaches, which weigh all contrasts equally without regard to their evolutionary context.

### Refining Robust Regression Methods for Comparison with PIC-O(D)GC

To avoid visual and interpretive complexity, we restricted comparisons to representative implementations within each methodological category rather than exhaustively presenting all possible variants. Specifically, we evaluated robust regression methods spanning two methodological types—PGLS-and PIC-based regression—combined with four robust estimators (L1, S, M, and MM), and selected representative implementations based on objective performance criteria derived from simulation results.

Under shift conditions, PIC-MM consistently ranked as the best or among the best-performing robust estimators (Supplementary Result S1). In no-shift simulations, PIC-MM continued to rank among the stronger-performing methods, although performance differences across estimators were generally modest. These conclusions were consistent under two benchmark criteria used to evaluate robust regression, suggesting the observed ranking is not sensitive to a particular evaluation standard. In line with recent results reported by Adams et al. (2024), we therefore retain PIC-MM as the primary robust comparator to PIC-O(D)GC.

### PIC-OGC: Performance Across Evolutionary Shifts and No-Shift Regimes

We first compared the performance of PIC-OGC with five alternative methods using simulations with evolutionary shifts on the 128-species fixed-balanced tree, focusing initially on scenarios with simultaneous shifts in both traits.

Under Felsenstein’s worst-case scenario with deep-root directional shifts (Fig. 2a), PIC-OGC exhibited the strongest control of type I error, maintaining a false positive rate of zero across all shift magnitudes examined. PIC-MM was similarly insensitive to increasing shift magnitude, although its false positive rate was slightly higher than that of PIC-OGC, remaining close to 0.05 across the entire gradient. Other methods—including PIC-Pearson, which serves as the most direct comparator to PIC-OGC—also showed low false positive rates under weak shifts. As shift magnitude increased, false positive rates for these methods rose sharply, and at shift magnitudes ≥ 4^3^, all approached or reached one, indicating near-complete loss of type I error control.

**Figure 2.**
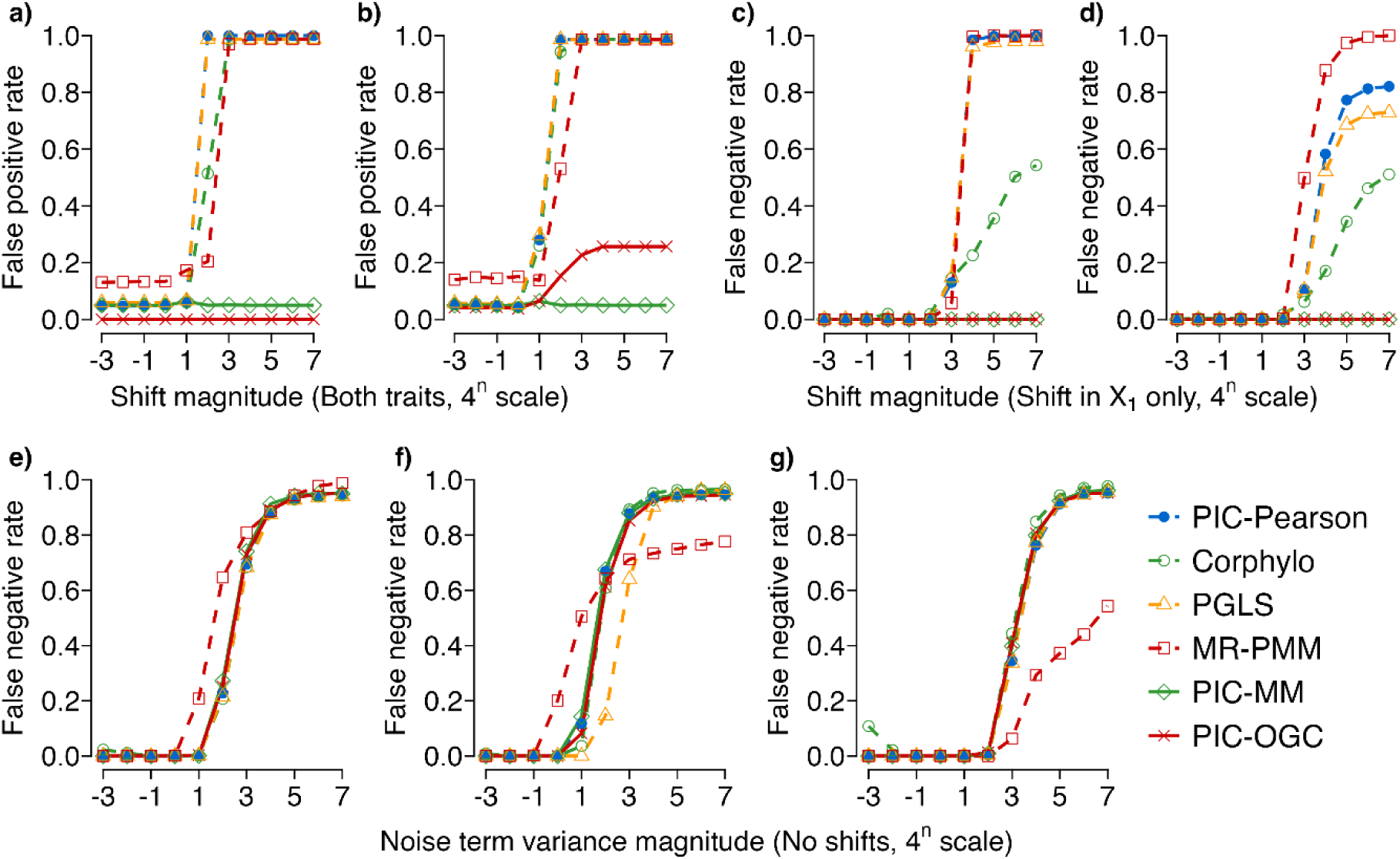
Performance of six methods across evolutionary shift and no-shift regimes simulated on a 128-species fixed-balanced tree. Panels a–b report false positive rates under simultaneous shifts in both *X*_1_ and *X*_2_, and panels c–d report false negative rates when only *X*_1_ is shifted. Panels a and c correspond to Felsenstein’s worst-case scenario, in which directional shifts are introduced on deep-root branches, whereas panels b and d show results from simulations with randomly located trait shifts. Panels e–g show performance under gradual evolutionary scenarios without abrupt shifts, in which traits follow the relationship *X*_2_ = *X*_1_ + *ε* with increasing noise variance. Three evolutionary background conditions are considered: BM–BM (panel e), BM–Norm (panel f), and Norm–BM (panel g). Shift and noise variance magnitudes are expressed on a 4^n^ scale, where the horizontal axis shows only the exponent (*n*) rather than the full value. Additional details on the simulation design, gradient configurations, and performance metrics are provided in the Materials and Methods. The data used to generate this figure are available in Supplementary Table S2.

When trait shifts were introduced at randomly selected locations on the tree (Fig. 2b), PIC-OGC no longer ranked as the best-performing method across the full shift gradient. Under weak shifts, PIC-OGC, PIC-MM, and PIC-Pearson jointly exhibited the lowest false positive rates. However, as shift magnitude increased, PIC-MM consistently maintained the lowest false positive rate, whereas the false positive rate of PIC-OGC increased markedly, reaching a maximum of approximately 0.26. The other methods—including PIC-Pearson—showed rapidly escalating false positive rates with increasing shift magnitude, again approaching one under strong shifts.

We next examined scenarios with univariate shifts affecting only *X*_1_. Regardless of whether shifts occurred under the worst-case (Fig. 2c) or randomly located design (Fig. 2d), PIC-OGC and PIC-MM showed identical performance, maintaining false negative rates of zero across all shift magnitudes. The remaining four methods also achieved near-zero false negative rates under weak shifts, but diverged markedly as shift magnitude increased. Under strong shifts, Corphylo exhibited the lowest false negative rates among these alternatives, whereas the other methods showed substantially reduced sensitivity.

In addition to simulations with abrupt evolutionary shifts, we also examined method performance under gradual evolutionary scenarios without discrete shifts. These analyses were conducted on the 128-species fixed-balanced tree, with trait relationships defined as *X*_2_ = *X*_1_ + *ε*, where the noise term *ε* varied across 11 gradients of variance (Figs. 2e–g). Three evolutionary background scenarios were considered: BM–BM, BM–Norm, and Norm–BM.

Across all three no-shift scenarios, overall performance patterns differed markedly from those observed under shift conditions. In particular, MR-PMM exhibited a distinct and inconsistent behavior, with its performance relative to other methods fluctuating noticeably across noise levels and evolutionary settings, setting it apart from the other methods without a consistent advantage or disadvantage.

With the exception of PGLS under the BM–Norm scenario—where its performance was clearly degraded—the remaining five methods showed broadly similar behavior across noise gradients. Under gradual evolution, differences among these methods were modest, and no approach consistently emerged as clearly superior or inferior across scenarios. This convergence in performance contrasts with the pronounced method-specific differences observed under abrupt shifts, underscoring that evolutionary shifts, rather than background noise alone, are the primary drivers of method divergence.

In the BM–Norm scenario, where the data-generating process conforms the assumptions of standard linear regression, correlation analysis on raw trait values (Raw-Pearson, equivalent to OLS) was included as a theoretical benchmark (Supplementary Fig. S1). Under increasing noise levels, Raw-Pearson closely tracked the performance of PGLS, with both methods forming a consistently low-performing group. This pattern is consistent with theoretical expectations: when residuals are phylogenetically independent, PGLS effectively reduces to OLS (Revell 2010, Symonds and Blomberg 2014).

### Sensitivity of Methods to the Phylogenetic Location of Evolutionary Shifts

The above results shown in Figures 2b and 3d summarize method performance under randomly located trait shifts, effectively averaging behavior across all possible phylogenetic positions. To further characterize method sensitivity to where a shift occurs on the tree, we conducted an additional set of diagnostic simulations in which shifts were introduced systematically at each phylogenetic depth. Based on the separation among methods observed in Figure 2, we selected a representative shift magnitude of 4^5^, and applied shifts independently at each branch level of the 128-species fixed-balanced tree.

**Figure 3.**
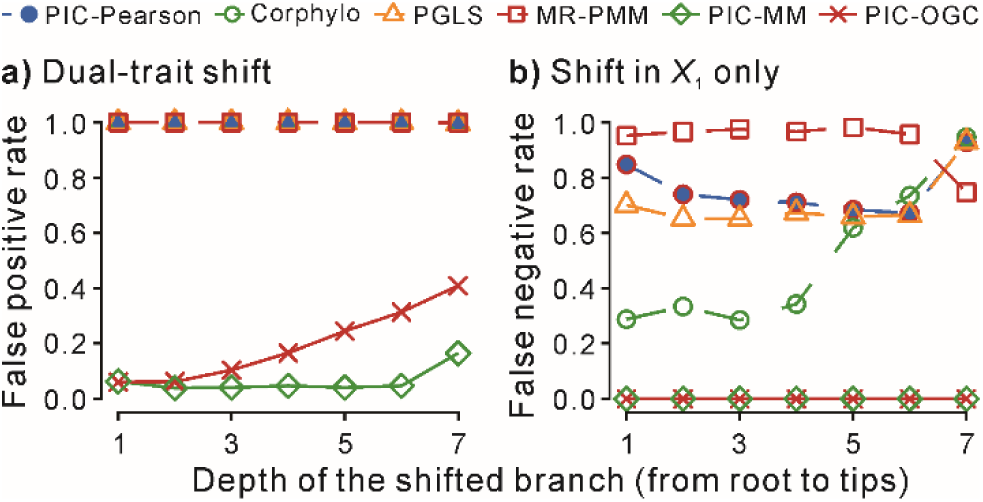
Sensitivity of method performance to the phylogenetic location of evolutionary shifts. Performance of six methods evaluated under shifts introduced at different depths of a 128-species fixed-balanced tree. A representative shift magnitude of 4^5^ was applied systematically at each phylogenetic depth. Panel a shows false positive rates under simultaneous shifts in both traits, whereas panel b shows false negative rates under univariate shifts affecting only *X*_1_. Additional details of the simulation design and performance metrics are provided in the Materials and Methods. The data used to generate this figure are available in Supplementary Table S3.

Under simultaneous shifts in both traits (Fig. 3a), both PIC-MM and PIC-OGC showed reduced robustness to shifts occurring near the tips compared to shifts on deeper branches closer to the root. This pattern indicates that terminal shifts pose a greater challenge for robust phylogenetic methods than deep-root shifts. Notably, PIC-OGC exhibited stronger sensitivity to shift location than PIC-MM, with false positive rates increasing more rapidly toward the tips.

In contrast, under univariate shifts affecting only *X*_1_ (Fig. 3b), PIC-MM and PIC-OGC consistently maintained false negative rates of zero across all branch levels, indicating stable power regardless of shift location. Interestingly, the remaining four methods showed pronounced sensitivity to shift position under univariate shifts, with false negative rates varying substantially across branch levels.

### Performance of the Methods Under Non-BM Evolutionary Conditions

While the application of OLS regression or Pearson correlation on PIC values requires the assumption that traits evolve under a BM model—so that PICs are statistically i.i.d.—the computation of PIC values themselves does not strictly rely on BM. Instead, PIC values are defined as algebraic contrasts on a phylogeny, and remain meaningful under a wide range of evolutionary models (Garland et al. 1992). It is the statistical interpretation (e.g., *p*-values) of PIC-OLS or PIC-Pearson that breaks down when the BM assumption is violated. In contrast, methods such as PIC-MM and PIC-OGC—though still operating on PIC values—employ robust estimators or nonparametric correlations that do not require the strict assumptions of BM. Thus, they are retained in our comparison alongside PGLS, MR-PMM, and Corphylo when evaluating method performance under simulated non-BM conditions. Importantly, this also means that their performance can be fairly compared to methods that are explicitly designed to accommodate a range of evolutionary models.

Although all simulations incorporated Brownian-motion–structured components, the realized tip-level trait patterns may deviate from ideal BM expectations due to evolutionary shifts, added stochastic noise, or their interaction. As a result, these data can sometimes be better approximated by alternative covariance structures (e.g., OU, λ, or EB) when fitted using PGLS. To capture this distinction, we used AIC-based model comparison as a diagnostic criterion to classify simulated datasets as either BM-consistent or non-BM-consistent at the tip level under standard PGLS covariance models.

Overall, performance patterns under non-BM conditions (Supplementary Fig. S2) closely resembled those observed across the full set of simulations (Fig. 2). With the exception of specific edge cases in which non-BM datasets are not defined (Supplementary Fig. S2a, c, d) and direct comparisons are therefore not possible, relative method behavior remained largely unchanged.

Across scenarios involving evolutionary shifts, including both bivariate shifts and randomly located univariate shifts, PIC-OGC and PIC-MM continued to show strong and stable performance under non-BM conditions, maintaining low error rates across a wide range of shift magnitudes (Supplementary Figs. S2a–d). As observed across the full set of simulations, under randomly located single-trait shifts at large magnitudes, PIC-OGC exhibited noticeably higher error rates than PIC-MM. Importantly, this deviation was limited in scope and did not alter the overall rank pattern of relative method performance.

In contrast, under gradual evolutionary scenarios without abrupt shifts, differences among methods under non-BM conditions were modest, and PIC-OGC and PIC-MM were not clearly distinguishable from the other approaches (Supplementary Figs. S2e–g). As observed across the full set of simulations, no method consistently emerged as either superior or inferior in the absence of shifts, indicating that background evolutionary heterogeneity alone is insufficient to drive substantial performance divergence.

### PIC-O(D)GC: Robust Performance across Tree Structure and Sample Size

To assess the robustness of the results on PIC-O(D)GC performance across evolutionary shifts and no-shift regimes, we repeated the analyses shown in Figure 2 on additional phylogenetic settings, including fixed-balanced trees with 16 and 256 species and randomly generated trees with 128 species. Across these additional tree topologies and sample sizes, PIC-O(D)GC and PIC-MM consistently ranked among the top-performing methods under evolutionary-shift scenarios (Supplementary Figs. S3–S5). Although the magnitude of performance differences varied with shift gradient, tree structure, and sample size—the slight advantage of PIC-O(D)GC over PIC-MM observed under the 128-species fixed-balanced tree (Fig. 2a) was no longer evident under other tree configurations—the overall relative ranking of methods remained unchanged.

### Secondary Benchmark Confirms Robustness of PIC-Based Methods

All results reported above are based on the *β*-based benchmark, which evaluates method performance relative to the generative coupling between traits. To provide a complementary perspective, we additionally evaluated all methods using a second, data-pattern–based benchmark. This alternative framework exploits the availability of ancestral states in simulations, enabling the computation of statistically independent branchwise changes (Δ*X*_1_/*L* and Δ*X*_2_/*L*) that can be evaluated using standard statistical methods (e.g., Pearson and Spearman correlations), rather than phylogenetic comparative methods.

When evaluated under the data-pattern–based benchmark, differences relative to the *β*-based benchmark are minimal across most simulation settings. In particular, under evolutionary-shift scenarios and across all three gradual-evolution scenarios (BM–BM, BM–Norm, and Norm–BM) at low noise levels, the two benchmarks yield largely mirrored conclusions: methods show low error rates when evaluated against the β-based benchmark and correspondingly high accuracy under the data-pattern–based benchmark. As noise variance increases under the tree gradual-evolution scenarios, however, the two benchmarks begin to diverge. False-negative rates under the *β*-based benchmark rise sharply, whereas accuracy under the data-pattern–based benchmark declines more gradually and remains substantially higher (Fig. 4).

**Figure 4.**
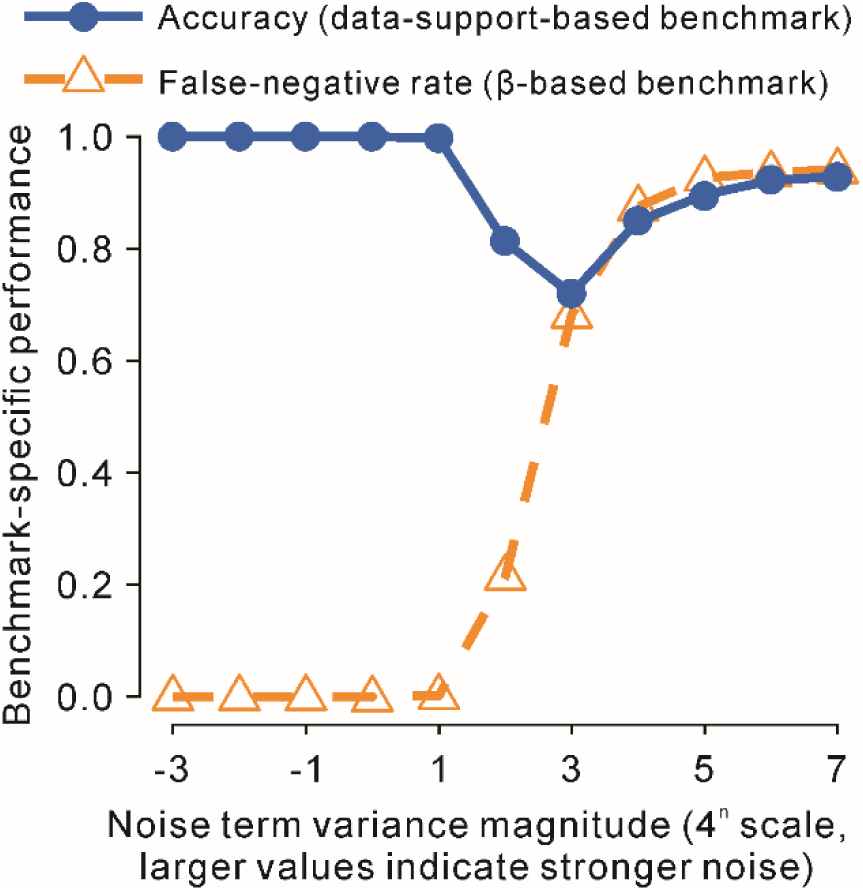
Illustrative comparison of benchmark outcomes for a representative method (PGLS) under a gradual-evolution simulation scenario (BM–BM), simulated on a fixed-balanced tree with 128 species. The figure contrasts evaluations under the *β*-based and data-pattern–based benchmarks and highlights their differing perspectives under noise-dominated conditions. Additional details of the simulation design and the two benchmarks are provided in the Materials and Methods. Detailed method-specific results under the data-pattern–based benchmark are reported in Supplementary Figs. S6–S12. The data used to generate this figure are available in Supplementary Table S4.

This divergence reflects differences in benchmark definition rather than changes in method behavior. Under extreme noise, structural coupling defined by *β* may persist in the generative model, while realized data provide little statistical support for association (see Supplementary Methods S1 for details). Methods that correctly infer absent support therefore remain accurate under the data-pattern–based benchmark even when *β* = 1.

Figure 4 provides a schematic illustration of this benchmark divergence using a representative method (PGLS), while detailed method-specific results under the data-pattern–based benchmark are reported in Supplementary Figs. S6–S12. Despite these benchmark-dependent differences, overall performance patterns under the data-pattern–based benchmark closely mirror those obtained under the *β*-based benchmark, with relative method rankings remaining largely unchanged and PIC-O(D)GC and PIC-MM continuing to rank among the top-performing methods under evolutionary-shift scenarios.

### Application of PIC-O(D)GC to Selected Empirical Datasets with Trait Shifts

Simulation results, validated against internal benchmarks from the generative model or recorded ancestral states, showed PIC-O(D)GC and PIC-MM to be most robust under abrupt shifts. To evaluate PIC-O(D)GC’s performance on real-world data, we analyzed three empirical trait datasets, derived either in full or in part from published phylogenetic resources or comparative studies (Jones et al. 2009, Scales et al. 2009, Makino and Kawata 2019).

#### Invertebrate propagule size vs. genome size

Across 44 invertebrate species, the PIC distribution revealed seven extreme contrasts (Fig. 6a). Notably, log₁₀-transformation of the original trait values did not eliminate these deviations, indicating that standard variance-stabilizing transformations are insufficient to remove lineage-specific extremes after phylogenetic correction. These contrasts likely reflect heterogeneous evolutionary shifts occurring along a small number of branches and illustrate how a few extreme observations can exert disproportionate influence on correlation-based comparative analyses (Fig. 5b). In this dataset, methods that explicitly account for extreme contrasts—such as PIC-OGC and PIC-MM—returned non-significant associations, whereas other methods inferred significant correlations. This discrepancy highlights the sensitivity of comparative correlation results to the treatment of extreme contrasts and underscores the importance of explicitly addressing such deviations in phylogenetically informed analyses.

**Figure 5.**
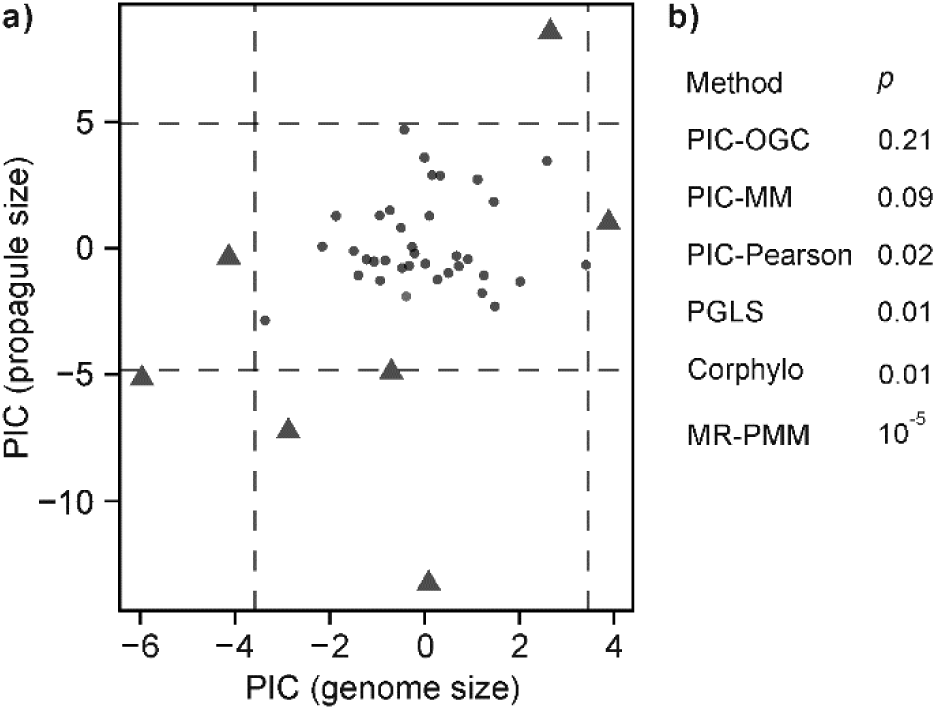
Relationship between propagule size and genome size across 44 invertebrates. a) Phylogenetically independent contrasts (PICs) between log_10_-transformed propagule size and genome size. Dashed lines indicate the outlier boundaries defined by the boxplot rule, Q1 − 1.5 × IQR and Q3 + 1.5 × IQR (Tukey 1977). Contrasts exceeding these thresholds in either trait are marked as filled triangles and treated as outliers, while remaining contrasts are shown as points. b) Statistical significance of the association between genome size and propagule size as assessed by six comparative methods. Detailed correlation coefficients and *p*-values, along with the original trait values and the PIC values from which the scatterplots were derived, are provided in Supplementary Table S5.

**Figure 6.**
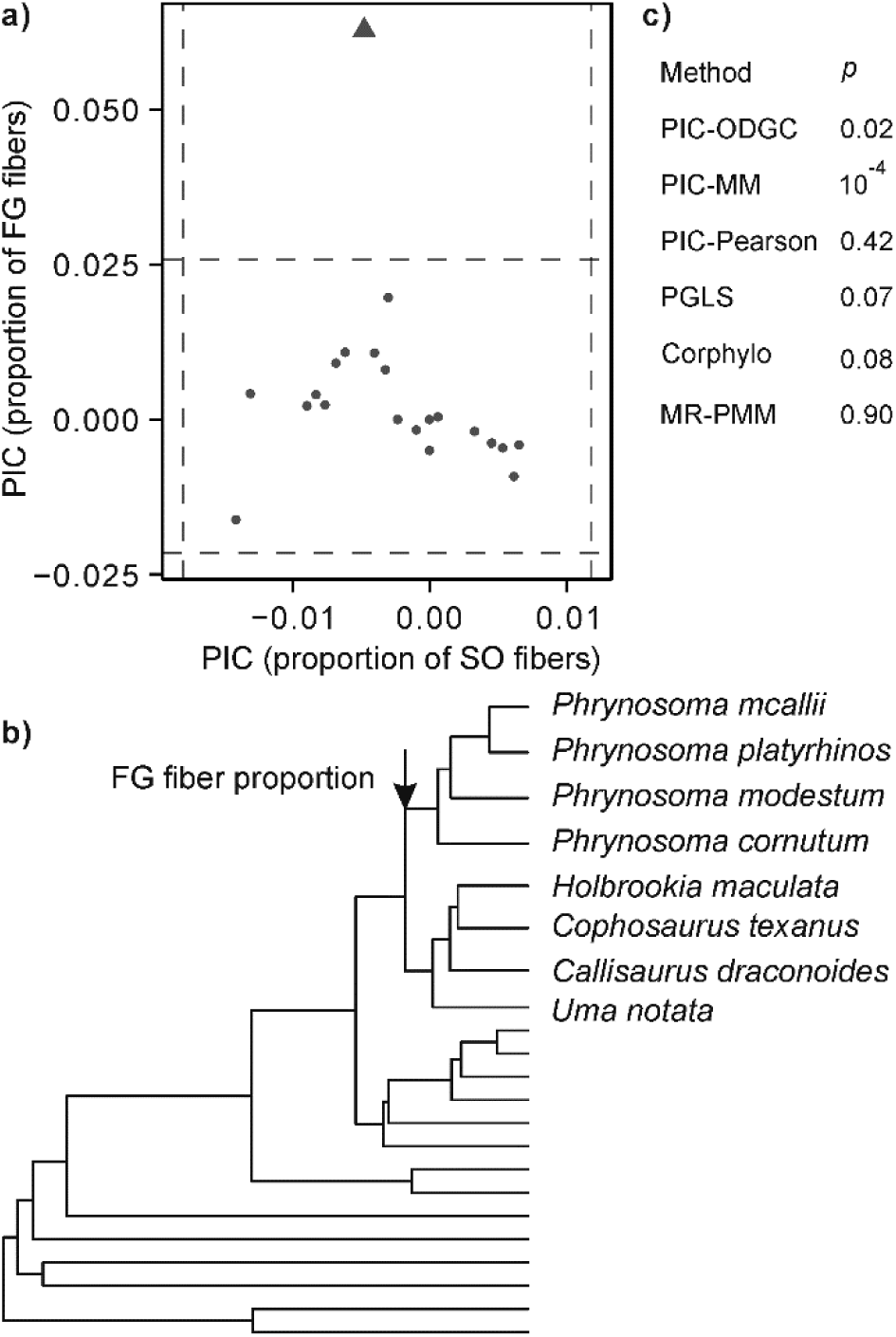
Relationship between slow oxidative (SO) and fast glycolytic (FG) muscle fiber proportions in 22 lizard species. a) Phylogenetically independent contrasts (PICs) between the proportions of SO and FG muscle fibers. Dashed lines indicate the outlier boundaries defined by the boxplot rule, Q1 − 1.5 × IQR and Q3 + 1.5 × IQR (Tukey 1977). A contrast exceeding these thresholds in FG fiber proportion is marked as filled triangles and treated as outliers, while remaining contrasts are shown as points. b) Phylogenetic tree highlighting the branch associated with the outlier contrast. The arrow marks the lineage corresponding to a pronounced reduction in FG fiber proportion. c) The *p*-values from multiple phylogenetic comparative methods testing for association between SO and FG fiber proportions. Detailed correlation coefficients and *p*-values, along with the original trait values and PIC values from which the scatterplots were derived, are provided in Supplementary Table S6.

#### Lizard fast glycolytic vs. slow oxidative fiber proportion

Across 22 lizard species, the PIC distribution revealed a single univariate outlier in FG fiber proportion (Fig. 6a), corresponding to a contrast between *Phrynosoma* and its sister clade (Fig. 6b). Inspection of the raw data revealed that this contrast is driven almost entirely by a marked reduction in FG fiber proportion across the four sampled *Phrynosoma* species, indicating a directional downsizing shift within this lineage. Across the full dataset, phylogenetic comparative methods produced mixed results (Fig. 6c). While PIC-ODGC and PIC-MM detected a statistically significant association between SO and FG fiber proportions, other methods such as PIC-Pearson, PGLS, and MR-PMM failed to do so.

To test the influence of the downsizing shift on methods’ performance, we re-ran all analyses after removing the four *Phrynosoma* species. Following this exclusion, all methods except MR-PMM returned *p*-values below 0.05, consistently indicating a negative association between SO and FG fiber proportions (Supplementary Table S6). This result suggests that the localized evolutionary shift within *Phrynosoma* both introduces statistical heterogeneity and suppresses correlation signals under certain comparative frameworks.

#### Body mass and population density in even-toed ungulates

The PIC distribution across 109 species exhibits seven univariate outliers with varying degrees of deviation (Fig. 7a), reflecting extreme evolutionary contrasts along specific branches. Despite all methods indicating a negative association between the two traits, their statistical support differs markedly (Fig. 7b). In particular, two methods sensitive to extreme values, PIC-Pearson and Corphylo, yield non-significant results, whereas approaches that explicitly account for outliers, including PIC-OGC and PIC-MM, detect a significant negative association.

**Figure 7.**
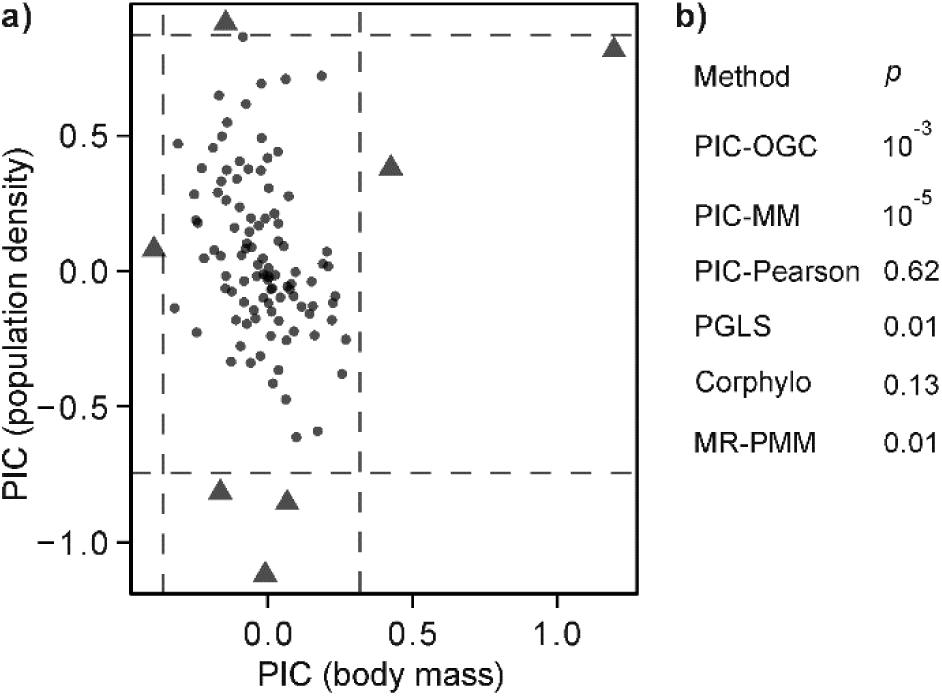
Relationship between body mass and population density in even-toed ungulates (Artiodactyla). a) Phylogenetically independent contrasts (PICs) between log_10_-transformed body mass and population density for 109 artiodactyl species. Dashed lines indicate the outlier boundaries defined by the boxplot rule, Q1 − 1.5 × IQR and Q3 + 1.5 ×IQR (Tukey 1977). Contrasts exceeding these thresholds in either trait are marked as filled triangles and treated as outliers, while remaining contrasts are shown as points. b) *p*-values from multiple comparative methods testing for an association between body mass and population density. Detailed correlation coefficients and *p*-values, along with the original trait values and PIC values from which the scatterplots were derived, are provided in Supplementary Table S7.

Because the direction of evolutionary change could not be confidently inferred from the phylogenetic positions of the most extreme contrast pairs, we adopted a conservative sensitivity analysis focusing on the two most pronounced univariate deviations in the PIC distribution (Fig. 7a). These two contrasts represent the most extreme deviations and are therefore the most likely to distort inference under methods that assume identically distributed contrasts; weaker departures were retained because their influence on overall results is expected to be negligible. Specifically, we removed the four terminal taxa involved in these two extreme contrasts—one associated with body mass (between *Syncerus caffer* and *Ammodorcas clarkei*) and the other associated with population density (between *Kobus leche* and *Kobus megaceros*). Given the large sample size (109 species), excluding these four taxa had a negligible impact on overall statistical power. Reanalysis after this exclusion yielded consistent results across all methods, with negative effect estimates and *p*-values below 0.01 in every case (Supplementary Table S7). These results indicate that a small number of highly extreme contrasts can substantially attenuate statistical support for otherwise robust trait associations.

### Runtime Comparison of Phylogenetic Correlation Methods

To assess computational efficiency, we compared the runtime of PIC-O(D)GC with the other five phylogenetic comparative methods across six datasets (Fig. 8). Among all methods, PIC-O(D)GC and PIC-MM showed the smallest increase in runtime as sample size grew, highlighting their strong scalability. The overall computational cost increased from PIC-Pearson to PIC-O(D)GC, PIC-MM, and PGLS, with MR-PMM and Corphylo being the two slowest methods.

**Figure 8.**
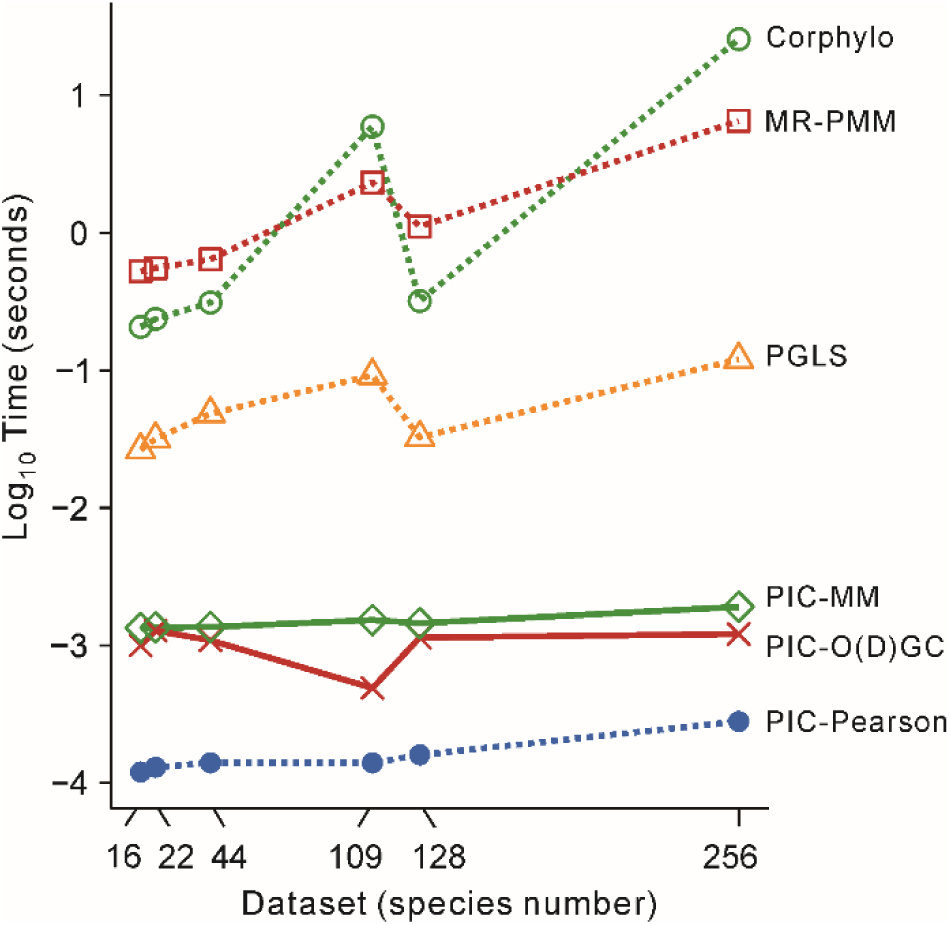
Computational time comparison across phylogenetic correlation methods. Average runtime (in seconds, log10-transformed) was measured for each method across six datasets, including three empirical datasets used in the above analyses (22, 44, and 109 species) and three simulated datasets based on fixed-balanced phylogenetic trees (16, 128, and 256 species). All simulated datasets were generated under the dual-trait shift scenario with an effect size of 4^−3^. Each method was repeated 100 times per dataset to obtain mean values. All runtime measurements were performed on a Dell server running CentOS Linux 7.9, equipped with an AMD EPYC 7702 64-core processor and 1 TB of RAM. Computational time did not always increase with sample size. For instance, both PGLS and Corphylo ran slower on the 109-species dataset than on the 128-species one. This was likely due to relatively long tip labels (e.g., Latin species names) in the 109-species phylogeny, which increased the time required for tree–trait matching. Due to noticeable fluctuations in system load on the shared computational environment, absolute runtimes varied considerably across replicates. However, the relative performance ranking of methods was stable and therefore provides a reliable basis for comparison. The raw timing data used to generate this figure are provided in Supplementary Table S8.

PIC-Pearson was the fastest methods, benefiting from linear computational complexity.

PIC-O(D)GC also exhibited low computational cost, relying on median-based outlier detection (O(n)) and, depending on outlier presence, applying either Pearson or Spearman correlation. When Spearman correlation is used, an O(n log n) cost is incurred due to ranking, which remains efficient even for large trees. In small datasets, a normality test (e.g., Shapiro–Wilk) is additionally applied to guide correlation choice, introducing modest overhead that is most apparent at small sample sizes. PIC-MM required more computation than PIC-O(D)GC, consistent with its robust two-stage estimation procedure: an initial S-estimation step (up to O(n^3^)), followed by iterative M-estimation steps (each O(n^2^)). PGLS was even slower than PIC-MM, driven by repeated matrix inversions. The remaining model-based approaches—MR-PMM, and Corphylo—incurred the highest computational time overall. For example, on the largest dataset (*n* = 256), PIC-Pearson, PIC-O(D)GC, and PIC-MM completed in 3×10⁻^4^, 10⁻^3^, and 2×10⁻^3^ seconds, respectively, while PGLS and MR-PMM required 0.12 and 6.5 seconds, respectively. Corphylo was particularly slow, taking 26 seconds on the same dataset.

These results underscore the scalability of PIC-O(D)GC, which achieves high computational efficiency without compromising robustness. This dual strength makes it particularly suitable for large-scale phylogenetic comparative analyses.

## Discussion

### Equivalence of Pearson Correlation and OLS in Context

Both Pearson correlation and OLS regression can be applied to PIC to assess trait associations in a phylogenetically aware manner (Felsenstein 1985, Felsenstein 1988, Murray et al. 2020, Horvath et al. 2022, Bai et al. 2023, Wang et al. 2023). When applied to PIC data under linear assumptions, these two approaches yield mathematically equivalent results in terms of statistical significance and correlation direction. In this study, we chose to use Pearson correlation for its computational simplicity and consistency with other nonparametric methods, such as Spearman correlation, which is a key component of the PIC-O(D)GC workflow. The equivalence between Pearson correlation and OLS regression ensures logical and statistical consistency with previous findings while reinforcing the relevance and robustness of our results within the context of phylogenetic comparative analyses.

### Comparing Statistical Methods for Detecting Trait Correlations

Our study evaluated the performance of various statistical methods in detecting trait correlations under diverse evolutionary scenarios, including with and without abrupt shifts.

PIC-O(D)GC and PIC-MM consistently demonstrated robust performance, particularly excelling under shift scenarios by minimizing false positives as well as false negatives, thereby maintaining high overall accuracy—consistent with prior evaluations of robust regression strategies in the presence of model violations and trait shifts (Adams et al. 2024). In contrast, traditional methods like PIC-Pearson and PGLS, as well as robust variants such as PGLS-MM, struggled under high shift gradients, highlighting their limitations in challenging conditions. In no-shift scenarios, both PIC-MM and PIC-O(D)GC performed comparably to other reliable methods, such as PIC-Pearson, PGLS, and Corphylo, underscoring that neither method was uniformly superior across all conditions. These complementary strengths highlight the importance of aligning method choice with expected data characteristics and analytical goals.

Notably, the abrupt trait shifts introduced in our simulations can be considered an extreme form of evolutionary heterogeneity—a concept emphasized by Mazel et al. (2016) as a key source of model violation in phylogenetic regression. While Mazel et al. focused on rate heterogeneity, our results show that discrete directional shifts can also distort inference and compromise methods assuming homogeneous evolutionary processes. This underscores the importance of methods capable of accommodating both continuous and discontinuous heterogeneity.

### Conceptual Simplicity and Practical Accessibility

In addition to the substantial runtime advantage demonstrated in Figure 8, PIC-O(D)GC also offers strengths in conceptual simplicity and user accessibility. By dynamically adapting to the presence or absence of outliers and data normality, it strikes an effective balance between robustness and interpretability. Its reliance on straightforward statistical principles, such as Spearman’s rank correlation, ensures that the method is logically intuitive and easy to understand, even for researchers with limited statistical training (Redmond and Keenan 2002). By avoiding rigid assumptions about data distribution, PIC-O(D)GC provides transparent and interpretable results, reducing the risk of misinterpretation and enhancing its accessibility to a broader range of biologists.

### From Statistical Artifacts to Evolutionary Signals: Interpreting PIC Outliers

Beyond its utility in assessing trait correlations, the outlier detection step in PIC-O(D)GC holds promise for identifying evolutionary shifts or jumps. Recent studies have highlighted the importance of models that accommodate abrupt changes in trait evolution, such as the singular events model (Uyeda et al. 2018). These rare but significant transitions may leave identifiable signatures in PIC data (Uyeda et al. 2018), including extreme contrast values, or pronounced localized changes along specific branches.

The importance of examining PIC scatterplots cannot be overstated. Even when an overall correlation appears statistically significant, the result may be disproportionately driven by a few extreme contrast values. Visual inspection of the PIC plot is therefore essential, not only as a diagnostic check, but also as a guide to more structured interpretation.

When such outliers are present, we recommend analyzing the relationship between two traits from three complementary perspectives: 1) Overall correlation can be assessed using robust methods such as PIC-MM or PIC-O(D)GC, which account for outliers and non-normality while maintaining statistical stability across heterogeneous evolutionary scenarios; 2) Shift-to-shift relationships should be interpreted separately, considering whether shifts are aligned or opposed in direction, which may reveal patterns of evolutionary coupling or divergence. 3) Background correlation refers to the relationship between traits during non-shift, gradual phases of evolution.

When a shift is located deep in the phylogeny—particularly near the root, as seen in Figure 1 (panels d, f, j, l)—an effective strategy is to partition the data into two clades at the shift node. This allows separate estimation of trait relationships within each group, providing insight into background evolutionary dynamics before and after the divergence introduced by the shift.

For shifts occurring on terminal or near-terminal branches (Fig. 3), we observed that PIC-O(D)GC, and even PIC-MM, may exhibit reduced sensitivity, reflecting the disproportionate influence of a small number of localized, large-effect contrasts. In these cases, removing the taxa involved in the detected shift can be an effective and often necessary step to limit their impact and recover the underlying background correlation structure.

Once the influence of shifts is appropriately filtered—whether through clade partitioning or tip removal—regression-based methods such as PIC-OLS, or PGLS (Felsenstein 1985, Martins and Hansen 1997, Pagel 1997, Pagel 1999, Ives et al. 2007, Revell 2010) become better suited for background-phase analysis by leveraging continuous trait variation and enabling effect-size estimation and model-based inference. In practice, the empirical examples of body mass and population density in even-toed ungulates and SO and FG muscle fiber proportions in lizard species illustrate how such regression-based approaches can provide clear directionality and statistically supported assessments of trait associations once shift effects are appropriately filtered. These regression-based approaches are preferred in background-phase analysis due to their interpretability and modeling efficiency. In contrast, robust contrast-based methods like PIC-O(D)GC or PIC-MM (Adams et al. 2024) may be overly conservative in this refined context.

Empirical evidence increasingly supports the idea that phenotypic evolution involves rare but consequential jumps embedded within longer periods of gradual change or stasis. Such discontinuities have been observed across a range of systems, including body size evolution in crocodylomorphs (Godoy et al. 2019), bursts of morphological innovation in lizards (Brennan et al. 2024), shifts in venom gene evolution linked to ecological opportunity in snakes (Barua and Mikheyev 2020), and abrupt changes in genomic traits such as GC content and genome size in bacteria (Gao and Wu 2022, Hu et al. 2022, Mahajan and Agashe 2022). These cases underscore the importance of not only identifying where and when such evolutionary shifts occur, but also understanding how they alter patterns of trait covariation. In particular, they highlight the need to distinguish among overall correlations across the tree, background correlations within stable lineages, and shift-to-shift relationships that may reflect directional evolutionary coupling or divergence between regimes.

The combined use of contrast-based diagnostics such as PIC-O(D)GC and targeted analyses of trait relationships offers a tractable workflow for detecting and interpreting evolutionary pulses. Unlike fully model-based methods, PIC analysis leverages branch-level resolution to identify candidate shifts without prior assumptions about evolutionary regimes. This approach complements theoretical and algorithmic models such as OU-with-jumps (Landis et al. 2013), Lévy process-based jump models (Duchen et al. 2017, Landis and Schraiber 2017), LASSO-regularized shift detection (Khabbazian et al. 2016, Bastide et al. 2018, Zhang et al. 2024), and singular events models (Uyeda et al. 2011, Uyeda et al. 2018), all of which recognize discrete shifts as critical components of macroevolutionary dynamics.

### Limitations and Future Directions

While correlation-based methods such as PIC-O(D)GC are robust to non-normal data and outliers, they are inherently limited to detecting monotonic associations and do not provide parameter estimates or accommodate multiple predictors. Thus, they should be viewed as complementary to parametric models like PGLS, which remain essential for estimating the strength and functional form of evolutionary relationships.

Further refinement of this workflow may be achieved by incorporating phylogenetic features such as branch lengths and trait-specific evolutionary rates. These additions could improve the specificity of shift detection and help distinguish genuine evolutionary transitions from background variation. Integrating PIC outlier diagnostics into hypothesis-driven macroevolutionary studies may thus provide a more nuanced view of how trait relationships evolve across lineages.

## Supporting information

All the supplementary figures.

Codes used in this study.

All the supplementary tables.

Supplementary results.

## Acknowledgments

We sincerely thank Professor Josef Uyeda (Associate Editor) and the anonymous reviewer for their constructive and insightful comments. Many sentences in this article were drafted with the assistance of ChatGPT-4 and its later versions, large language models developed by OpenAI. The authors thoroughly reviewed and revised each sentence and took full responsibility for the language and content of the entire article.

## Data availability

Supplementary Methods and supplementary analyses are provided in Supplementary_Methods_S1.pdf and Supplementary_Results_S1.pdf, respectively. Supplementary Figures S1–S12 are available in Supplementary_Figures.pdf. All data supporting the figures presented in the main text and supplementary materials, including Tables S1–S28, are provided in Supplementary_Tables.xlsx. All source code used for simulations and analyses is available in Code.zip. We also have implemented a suite of functions for conducting PIC-O(D)GC in the R package PICODGC (Phylogenetic Independent Contrasts with Outlier-and Distribution-Guided Correlation), freely available at https://github.com/zhenglinchen/PICODGC.

## References

Adams R., Cain Z., Assis R., DeGiorgio M. 2024. Robust phylogenetic regression. Syst. Biol., 73:140–157.

Bai X.-L., Yang D., Sher J., Zhang Y.-B., Zhang K.-Y., Liu Q., Wen H.-D., Zhang J.-L., Slot M. 2023. Divergences in stem and leaf traits between lianas and coexisting trees in a subtropical montane forest. Journal of Plant Ecology, 17:rtad037.

Barua A., Mikheyev A.S. 2020. Toxin expression in snake venom evolves rapidly with constant shifts in evolutionary rates. Proc. R. Soc. B, 287:20200613.

Bastide P., Ané C., Robin S., Mariadassou M. 2018. Inference of adaptive shifts for multivariate correlated traits. Syst. Biol., 67:662–680.

Blomberg S.P., Lefevre J.G., Wells J.A., Waterhouse M. 2012. Independent contrasts and PGLS regression estimators are equivalent. Syst. Biol., 61:382–391.

Brennan I.G., Chapple D.G., Keogh J.S., Donnellan S. 2024. Evolutionary bursts drive morphological novelty in the world’s largest skinks. Curr. Biol., 34:3905–3916.e3905.

Cornwallis C.K., Griffin A.S. 2024. A guided tour of phylogenetic comparative methods for studying trait evolution. Annu. Rev. Ecol. Evol. Syst., 55:181–204.

Dewar A.E., Belcher L.J., West S.A. 2025. A phylogenetic approach to comparative genomics. Nat. Rev. Genet., 26:395–405.

Duchen P., Leuenberger C., Szilagyi S.M., Harmon L., Eastman J., Schweizer M., Wegmann D. 2017. Inference of evolutionary jumps in large phylogenies using Levy processes. Syst. Biol., 66:950–963.

Felsenstein J. 1985. Phylogenies and the comparative method. Amer. Natur., 125:1–15.

Felsenstein J. 1988. Phylogenies and quantitative characters. Annu. Rev. Ecol. Syst., 19:445–471.

Freckleton R.P., Harvey P.H., Pagel M. 2002. Phylogenetic analysis and comparative data: A test and review of evidence. Amer. Natur., 160:712–726.

Gao Y., Wu M. 2022. Microbial genomic trait evolution is dominated by frequent and rare pulsed evolution. Sci. Adv., 8:eabn1916.

Garamszegi L.Z. 2014. Modern Phylogenetic Comparative Methods and Their Application in Evolutionary Biology: Concepts and Practice. Berlin, Springer.

Garland T., Jr, Harvey P.H., Ives A.R. 1992. Procedures for the analysis of comparative data using phylogenetically independent contrasts. Syst. Biol., 41:18–32.

Godoy P.L., Benson R.B.J., Bronzati M., Butler R.J. 2019. The multi-peak adaptive landscape of crocodylomorph body size evolution. BMC Evol. Biol., 19:167.

Grafen A. 1989. The phylogenetic regression. Philos. Trans. R. Soc. Lond. B Biol. Sci., 326:119–157.

Hadfield J.D. 2010. MCMC methods for multi-response generalized linear mixed models: the MCMCglmm R package. J. Stat. Softw., 33:1–22.

Hadfield J.D., Nakagawa S. 2010. General quantitative genetic methods for comparative biology: phylogenies, taxonomies and multi-trait models for continuous and categorical characters. J. Evol. Biol., 23:494–508.

Halliwell B., Holland B.R., Yates L.A. 2025. Multi-response phylogenetic mixed models: concepts and application. Biol. Rev., 100:1294–1316.

Hansen T.F. 1997. Stabilizing selection and the comparative analysis of adaptation. Evolution, 51:1341–1351.

Harmon L.J., Losos J.B., Jonathan Davies T., Gillespie R.G., Gittleman J.L., Bryan Jennings W., Kozak K.H., McPeek M.A., Moreno-Roark F., Near T.J., Purvis A., Ricklefs R.E., Schluter D., Schulte Ii J.A., Seehausen O., Sidlauskas B.L., Torres-Carvajal O., Weir J.T., Mooers A.Ø. 2010. Early bursts of body size and shape evolution are rare in comparative data. Evolution, 64:2385–2396.

Ho L.S.T., Ane C. 2014. A linear-time algorithm for Gaussian and non-Gaussian trait evolution models. Syst. Biol., 63:397–408.

Hoaglin D.C., Iglewicz B., Tukey J.W. 1986. Performance of Some Resistant Rules for Outlier Labeling. J Am Stat Assoc, 81:991–999.

Horvath S., Lu A.T., Haghani A., Zoller J.A., Li C.Z., Lim A.R., Brooke R.T., Raj K., Serres-Armero A., Dreger D.L., Hogan A.N., Plassais J., Ostrander E.A. 2022. DNA methylation clocks for dogs and humans. Proc. Natl. Acad. Sci. USA, 119:e2120887119.

Hu E.-Z., Lan X.-R., Liu Z.-L., Gao J., Niu D.-K. 2022. A positive correlation between GC content and growth temperature in prokaryotes. BMC Genomics, 23:110.

Ives A.R. 2022. Random errors are neither: On the interpretation of correlated data. Methods Ecol. Evol., 13:2092–2105.

Ives A.R., Helmus M.R. 2011. Generalized linear mixed models for phylogenetic analyses of community structure. Ecol. Monographs, 81:511–525.

Ives A.R., Midford P.E., Garland T., Jr. 2007. Within-species variation and measurement error in phylogenetic comparative methods. Syst. Biol., 56:252–270.

Jones K.E., Bielby J., Cardillo M., Fritz S.A., O’Dell J., Orme C.D.L., Safi K., Sechrest W., Boakes E.H., Carbone C., Connolly C., Cutts M.J., Foster J.K., Grenyer R., Habib M., Plaster C.A., Price S.A., Rigby E.A., Rist J., Teacher A., Bininda-Emonds O.R.P., Gittleman J.L., Mace G.M., Purvis A. 2009. PanTHERIA: a species-level database of life history, ecology, and geography of extant and recently extinct mammals. Ecology, 90:2648–2648.

Khabbazian M., Kriebel R., Rohe K., Ané C. 2016. Fast and accurate detection of evolutionary shifts in Ornstein–Uhlenbeck models. Methods Ecol. Evol., 7:811–824.

Landis M.J., Schraiber J.G. 2017. Pulsed evolution shaped modern vertebrate body sizes. Proc. Natl. Acad. Sci. USA, 114:13224–13229.

Landis M.J., Schraiber J.G., Liang M. 2013. Phylogenetic analysis using Levy processes: finding jumps in the evolution of continuous traits. Syst. Biol., 62:193–204.

Lynch M. 1991. Methods for the analysis of comparative data in evolutionary biology. Evolution, 45:1065–1080.

Mahajan S., Agashe D. 2022. Evolutionary jumps in bacterial GC content. G3 Genes|Genomes|Genetics, 12:jkac108.

Makino T., Kawata M. 2019. Invasive invertebrates associated with highly duplicated gene content. Mol. Ecol., 28:1652–1663.

Martins E.P., Hansen T.F. 1997. Phylogenies and the comparative method: A general approach to incorporating phylogenetic information into the analysis of interspecific data. Amer. Natur., 149:646–667.

Mazel F., Davies T.J., Georges D., Lavergne S., Thuiller W., Peres-Neto P.R. 2016. Improving phylogenetic regression under complex evolutionary models. Ecology, 97:286–293.

Murray G.G.R., Charlesworth J., Miller E.L., Casey M.J., Lloyd C.T., Gottschalk M., Tucker A.W., Welch J.J., Weinert L.A. 2020. Genome reduction is associated with bacterial pathogenicity across different scales of temporal and ecological divergence. Mol. Biol. Evol., 38:1570–1579.

O’Meara B. 2016. Phylogenetic Comparative Method. In: Kliman RM editor. Encyclopedia of Evolutionary Biology. Oxford, Academic Press, p. 254–256.

Pagel M. 1997. Inferring evolutionary processes from phylogenies. Zool. Scr., 26:331–348.

Pagel M. 1999. Inferring the historical patterns of biological evolution. Nature, 401:877–884.

Paradis E., Schliep K. 2019. ape 5.0: an environment for modern phylogenetics and evolutionary analyses in R. Bioinformatics, 35:526–528.

R Core Team. 2020. R: A language and environment for statistical computing. R Foundation for Statistical Computing, Vienna, Austria. https://www.R-project.org/.

Redmond A.C., Keenan A.M. 2002. Understanding statistics. Putting p-values into perspective. J Am Podiatr Med Assoc, 92:297–305.

Revell L.J. 2010. Phylogenetic signal and linear regression on species data. Methods Ecol. Evol., 1:319–329.

Revell L.J. 2024. phytools 2.0: an updated R ecosystem for phylogenetic comparative methods (and other things). PeerJ, 12:e16505.

Rohlf F.J. 2001. Comparative methods for the analysis of continuous variables: geometric interpretations. Evolution, 55:2143–2160.

Romiguier J., Gayral P., Ballenghien M., Bernard A., Cahais V., Chenuil A., Chiari Y., Dernat R., Duret L., Faivre N., Loire E., Lourenco J.M., Nabholz B., Roux C., Tsagkogeorga G., Weber A.A.T., Weinert L.A., Belkhir K., Bierne N., Glémin S., Galtier N. 2014. Comparative population genomics in animals uncovers the determinants of genetic diversity. Nature, 515:261–263.

Royston P. 1992. Approximating the Shapiro-Wilk W-test for non-normality. Statistics and Computing, 2:117–119.

Scales J.A., King A.A., Butler M.A. 2009. Running for your life or running for your dinner: what drives fiber-type evolution in lizard locomotor muscles? Amer. Natur., 173:543–553.

Slater G.J., Pennell M.W. 2014. Robust regression and posterior predictive simulation increase power to detect early bursts of trait evolution. Syst. Biol., 63:293–308.

Sorensen D., Gianola D. 2002. Likelihood, Bayesian, and MCMC Methods in Quantitative Genetics. New York, Springer.

Sumner S., Favreau E., Geist K., Toth A.L., Rehan S.M. 2023. Molecular patterns and processes in evolving sociality: lessons from insects. Philos Trans R Soc Lond B Biol Sci, 378:20220076.

Symonds M.R.E., Blomberg S.P. 2014. A primer on phylogenetic generalised least squares. In: Garamszegi LZ editor. Modern Phylogenetic Comparative Methods and Their Application in Evolutionary Biology: Concepts and Practice. Berlin, Heidelberg, Springer Berlin Heidelberg, p. 105–130.

Tukey J.W. 1977. Exploratory Data Analysis. Reading, MA, Addison-Wesley.

Upham N.S., Esselstyn J.A., Jetz W. 2019. Inferring the mammal tree: Species-level sets of phylogenies for questions in ecology, evolution, and conservation. PLoS Biol., 17:e3000494.

Uyeda J.C., Hansen T.F., Arnold S.J., Pienaar J. 2011. The million-year wait for macroevolutionary bursts. Proc. Natl. Acad. Sci. USA, 108:15908–15913.

Uyeda J.C., Zenil-Ferguson R., Pennell M.W. 2018. Rethinking phylogenetic comparative methods. Syst. Biol., 67:1091–1109.

Wang B., Tong Z.-Y., Xiong Y.-Z., Wang X.-F., Scott Armbruster W., Huang S.-Q. 2023. The evolution of flower–pollinator trait matching, and why do some alpine gingers appear to be mismatched? Ann. Bot., 132:1073–1088.

Westoby M., Yates L., Holland B., Halliwell B. 2023. Phylogenetically conservative trait correlation: Quantification and interpretation. J Ecol, 111:2105–2117.

Zhang W., Kenney T., Ho L.S.T. 2024. Evolutionary shift detection with ensemble variable selection. BMC Ecology and Evolution, 24:11.

Zheng L., Ives A.R., Garland T., Larget B.R., Yu Y., Cao K. 2009. New multivariate tests for phylogenetic signal and trait correlations applied to ecophysiological phenotypes of nine *Manglietia* species. Funct. Ecol., 23:1059–1069.

