## Supplementary material for "Improving the Robustness of Phylogenetic Independent Contrasts: Addressing Abrupt Evolutionary Shifts with Outlier-and Distribution-Guided Correlation": All the supplementary figures.

### Supplementary Figures S1-S12

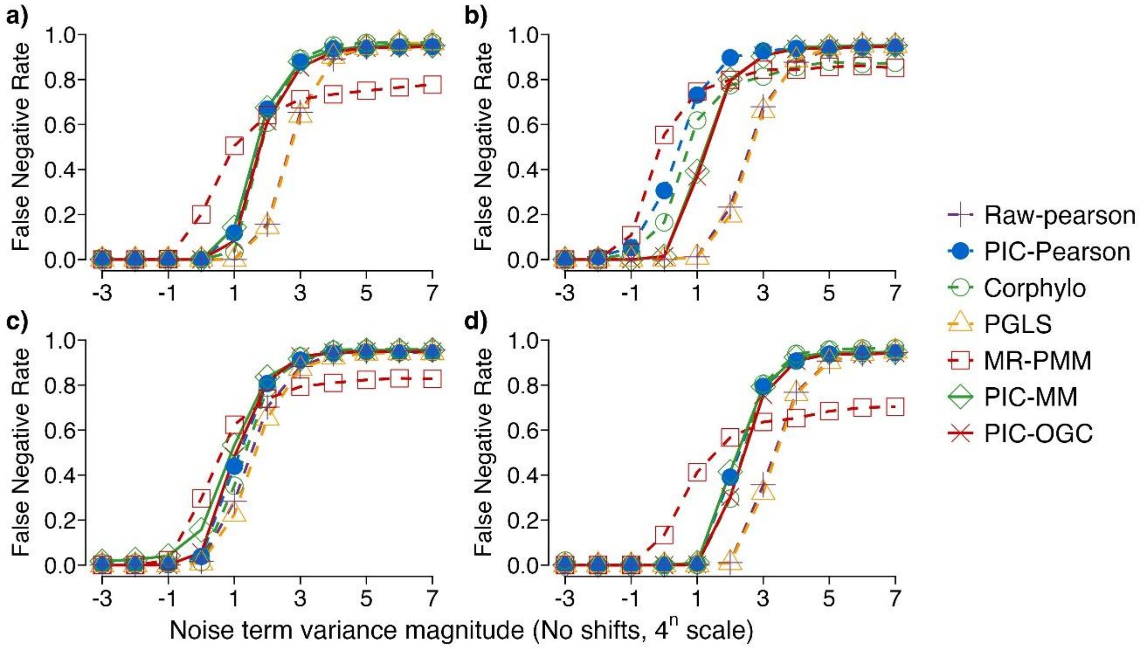

Supplementary Figure S1. Error rates of seven methods under the BM + Norm scenario evaluated on four phylogenetic settings using the $\beta$-based benchmark.

**a)** a fixed-balanced tree with 128 species; **b)** an ensemble of random trees with 128 species; **c)** a fixed-balanced tree with 16 species; and **d)** a fixed-balanced tree with 256 species. The horizontal axis shows the variance of the added noise term as $4^n$; only the exponent n is printed (e.g. $n$=−3 represents $4^{-3}$, and $n$=7 represents $4^7$). “Raw-Pearson” refers to applying the standard Pearson correlation directly to the original species-level trait data, without any phylogenetic transformation. Additional details on the simulation design, gradient configurations, and performance metrics are provided in the Materials and Methods. The data used to plot this figure are available in Supplementary Table S9.

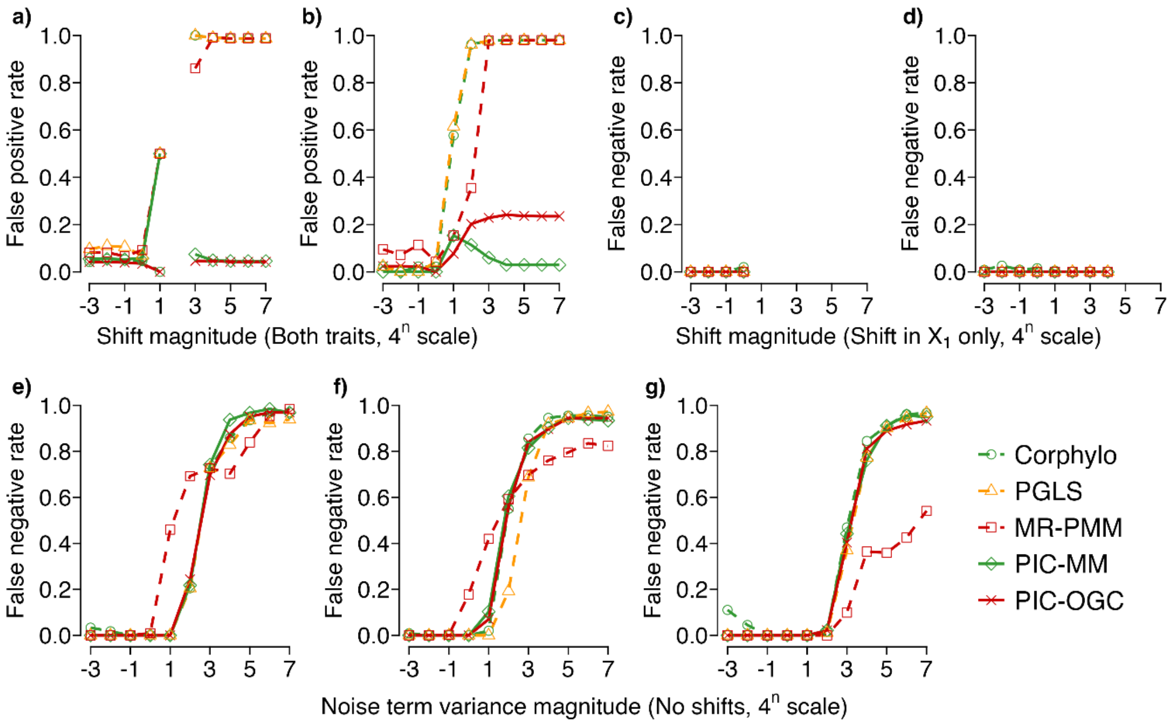

Supplementary Figure S2. Performance of five methods on datasets that deviate from Brownian motion (BM) assumptions on a 128-species fixed-balanced tree using the $\beta$-based benchmark.

Panels **a–b** show false positive rates under non-BM datasets with evolutionary shifts in both $X_1$ and $X_2$, where shifts are introduced at the root branch (Felsenstein's worst-case scenario; **a**) or at randomly selected branch locations (**b**). Panels **c–d** show false negative rates under non-BM datasets with evolutionary shifts only in $X_1$, again comparing worst-case root shifts (**c**) and randomly located shifts (**d**). Panels **e–g** show false negative rates under non-BM datasets without abrupt evolutionary shifts, across three gradual-evolution scenarios: BM–BM (panel **e**), BM–Norm (panel **f**), and Norm–BM (panel **g**). Shift and noise variance values are represented as $4^n$, with the horizontal axis showing only the exponent $n$ (e.g., n = −3 corresponds to $4^{-3}$, and n = 7 corresponds to $4^7$). All datasets were simulated on a fixed-balanced phylogenetic tree with 128 species. See the Materials and Methods for additional details on the simulation design, gradient configurations, and performance metrics. The data used to generate this figure are provided in Supplementary Table S10**.**

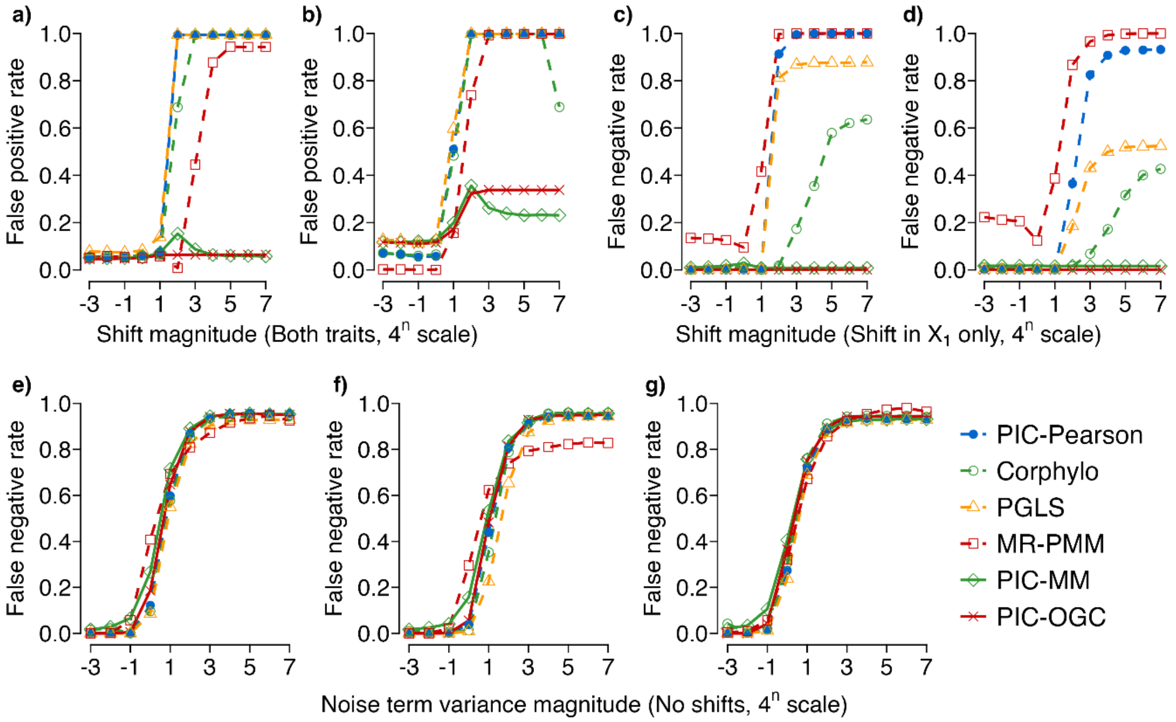

Supplementary Figure S3. Performance of six methods across evolutionary shift and no-shift regimes simulated on a 16-species fixed-balanced tree using the $\beta$-based benchmark.

Panels **a** and **c** correspond to Felsenstein's worst-case scenario, in which directional shifts are introduced on deep-root branches, whereas panels **b** and **d** show results from simulations with randomly located trait shifts. Panels **a**–**b** report false positive rates under simultaneous shifts in both $X_1$ and $X_2$, and panels **c**–**d** report false negative rates when only $X_1$ is shifted. Panels **e**–**g** show performance under gradual evolutionary scenarios without abrupt shifts, in which traits follow the relationship $X_2 = X_1 + \varepsilon$ with increasing noise variance. Three evolutionary background conditions are considered: BM–BM (panel **e**), BM–Norm (panel **f**), and Norm–BM (panel **g**). Shift and noise variance magnitudes are expressed on a $4^n$ scale, where the horizontal axis shows only the exponent ($n$) rather than the full value. Additional details on the simulation design, gradient configurations, and performance metrics are provided in the Materials and Methods. The data used to generate this figure are available in Supplementary Table S11.

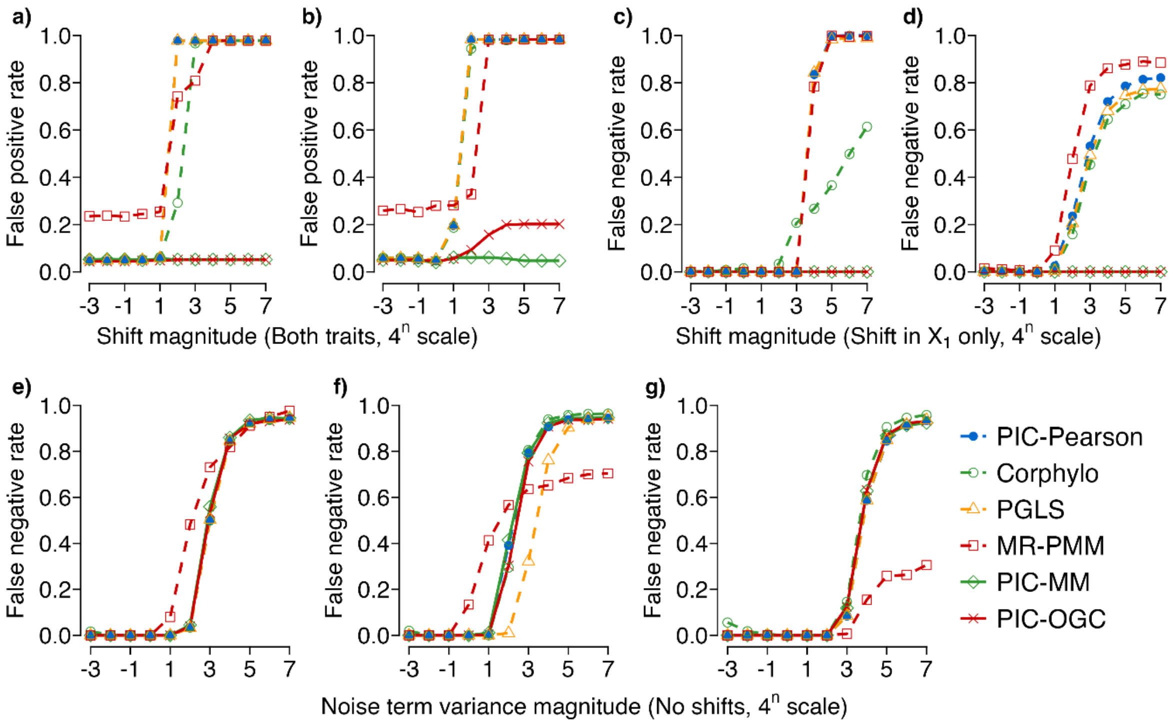

Supplementary Figure S4. Performance of six methods across evolutionary shift and no-shift regimes simulated on a 256-species fixed-balanced tree using the β-based benchmark.

Panels **a** and **c** correspond to Felsenstein's worst-case scenario, in which directional shifts are introduced on deep-root branches, whereas panels **b** and **d** show results from simulations with randomly located trait shifts. Panels **a**–**b** report false positive rates under simultaneous shifts in both $X_1$ and $X_2$, and panels **c**–**d** report false negative rates when only $X_1$ is shifted. Panels **e**–**g** show performance under gradual evolutionary scenarios without abrupt shifts, in which traits follow the relationship $X_2 = X_1 + \varepsilon$ with increasing noise variance. Three evolutionary background conditions are considered: BM–BM (panel **e**), BM–Norm (panel **f**), and Norm–BM (panel **g**). Shift and noise variance magnitudes are expressed on a $4^n$ scale, where the horizontal axis shows only the exponent ($n$) rather than the full value. Additional details on the simulation design, gradient configurations, and performance metrics are provided in the Materials and Methods. The data used to generate this figure are available in Supplementary Table S12.

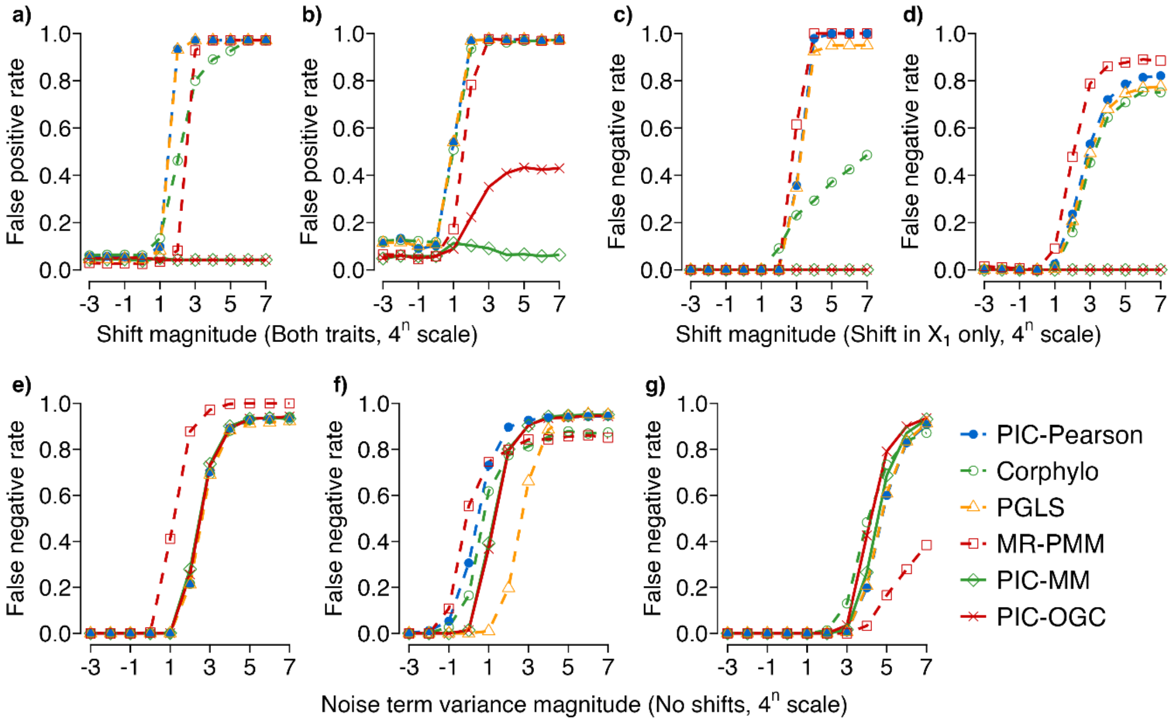

Supplementary Figure S5. Performance of six methods across evolutionary shift and no-shift regimes simulated on 128-species random trees using the β-based benchmark.

Panels **a** and **c** correspond to Felsenstein's worst-case scenario, in which directional shifts are introduced on deep-root branches, whereas panels **b** and **d** show results from simulations with randomly located trait shifts. Panels **a**–**b** report false positive rates under simultaneous shifts in both $X_1$ and $X_2$, and panels **c**–**d** report false negative rates when only $X_1$ is shifted. Panels **e**–**g** show performance under gradual evolutionary scenarios without abrupt shifts, in which traits follow the relationship $X_2 = X_1 + \varepsilon$ with increasing noise variance. Three evolutionary background conditions are considered: BM–BM (panel **e**), BM–Norm (panel **f**), and Norm–BM (panel **g**). Shift and noise variance magnitudes are expressed on a $4^n$ scale, where the horizontal axis shows only the exponent ($n$) rather than the full value. Additional details on the simulation design, gradient configurations, and performance metrics are provided in the Materials and Methods. The data used to generate this figure are available in Supplementary Table S13.

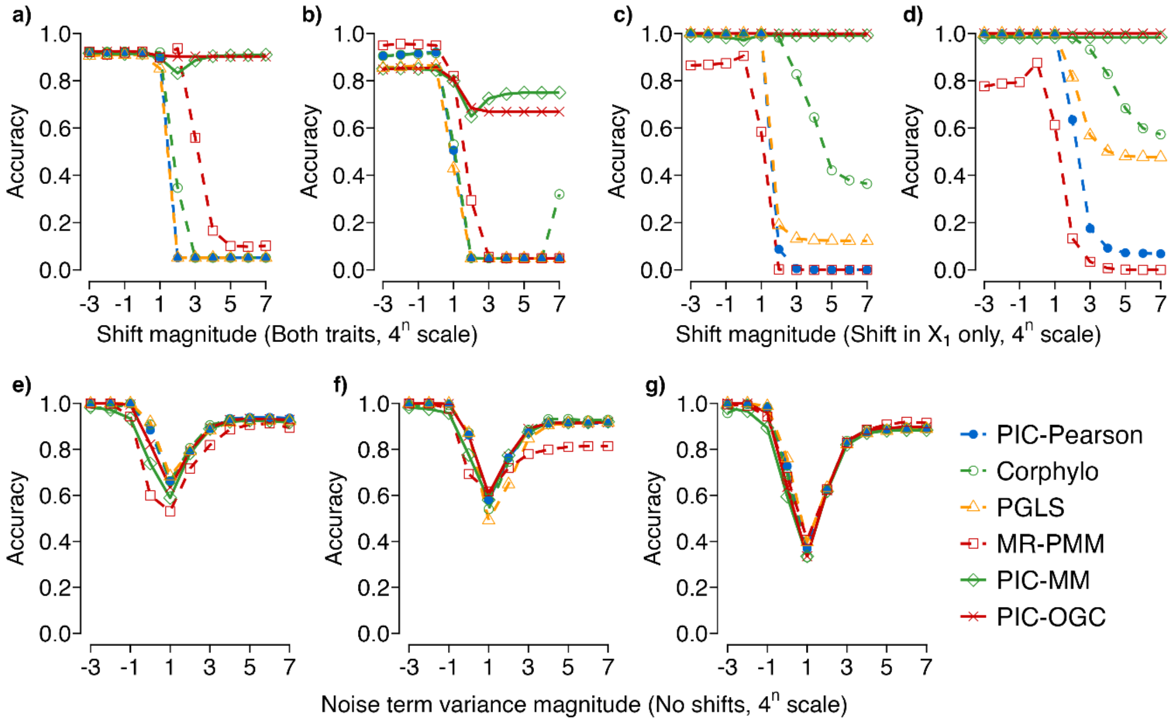

Supplementary Figure S6. Performance of six methods across evolutionary shift and no-shift regimes simulated on a 16-species fixed-balanced tree using the data-pattern–based benchmark.

Panels **a** and **c** correspond to Felsenstein's worst-case scenario, in which directional shifts are introduced on deep-root branches, whereas panels **b** and **d** show results from simulations with randomly located trait shifts. Panels **a**–**b** show performance under simultaneous shifts in both $X_1$ and $X_2$, and panels **c**–**d** show performance when only $X_1$ is shifted. Panels **e**–**g** show performance under gradual evolutionary scenarios without abrupt shifts, in which traits follow the relationship $X_2 = X_1 + \varepsilon$ with increasing noise variance. Three evolutionary background conditions are considered: BM–BM (panel **e**), BM–Norm (panel **f**), and Norm–BM (panel **g**). Shift and noise variance magnitudes are expressed on a $4^n$ scale, where the horizontal axis shows only the exponent ($n$) rather than the full value. Additional details on the simulation design, gradient configurations, and performance metrics are provided in the Materials and Methods. The data used to generate this figure are available in Supplementary Table S14.

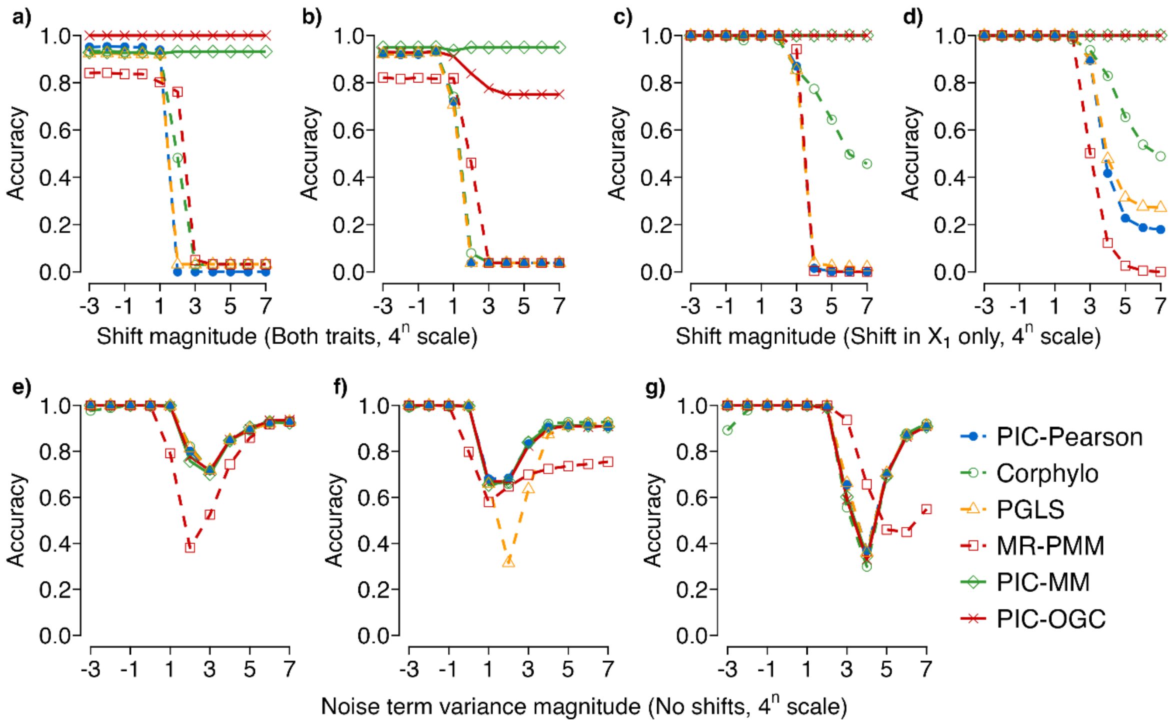

Supplementary Figure S7. Performance of six methods across evolutionary shift and no-shift regimes simulated on a 128-species fixed-balanced tree using the data-pattern–based benchmark.

Panels **a** and **c** correspond to Felsenstein's worst-case scenario, in which directional shifts are introduced on deep-root branches, whereas panels **b** and **d** show results from simulations with randomly located trait shifts. Panels **a**–**b** show performance under simultaneous shifts in both $X_1$ and $X_2$, and panels **c**–**d** show performance when only $X_1$ is shifted. Panels **e**–**g** show performance under gradual evolutionary scenarios without abrupt shifts, in which traits follow the relationship $X_2 = X_1 + \varepsilon$ with increasing noise variance. Three evolutionary background conditions are considered: BM–BM (panel **e**), BM–Norm (panel **f**), and Norm–BM (panel **g**). Shift and noise variance magnitudes are expressed on a $4^n$ scale, where the horizontal axis shows only the exponent ($n$) rather than the full value. Additional details on the simulation design, gradient configurations, and performance metrics are provided in the Materials and Methods. The data used to generate this figure are available in Supplementary Table S15.

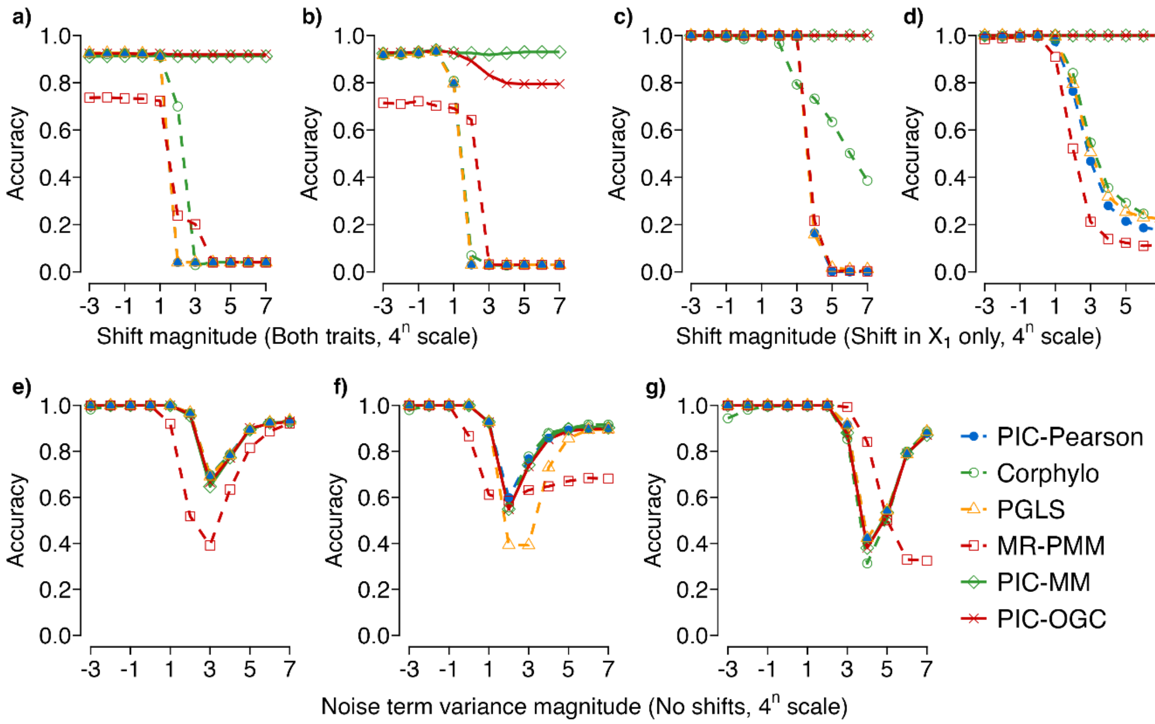

Supplementary Figure S8. Performance of six methods across evolutionary shift and no-shift regimes simulated on a 256-species fixed-balanced tree using the data-pattern–based benchmark.

Panels **a** and **c** correspond to Felsenstein's worst-case scenario, in which directional shifts are introduced on deep-root branches, whereas panels **b** and **d** show results from simulations with randomly located trait shifts. Panels **a**–**b** show performance under simultaneous shifts in both $X_1$ and $X_2$, and panels **c**–**d** show performance when only $X_1$ is shifted. Panels **e**–**g** show performance under gradual evolutionary scenarios without abrupt shifts, in which traits follow the relationship $X_2 = X_1 + \varepsilon$ with increasing noise variance. Three evolutionary background conditions are considered: BM–BM (panel **e**), BM–Norm (panel **f**), and Norm–BM (panel **g**). Shift and noise variance magnitudes are expressed on a $4^n$ scale, where the horizontal axis shows only the exponent ($n$) rather than the full value. Additional details on the simulation design, gradient configurations, and performance metrics are provided in the Materials and Methods. The data used to generate this figure are available in Supplementary Table S16.

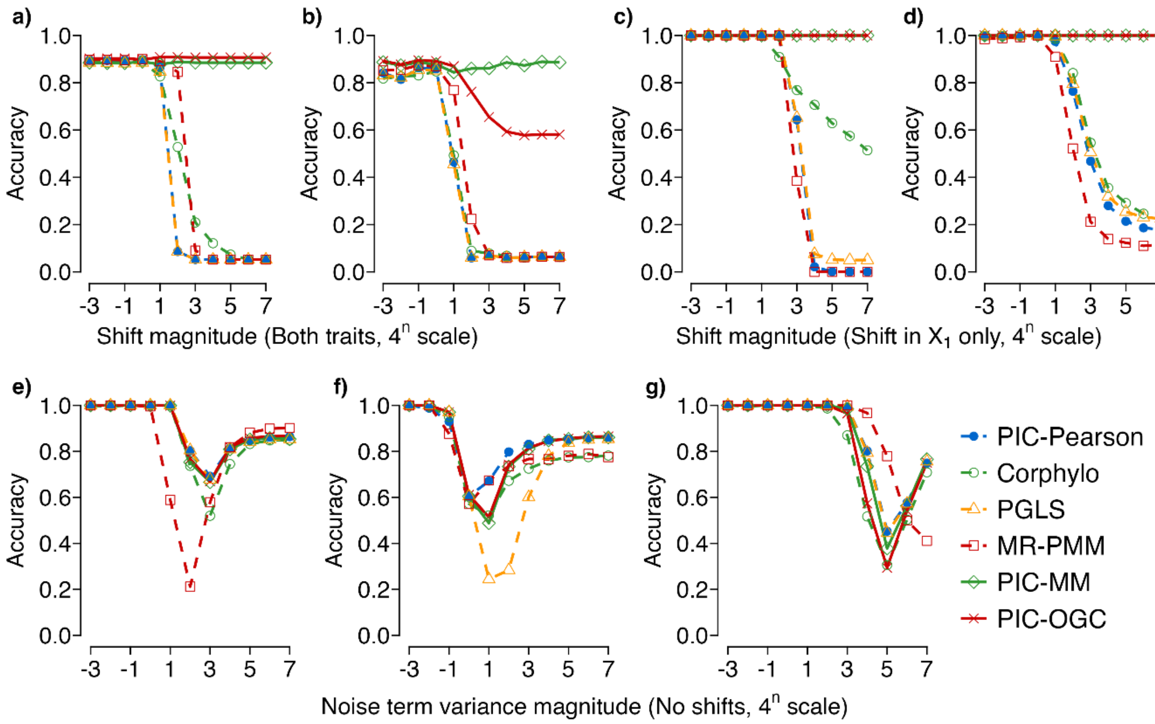

Supplementary Figure S9. Performance of six methods across evolutionary shift and no-shift regimes simulated on 128-species random trees using the data-pattern–based benchmark.

Panels **a** and **c** correspond to Felsenstein's worst-case scenario, in which directional shifts are introduced on deep-root branches, whereas panels **b** and **d** show results from simulations with randomly located trait shifts. Panels **a**–**b** show performance under simultaneous shifts in both $X_1$ and $X_2$, and panels **c**–**d** show performance when only $X_1$ is shifted. Panels **e**–**g** show performance under gradual evolutionary scenarios without abrupt shifts, in which traits follow the relationship $X_2 = X_1 + \varepsilon$ with increasing noise variance. Three evolutionary background conditions are considered: BM–BM (panel **e**), BM–Norm (panel **f**), and Norm–BM (panel **g**). Shift and noise variance magnitudes are expressed on a $4^n$ scale, where the horizontal axis shows only the exponent ($n$) rather than the full value. Additional details on the simulation design, gradient configurations, and performance metrics are provided in the Materials and Methods. The data used to generate this figure are available in Supplementary Table S17.

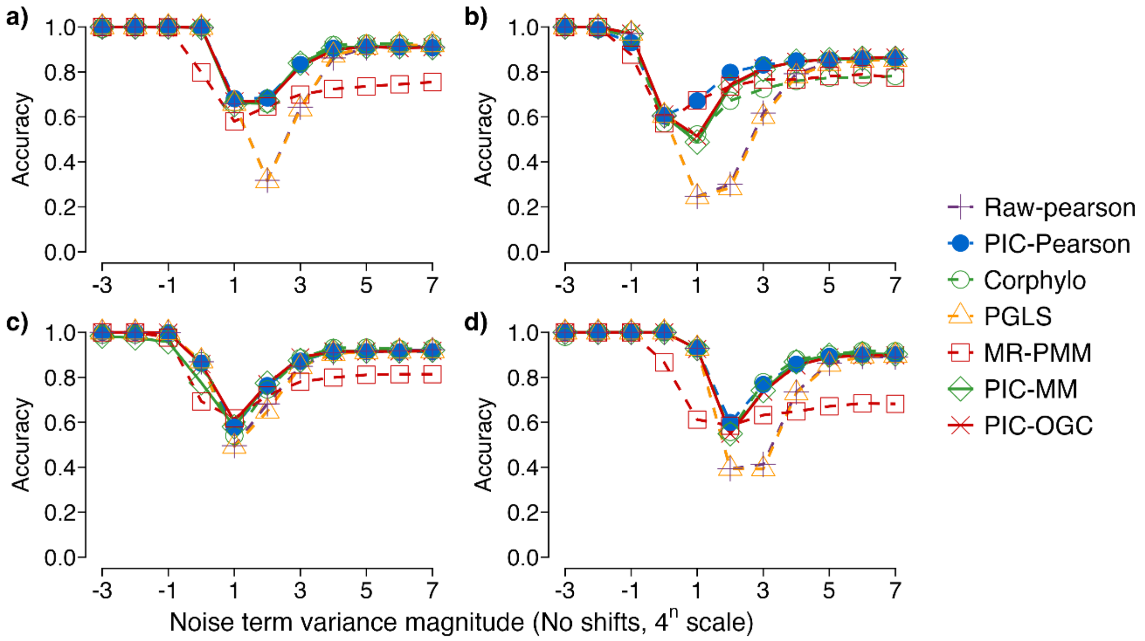

Supplementary Figure S10. Accuracies of seven methods under the BM + Norm scenario evaluated on four phylogenetic settings using the data-pattern–based benchmark.

**a)** a fixed-balanced tree with 128 species; **b)** an ensemble of random trees with 128 species; **c)** a fixed-balanced tree with 16 species; and **d)** a fixed-balanced tree with 256 species. The horizontal axis shows the variance of the added noise term as $4^n$; only the exponent n is printed (e.g. $n$=−3 represents $4^{-3}$, and $n$=7 represents $4^7$). "Raw-Pearson" refers to applying the standard Pearson correlation directly to the original species-level trait data, without any phylogenetic transformation. Additional details on the simulation design, gradient configurations, and performance metrics are provided in the Materials and Methods. The data used to plot this figure are available in Supplementary Table S18.

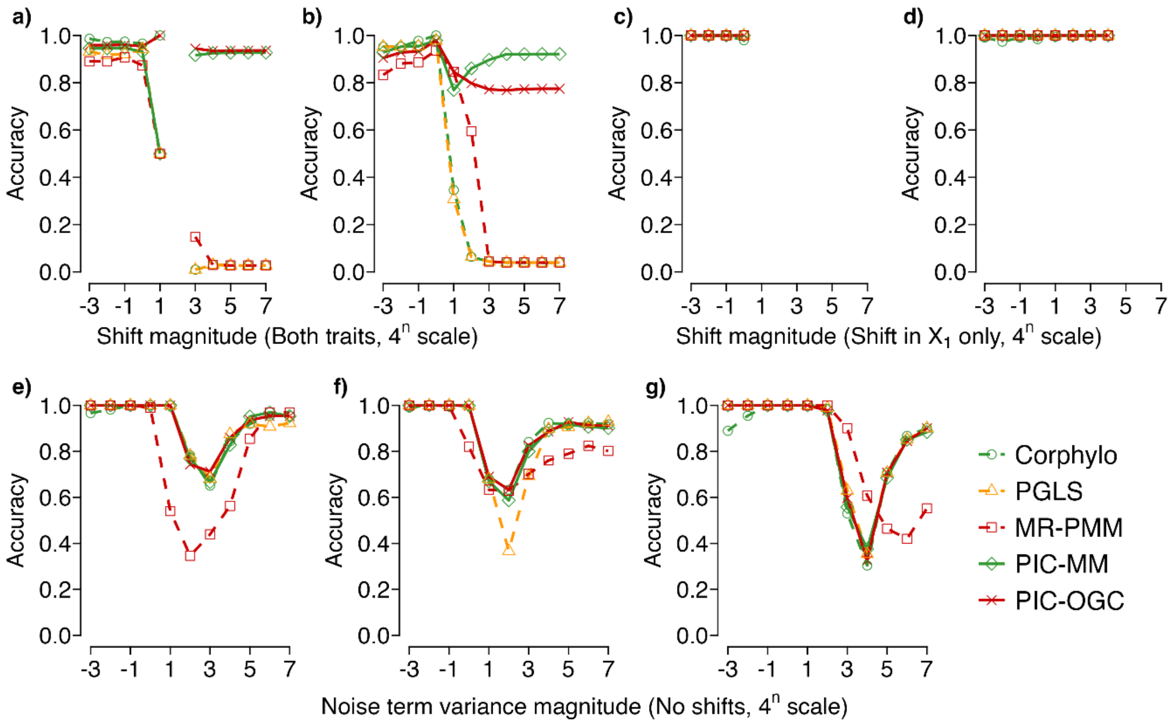

Supplementary Figure S11. Performance of five methods on datasets that deviate from Brownian motion (BM) assumptions on a 128-species fixed-balanced tree using the data-pattern–based benchmark.

Panels **a–b** show performance under non-BM datasets with evolutionary shifts in both $X_1$ and $X_2$, where shifts are introduced at the root branch (Felsenstein's worst-case scenario; **a**) or at randomly selected branch locations (**b**). Panels **c–d** show performance under non-BM datasets with evolutionary shifts only in $X_1$, again comparing worst-case root shifts (**c**) and randomly located shifts (**d**). Panels **e–g** show performance under non-BM datasets without abrupt evolutionary shifts, across three gradual-evolution scenarios: BM–BM (panel **e**), BM–Norm (panel **f**), and Norm–BM (panel **g**). Shift and noise variance values are represented as $4^n$, with the horizontal axis showing only the exponent *n* (e.g., n = −3 corresponds to $4^{-3}$, and n = 7 corresponds to $4^7$). All datasets were simulated on a fixed-balanced phylogenetic tree with 128 species. See the Materials and Methods for additional details on the simulation design, gradient configurations, and performance metrics. The data used to generate this figure are provided in Supplementary Table S19**.**

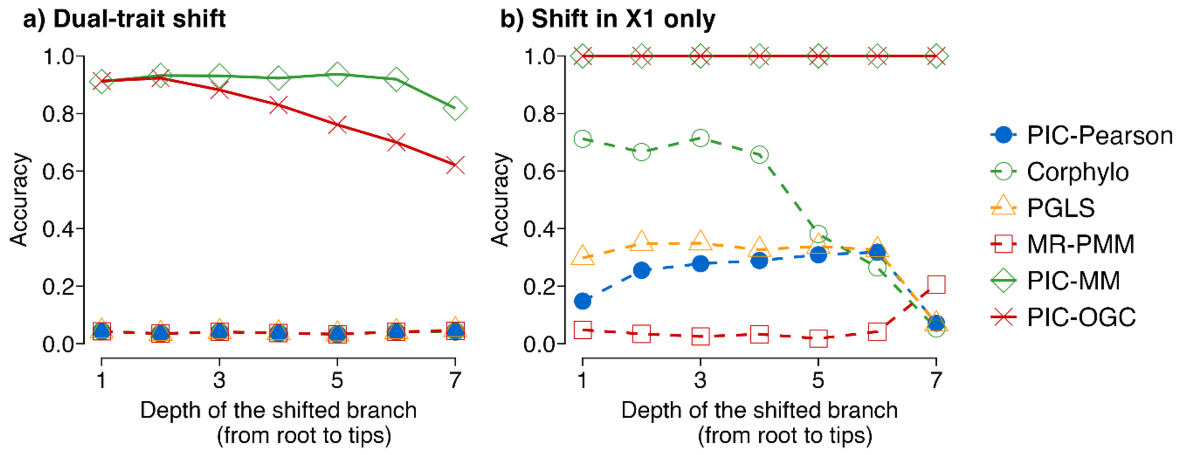

Supplementary Figure S12. Sensitivity of method performance to the phylogenetic location of evolutionary shifts using the data-pattern–based benchmark.

Performance of six methods evaluated under shifts introduced at different depths of a 128-species fixed-balanced tree. A representative shift magnitude of $4^5$ was applied systematically at each phylogenetic depth. Panel a shows false positive rates under simultaneous shifts in both traits, whereas panel b shows false negative rates under univariate shifts affecting only $X_1$. False positive and false negative rates are reported as proportions. Additional details of the simulation design and performance metrics are provided in the Materials and Methods. The data used to generate this figure are provided in Supplementary Table S20**.**
