## Supplementary methods. for "Improving the Robustness of Phylogenetic Independent Contrasts: Addressing Abrupt Evolutionary Shifts with Outlier-and Distribution-Guided Correlation"

### *Supplementary Methods S1*

### *Mathematical and Statistical Foundations of the Second Benchmarking Framework*

#### S1.1 Overview

This supplementary section provides the mathematical and statistical foundations underlying the benchmarking frameworks described in the main Methods. Specifically, we (i) formalize the population-level correlation implied by the generative model used in the simulations, $X_2 = \beta X_1 + \varepsilon$, (ii) clarify how stochastic variance shapes the expected strength of association under this model, and (iii) explain why benchmarks based on branch-level evolutionary changes provide an informative and statistically valid reference for evaluating correlation inference methods.

Together, these derivations establish the conceptual distinction between structural relationships encoded by the coupling parameter $\beta$ in the generative model and associations that are statistically supported by realized data, thereby motivating the complementary benchmarking strategies adopted in this study.

#### S1.2 Generative model

Across all simulation settings, the dependent trait was generated according to:

$$X_2 = \beta X_1 + \varepsilon, \quad X_1 \perp \varepsilon,$$

where $X_1$ represents the value of the predictor trait and $\varepsilon$ is an independent stochastic term.

Depending on the simulation scenario, $\varepsilon$ represents either normally distributed noise or an independent Brownian motion (BM) process evolving on the same phylogeny as $X_1$. Independence ensures that

$$Cov(X_1, \varepsilon) = 0.$$

Let:

$$Var(X_1) = \sigma_{X_1}^2, \quad Var(\varepsilon) = \sigma_\varepsilon^2.$$

These expressions denote the marginal variances at the tips; for BM models, these variances implicitly incorporate the shared-branch covariance structure induced by the phylogeny.

#### S1.3 Covariance relationships

By the generative model:

$$Cov(X_1, X_2) = Cov(X_1, \beta X_1 + \varepsilon) = \beta Var(X_1) = \beta \sigma_{X_1}^2.$$

Similarly:

$$Var(X_2) = \beta^2 \sigma_{X_1}^2 + \sigma_\varepsilon^2.$$

For BM models evolving on a phylogeny, these identities continue to hold because independent BM processes accumulate variance additively along branches, and independence ensures the absence of cross-covariance terms between $X_1$ and $\varepsilon$.

#### S1.4 Population-level correlation under the generative model

Under the generative model above, the population-level correlation between $X_1$ and $X_2$ is given by:

$$\rho = Corr(X_1, X_2) = \frac{\beta\sigma_{X_1}^2}{\sqrt{\sigma_{X_1}^2(\beta^2\sigma_{X_1}^2 + \sigma_\varepsilon^2)}} = \frac{\beta}{\sqrt{\beta^2 + \sigma_\varepsilon^2/\sigma_{X_1}^2}}.$$

Define the noise-to-signal ratio:

$$\eta = \frac{\sigma_\varepsilon^2}{\sigma_{X_1}^2}.$$

Then

$$\rho = \frac{\beta}{\sqrt{\beta^2 + \eta}}.$$

This equation gives the *true population correlation* in the classical statistical sense: the correlation that would be obtained in the limit of infinitely many replicates generated under the same model.

#### S1.5 Dependence of population-level correlation on stochastic variance

In all simulations, we set $Var(X_1) = 1$, which implies that the noise-to-signal variance ratio

$$\eta = \frac{\sigma_\varepsilon^2}{\sigma_{X_1}^2}$$

simplifies to

$$\eta = \sigma_\varepsilon^2.$$

Thus, population-level correlation follows directly from

$$\rho = \frac{\beta}{\sqrt{\beta^2 + \sigma_\varepsilon^2}}.$$

Under the assumptions used in most simulations—specifically, $\beta = 1$ and $\mathrm{Var}(X_1) = 1$—we computed the population-level correlation $\rho$ corresponding to each value of the noise variance $\sigma_\varepsilon^2$ examined in the study. The resulting values of $\rho$ are summarized in Table AS1, which reports the expected population-level correlation implied by the generative model ($X_2 = \beta X_1 + \varepsilon$) across the full range of stochastic variance regimes considered.

As shown in Table AS1, population-level correlation remains high when noise variance is small but declines steeply as $\sigma_\varepsilon^2$ increases. Consequently, even when the generative model structurally specifies a dependence between traits (e.g., $X_2 = \beta X_1 + \varepsilon$), the expected population correlation can become arbitrarily weak under sufficiently large stochastic variance. This continuum of correlation strength defines the strong-signal and weak-signal regimes that form the basis for the benchmarking analyses reported in the main text.

**Table AS1. Population-level correlation ($\rho$) implied by the generative model under $\beta$ = 1 across the noise-variance regimes used in the simulations.**

| Noise variance | Population-level correlation $\rho$ |
|---|---|
| $4^{-3}$ = 0.015625 | 0.992 |
| $4^{-2}$ = 0.0625 | 0.970 |
| $4^{-1}$ = 0.25 | 0.894 |
| $4^{-0}$ = 1 | 0.707 |
| 3 (*reference*) | 0.500 |
| $4^1$ = 4 | 0.447 |
| $4^2$ = 16 | 0.243 |
| $4^3$ = 64 | 0.124 |
| $4^4$ = 256 | 0.062 |
| $4^5$ = 1024 | 0.031 |
| $4^6$ = 4096 | 0.016 |
| $4^7$ = 16384 | 0.008 |

The table reports the expected population-level correlation corresponding to different values of the noise variance $\sigma_\varepsilon^2$, spanning a wide range from $4^{-3}$ to $4^7$. These values define the full gradient of stochastic intensities examined in the study, from strong-signal to extremely weak-signal regimes. For reference, one intermediate variance value ($\sigma_\varepsilon^2$ = 3) is included because it yields $\rho$ = 0.50, which serves as a reference value representing a moderate expected correlation and helps illustrate the progressive weakening of association as $\sigma_\varepsilon^2$ increases.

#### S1.6 Benchmark label definition and performance metrics

Although the noise variance uniquely determines the expected population-level correlation under the generative model (Table AS1), it does not uniquely determine whether a particular realized dataset provides statistical support for association. Datasets generated under the same noise regime may or may not exhibit significant correlation due to finite sample size and stochastic variation. Consequently, benchmark labels cannot be inferred directly from noise variance alone and must instead be defined at the replicate level based on realized data patterns.

For each simulation replicate, evidence for association was evaluated using correlation tests applied to the normalized branch-level trait changes ($\Delta X_1/L$ and $\Delta X_2/L$). Because these normalized branch-level changes—like phylogenetically independent contrasts (PICs)—can exhibit extreme outliers or non-normal distributions under strong evolutionary shifts, a data-adaptive correlation strategy was adopted to define benchmark labels. Specifically, Pearson or Spearman correlation was selected based on the presence of outliers, following criteria analogous to those used in the PIC-O(D)GC workflow. For smaller datasets (16 species), additional normality testing was applied to inform the choice of correlation statistic.

Replicates exhibiting statistically significant correlation ($p < 0.05$) under this adaptive procedure were classified as benchmark-positive, whereas replicates lacking significant evidence were classified as benchmark-negative. Method performance under this benchmark was quantified using classification accuracy, defined as the proportion of replicates for which inferred results agreed with these data-driven benchmark labels.

#### S1.7 Conceptual implications for benchmarking

The derivations above clarify why structural dependence encoded by the coupling parameter $\beta$ does not directly translate into statistical support for association in realized datasets. Although the coupling parameter ($\beta$) in the generative model may specify a deterministic relationship between traits, stochastic evolutionary variation ($\varepsilon$) can substantially weaken the corresponding statistical signal observed in finite datasets. The concept of population-level correlation provides a useful intermediate reference for understanding this attenuation, but does not itself determine whether a particular dataset exhibits statistically supported association.

Conversely, when association is supported by the realized data, it represents a legitimate statistical pattern that inference methods should aim to reflect, irrespective of whether the underlying evolutionary process is deterministic or stochastic (Ives 2022).

#### Reference

Ives A.R. 2022. Random errors are neither: On the interpretation of correlated data. Methods Ecol. Evol., 13:2092-2105.
