## Supplementary material for "Improving the Robustness of Phylogenetic Independent Contrasts: Addressing Abrupt Evolutionary Shifts with Outlier-and Distribution-Guided Correlation": Codes used in this study.: 01Simulation_RandomTree.R

```
library(ape)
library(phytools)
args <- commandArgs()
simulate.seed <- args[6]

creat_random_tree <- function(n1,n2){

  root_tr<-rcoal(2)
  sub_tr1 <- pbtree(n = n1, scale = 1) 
  sub_tr2<- sub_tr1
  
  root_tr$tip.label[1]<-"NA"
  sub_tr1$root.edge<-0
  
  bind_tr1<-paste.tree(root_tr,sub_tr1)
  
  bind_tr1$tip.label[1]<-"NA"
  sub_tr2$root.edge<-0
  
  final_tr<-paste.tree(bind_tr1,sub_tr2)
  
  return(final_tr)
}
```
