## Supplementary material for "Improving the Robustness of Phylogenetic Independent Contrasts: Addressing Abrupt Evolutionary Shifts with Outlier-and Distribution-Guided Correlation": Codes used in this study.: 02Simulation_BM1&BM1+BM2.R

```
library(ape)
library(geiger)
library(ggplot2)
library(phytools)
args <- commandArgs()
simulate.seed <- args[6]
#simulate.seed <- 1
#set.seed(1)
#set.seed(123)
## Using phylogenetic information to impute missing functional trait values in ecological databases
# Main functions

# Function to data simulation, imputation by missForest and to get the NRMSE
# parameters = A data.frame with only one row and eight columns, the names of columns are all parameters setting to run the simulation
PAR1 <- expand.grid(diff_sd = c(sqrt(4^-5),sqrt(4^-4),sqrt(4^-3),sqrt(4^-2),sqrt(4^-1),sqrt(1),
                                sqrt(4),sqrt(16),sqrt(64),sqrt(256),sqrt(1024),sqrt(4096),sqrt(16384)),
                    n1.spp = c(8), 
                    n2.spp = c(8),
                    n.spp = c(16), # Number of species in the phylogenetic tree and in the dataset
                    feature_num = 2) 

get.simulate.df <- function(parameters){
  tips_num <- parameters$n.spp
  spp_n1 <- parameters$n1.spp
  spp_n2 <- parameters$n2.spp
  
  feature_num <- parameters$feature_num
  diff_sd <- parameters$diff_sd
  all_state_df <- NULL
  tips_state_df <- NULL
  nodes_state_df <- NULL
  
  my_tree <- creat_tree(spp_n1,spp_n2)
  set.seed(simulate.seed)
  my_tree$tip.label <- c(1:length(my_tree$tip.label))
  tips_label <- my_tree$tip.label
  N <- my_tree$Nnode
  nodes_names <- c((tips_num+1):(tips_num+N))
  my_tree$node.label <- nodes_names  ###修改nodes名字
  ID <- names(rTraitCont(my_tree,"BM",ancestor = T,root.value = 0,sigma = 1))
  my_feature_df <- data.frame('ID'=ID)
  
  return_tree <- my_tree
  rescale_tree <- my_tree
  #rescale_tree <- rescale(my_tree,"lambda",1)
  X1 <- rTraitCont(rescale_tree,"BM",ancestor = T,root.value = 1,sigma = 1)
  X2 <- rTraitCont(rescale_tree,"BM",ancestor = T,root.value = 1,sigma = diff_sd)
  for (i in c(1:feature_num)){
    feature_name <- paste("X",i,sep="")
    my_feature_df[feature_name] <- (2-i)*X1 + 1*(i-1)*X1 + 1*(i-1)*X2
  }
  
  tips_df <- my_feature_df[my_feature_df$ID %in% tips_label,]
  return(list(return_tree,tips_df,my_feature_df))
}
index_num <- length(rownames(PAR1))

for (index in c(1:index_num)){
  param_vec <- PAR1[index,]
  return_list <- get.simulate.df(param_vec)
  my_tree <- return_list[[1]]
  tips_df <- return_list[[2]]
  final_all_df <- return_list[[3]]
  command <- paste0("mkdir ./results/",index)
  system(command = command)
  tips_df_out_file <- paste0("results/",index,"/tips_states.csv")
  tree_out_file <- paste0("results/",index,"/mytree.nwk")
  rela_out_file <- paste0("results/",index,"/Parent2Child.txt")
  all_df_out_file <- paste0("results/",index,"/all_states.csv")
  write.tree(my_tree,tree_out_file)
  write.csv(tips_df,tips_df_out_file,quote=F,row.names = F)
  write.csv(PAR1,"results/parameters.csv",quote=F)
  edge_res <- my_tree$edge
  Parent <- edge_res[,1]
  Child <- edge_res[,2]
  relation_df <- data.frame(Parent,Child)
  write.table(relation_df,file = rela_out_file,quote = FALSE,row.names = FALSE,
              col.names = T)
  write.csv(final_all_df,file = all_df_out_file,quote=F,row.names = F)
}
```
