## Supplementary material for "Improving the Robustness of Phylogenetic Independent Contrasts: Addressing Abrupt Evolutionary Shifts with Outlier-and Distribution-Guided Correlation": Codes used in this study.: 02Simulation_ShiftsinBothTraits_fixedlocation.R

```
library(ape)
library(phytools)
args <- commandArgs()
simulate.seed <- args[6]
#simulate.seed <- 1

PAR1 <- expand.grid(shift = c(4^-5,4^-4,4^-3,(4^-2),(4^-1),(1),
                              (4),(16),(64),(256),(1024),(4096),(16384)),# overall BM model variance
                    n1.spp = c(8), 
                    n2.spp = c(8),
                    n.spp = c(16),# Number of species in the phylogenetic tree and in the dataset
                    feature_num = 2) 


get.simulate.df <- function(parameters){
  tips_num <- parameters$n.spp
  spp_n1 <- parameters$n1.spp
  spp_n2 <- parameters$n2.spp
  
  feature_num <- parameters$feature_num
  shift <- parameters$shift
  all_state_df <- NULL
  tips_state_df <- NULL
  nodes_state_df <- NULL
  
  
  my_tree <- creat_tree(spp_n1,spp_n2) # creat_balanced_tree() or creat_random_tree()
  set.seed(simulate.seed)
  my_tree$tip.label <- c(1:length(my_tree$tip.label))
  tips_label <- my_tree$tip.label
  N <- my_tree$Nnode
  nodes_names <- c((tips_num+1):(tips_num+N))
  my_tree$node.label <- nodes_names  ### rename the Node  
  ## find ROOT and corresponding SON nodes
  root_node <- my_tree$edge[1,1]
  son_nodes <- my_tree$edge[c(my_tree$edge[,1]==root_node),][,2]
  
  ## pickup a shift node 
  modify_node <- son_nodes[1]
  modify_tree <- extract.clade(my_tree, node = modify_node)
  modify_node
  
  overall_X1 <- rTraitCont(my_tree,"BM",ancestor = T,root.value = 1,sigma = 1) 
  overall_X2 <- rTraitCont(my_tree,"BM",ancestor = T,root.value = 1,sigma = 1)
  
  ID <- names(overall_X1)
  my_feature_df <- data.frame('ID'=ID)
  
  ## modify the SON node status to 10 times of ROOT
  modify_X1 <- rTraitCont(modify_tree,"BM",ancestor = T,root.value = 1*shift,sigma = 1)
  modify_X2 <- rTraitCont(modify_tree,"BM",ancestor = T,root.value = 1*shift,sigma = 1)
  
  final_X1 <- overall_X1
  final_X2 <- overall_X2
  
  final_X1[names(modify_X1)] <- modify_X1
  final_X2[names(modify_X2)] <- modify_X2
  
  my_feature_df[,"X1"] <- final_X1
  my_feature_df[,"X2"] <- final_X2
  
  tips_df <- my_feature_df[my_feature_df$ID %in% tips_label,]
  return(list(my_tree,tips_df,my_feature_df))
}


index_num <- length(rownames(PAR1))

for (index in c(1:index_num)){
  param_vec <- PAR1[index,]
  return_list <- get.simulate.df(param_vec)
  my_tree <- return_list[[1]]
  tips_df <- return_list[[2]]
  final_all_df <- return_list[[3]]
  command <- paste0("mkdir ./results/",index)
  system(command = command)
  tips_df_out_file <- paste0("results/",index,"/tips_states.csv")
  tree_out_file <- paste0("results/",index,"/mytree.nwk")
  rela_out_file <- paste0("results/",index,"/Parent2Child.txt")
  all_df_out_file <- paste0("results/",index,"/all_states.csv")
  write.tree(my_tree,tree_out_file)
  write.csv(tips_df,tips_df_out_file,quote=F,row.names = F)
  write.csv(PAR1,"results/parameters.csv",quote=F)
  edge_res <- my_tree$edge
  Parent <- edge_res[,1]
  Child <- edge_res[,2]
  relation_df <- data.frame(Parent,Child)
  write.table(relation_df,file = rela_out_file,quote = FALSE,row.names = FALSE,
              col.names = T)
  write.csv(final_all_df,file = all_df_out_file,quote=F,row.names = F)
}
```
