## Supplementary material for "Improving the Robustness of Phylogenetic Independent Contrasts: Addressing Abrupt Evolutionary Shifts with Outlier-and Distribution-Guided Correlation": Codes used in this study.: 02Simulation_ShiftsinOneTrait_fixedlocation.R

```
library(ape)
library(phytools)
args <- commandArgs()
simulate.seed <- args[6]
#simulate.seed <- 1

PAR1 <- expand.grid(shift = c(4^-5,4^-4,4^-3,(4^-2),(4^-1),(1),
                              (4),(16),(64),(256),(1024),(4096),(16384)),# overall BM model variance
                    n1.spp = c(8), 
                    n2.spp = c(8),
                    n.spp = c(16),# Number of species in the phylogenetic tree and in the dataset
                    feature_num = 2) 

get.simulate.df <- function(parameters,seed){
  tips_num <- parameters$n.spp
  spp_n1 <- parameters$n1.spp
  spp_n2 <- parameters$n2.spp
  
  feature_num <- parameters$feature_num
  shift <- parameters$shift
  all_state_df <- NULL
  tips_state_df <- NULL
  nodes_state_df <- NULL
  
  seed1 <- as.numeric(seed)
  seed2 <- seed1 + 1
  
  my_tree <- creat_tree(spp_n1,spp_n2)
  my_tree$tip.label <- c(1:length(my_tree$tip.label))
  tips_label <- my_tree$tip.label
  N <- my_tree$Nnode
  nodes_names <- c((tips_num+1):(tips_num+N))
  my_tree$node.label <- nodes_names  ### rename the Node  
  ## find ROOT and corresponding SON nodes
  root_node <- my_tree$edge[1,1]
  son_nodes <- my_tree$edge[c(my_tree$edge[,1]==root_node),][,2]
  
  ## pickup a shift node 
  modify_node <- son_nodes[1]
  modify_tree <- extract.clade(my_tree, node = modify_node)
  modify_node
  
  ## pickup a remain node
  remain_node <- son_nodes[2]
  remain_tree <- extract.clade(my_tree, node = remain_node)
  remain_node
  
  ## modify the modify_tree data ###
  set.seed(seed1)
  modify_X1 <- rTraitCont(modify_tree,"BM",ancestor = T,root.value = 1*shift,sigma = 1)
  set.seed(seed1)
  original_X2 <- rTraitCont(modify_tree,"BM",ancestor = T,root.value = 1,sigma = 1)
  e <- rnorm(length(original_X2),1,0.1)
  modify_X2 <- original_X2 + e
  
  ## simulate the all_tree data for the remain_tree's root ###
  set.seed(seed1)
  try_X1 <- rTraitCont(my_tree,"BM",ancestor = T,root.value = 1,sigma = 1)

  remain_root_X1 <- try_X1[names(try_X1)==remain_node]
  
  
  ## simulate the remain_tree data ###
  set.seed(seed2)
  remain_X1 <- rTraitCont(remain_tree,"BM",ancestor = T,root.value = remain_root_X1,sigma = 1)
  e <- rnorm(length(original_X2),1,0.1)
  remain_X2 <- remain_X1 + e
  
  
  overall_X1 <- c(modify_X1,try_X1[names(try_X1)==root_node],remain_X1)
  overall_X2 <- c(modify_X2,try_X1[names(try_X1)==root_node],remain_X2)
  
  reordered_X1 <- overall_X1[order(as.numeric(names(overall_X1)))]
  reordered_X2 <- overall_X2[order(as.numeric(names(overall_X2)))]
  
  my_feature_df <- as.data.frame(cbind(names(reordered_X1),
                                       as.numeric(reordered_X1),
                                       as.numeric(reordered_X2)))
  colnames(my_feature_df) <- c("ID","X1","X2")
  
  my_feature_df$X1 <- as.numeric(my_feature_df$X1)
  my_feature_df$X2 <- as.numeric(my_feature_df$X2)
  
  tips_df <- my_feature_df[my_feature_df$ID %in% tips_label,]
  #tips_df[,"label"] <- c(rep("1",64),rep("2",64))
  return(list(my_tree,tips_df,my_feature_df))
}

index_num <- length(rownames(PAR1))

for (index in c(1:index_num)){
  param_vec <- PAR1[index,]
  return_list <- get.simulate.df(param_vec,simulate.seed)
  my_tree <- return_list[[1]]
  tips_df <- return_list[[2]]
  final_all_df <- return_list[[3]]
  command <- paste0("mkdir ./results1/",index)
  system(command = command)
  tips_df_out_file <- paste0("results1/",index,"/tips_states.csv")
  tree_out_file <- paste0("results1/",index,"/mytree.nwk")
  rela_out_file <- paste0("results1/",index,"/Parent2Child.txt")
  all_df_out_file <- paste0("results1/",index,"/all_states.csv")
  write.tree(my_tree,tree_out_file)
  write.csv(tips_df,tips_df_out_file,quote=F,row.names = F)
  write.csv(PAR1,"results1/parameters.csv",quote=F)
  edge_res <- my_tree$edge
  Parent <- edge_res[,1]
  Child <- edge_res[,2]
  relation_df <- data.frame(Parent,Child)
  write.table(relation_df,file = rela_out_file,quote = FALSE,row.names = FALSE,
              col.names = T)
  write.csv(final_all_df,file = all_df_out_file,quote=F,row.names = F)
}
```
