## Supplementary material for "Improving the Robustness of Phylogenetic Independent Contrasts: Addressing Abrupt Evolutionary Shifts with Outlier-and Distribution-Guided Correlation": Codes used in this study.: 02Simulation_ShiftsinOneTrait_randomlocation.R

```
library(ape)
library(phytools)
args <- commandArgs()
simulate.seed <- args[6]
#simulate.seed <- 1

PAR1 <- expand.grid(shift = c(4^-5,4^-4,4^-3,(4^-2),(4^-1),(1),
                              (4),(16),(64),(256),(1024),(4096),(16384)),# overall BM model variance
                    n1.spp = c(8), 
                    n2.spp = c(8),
                    n.spp = c(16),# Number of species in the phylogenetic tree and in the dataset
                    feature_num = 2) 

get.simulate.df <- function(parameters,seed){
  tips_num <- parameters$n.spp
  spp_n1 <- parameters$n1.spp
  spp_n2 <- parameters$n2.spp
  
  feature_num <- parameters$feature_num
  shift <- parameters$shift
  all_state_df <- NULL
  tips_state_df <- NULL
  nodes_state_df <- NULL
  
  seed1 <- as.numeric(seed)
  seed2 <- seed1 + 1
  
  my_tree <- creat_tree(spp_n1,spp_n2)
  my_tree$tip.label <- c(1:length(my_tree$tip.label))
  tips_label <- my_tree$tip.label
  N <- my_tree$Nnode
  nodes_names <- c((tips_num+1):(tips_num+N))
  my_tree$node.label <- nodes_names  ### rename the Node  
  ## find ROOT and corresponding SON nodes
  root_node <- sample(my_tree$edge[,1],1)
  son_nodes <- my_tree$edge[c(my_tree$edge[,1]==root_node),][,2]
  
  ## pickup a shift node 
  modify_node <- son_nodes[1]
  remain_node <- son_nodes[2]
  
  if (modify_node %in% my_tree$edge[,1]){
    modify_tree <- extract.clade(my_tree, node = modify_node)
    ## pickup a remain node
    remain_tree <- extract.clade(my_tree, node = remain_node)
    remain_node
    
    #### all_raw traits simualtion ### 
    set.seed(seed1)
    all_raw_X1 <- rTraitCont(my_tree,"BM",ancestor = T,root.value = 1,sigma = 1)
    set.seed(seed1)
    e <- rnorm(length(all_raw_X1),1,0.1)
    all_raw_X2 <- all_raw_X1 + e
    
    #### re-simulate X1 ####
    set.seed(seed1)
    modify_node_X1 <- all_raw_X1[modify_node]
    modify_node_X2 <- all_raw_X2[modify_node]
    
    modify_X1 <- rTraitCont(modify_tree,"BM",ancestor = T,root.value = modify_node_X1*shift,sigma = 1)
    
    set.seed(seed1)
    modify_original_X1 <- rTraitCont(modify_tree,"BM",ancestor = T,root.value = modify_node_X2,sigma = 1)
    
    modify_e <- rnorm(length(modify_original_X1),1,0.1)
    modify_X2 <- modify_original_X1 + modify_e
    
    final_X1 <- all_raw_X1
    final_X2 <- all_raw_X2
    
    final_X1[names(final_X1) %in% names(modify_X1)] <- modify_X1
    final_X2[names(final_X2) %in% names(modify_X2)] <- modify_X2
    
    
    my_feature_df <- as.data.frame(cbind(names(final_X1),
                                         as.numeric(final_X1),
                                         as.numeric(final_X2)))
    colnames(my_feature_df) <- c("ID","X1","X2")
    
    my_feature_df$X1 <- as.numeric(my_feature_df$X1)
    my_feature_df$X2 <- as.numeric(my_feature_df$X2)
    
    tips_df <- my_feature_df[my_feature_df$ID %in% tips_label,]
  }else{
    set.seed(seed1)
    raw_X1 <- rTraitCont(my_tree,"BM",ancestor = T,root.value = 1,sigma = 1)
    set.seed(seed1)
    raw_X2 <- rTraitCont(my_tree,"BM",ancestor = T,root.value = 1,sigma = 1)
    e <- rnorm(length(raw_X2),1,0.1)
    final_X2 <- raw_X2 + e
    
    final_X1 <- raw_X1
    final_X1[names(final_X1)==modify_node] <- final_X1[names(final_X1)==modify_node] * shift
    
    my_feature_df <- as.data.frame(cbind(names(final_X1),
                                         as.numeric(final_X1),
                                         as.numeric(final_X2)))
    colnames(my_feature_df) <- c("ID","X1","X2")
    
    my_feature_df$X1 <- as.numeric(my_feature_df$X1)
    my_feature_df$X2 <- as.numeric(my_feature_df$X2)
    
    tips_df <- my_feature_df[my_feature_df$ID %in% tips_label,]
  }
  
  #tips_df[,"label"] <- c(rep("1",64),rep("2",64))
  return(list(my_tree,tips_df,my_feature_df))
}

index_num <- length(rownames(PAR1))

for (index in c(1:index_num)){
  param_vec <- PAR1[index,]
  return_list <- get.simulate.df(param_vec,simulate.seed)
  my_tree <- return_list[[1]]
  tips_df <- return_list[[2]]
  final_all_df <- return_list[[3]]
  command <- paste0("mkdir ./results31/",index)
  system(command = command)
  tips_df_out_file <- paste0("results31/",index,"/tips_states.csv")
  tree_out_file <- paste0("results31/",index,"/mytree.nwk")
  rela_out_file <- paste0("results31/",index,"/Parent2Child.txt")
  all_df_out_file <- paste0("results31/",index,"/all_states.csv")
  write.tree(my_tree,tree_out_file)
  write.csv(tips_df,tips_df_out_file,quote=F,row.names = F)
  write.csv(PAR1,"results31/parameters.csv",quote=F)
  edge_res <- my_tree$edge
  Parent <- edge_res[,1]
  Child <- edge_res[,2]
  relation_df <- data.frame(Parent,Child)
  write.table(relation_df,file = rela_out_file,quote = FALSE,row.names = FALSE,
              col.names = T)
  write.csv(final_all_df,file = all_df_out_file,quote=F,row.names = F)
}
```
