## Supplementary material for "Improving the Robustness of Phylogenetic Independent Contrasts: Addressing Abrupt Evolutionary Shifts with Outlier-and Distribution-Guided Correlation": Codes used in this study.: 03Analysis_01_PIC.R

```
command <- paste0("ls data | wc -l")
system(command)
args <- commandArgs(trailingOnly = TRUE)
library(ape)


get_history_change <- function(all_df,relation_df,X){
  L <- length(rownames(relation_df))
  res_vec <- NULL
  for(i in c(1:L)){
    parent <- relation_df$Parent[i]
    child <- relation_df$Child[i]
    parent_trait_X <- all_df[all_df$ID==parent,X]
    child_trait_X <- all_df[all_df$ID==child,X]
    change_X <- (child_trait_X - parent_trait_X)/relation_df$distance[i]
    res_vec <- c(res_vec,change_X)
  }  
  return(res_vec)
}

index_num <- 13

t_pearson_est_vec <- NULL
t_pearson_p_vec <- NULL
t_spearman_est_vec <- NULL
t_spearman_p_vec <- NULL
t_cor_est_vec <- NULL
t_cor_p_vec <- NULL


PIC_spearman_est_vec <- NULL
PIC_spearman_p_vec <- NULL
PIC_pearson_est_vec <- NULL
PIC_pearson_p_vec <- NULL
PIC_est_vec <- NULL
PIC_p_vec <- NULL

final_est_vec <- NULL
final_p_vec <- NULL


for (index in c(1:index_num)){
  sig_label <- "O"
  tips_df_file <- paste0("data/",index,"/tips_states.csv")
  all_df_file <- paste0("data/",index,"/all_states.csv")
  relation_file <- paste0("data/",index,"/Parent2Child.txt")
  my_tree_file <- paste0("data/",index,"/mytree.nwk")
  relation_df <- read.table(relation_file,sep = " ",header = T)
  all_df <- read.csv(all_df_file)
  tips_df <- read.csv(tips_df_file)
  my_tree <- read.tree(my_tree_file)
  
  tips_num <- length(tips_df$ID)
  relation_df$distance <- my_tree$edge.length
  final_est <- NULL
  final_p <- NULL
  
  
  ## golden standard ###
  X1_change <- get_history_change(all_df,relation_df,"X1")
  X2_change <- get_history_change(all_df,relation_df,"X2")
  
  change_formula <- as.formula(paste0("X1_change~X2_change"))
  change_est <- summary(lm(change_formula))$coefficients[2,1]
  change_p <- summary(lm(change_formula))$coefficients[2,4]
  spearman_est <- cor.test(X1_change,X2_change,method="spearman")$estimate
  spearman_p <- cor.test(X1_change,X2_change,method="spearman")["p.value"]$p.value
  
  ## normality test ###
  X1_normal_p <- shapiro.test(X1_change)$p.value
  X2_normal_p <- shapiro.test(X2_change)$p.value
  
  if (X1_normal_p<0.05 | X2_normal_p<0.05){
    final_est <- spearman_est
    final_p <- spearman_p
  }else{
    final_est <- change_est
    final_p <- change_p
  }
  
  final_est_vec <- c(final_est_vec,final_est)
  final_p_vec <- c(final_p_vec,final_p)
  
  
  ### Traits pearson ##
  X1 <- tips_df[,"X1"]
  X2 <- tips_df[,"X2"]
  t_pearson_est <- cor.test(X1,X2,method="pearson")$estimate
  t_pearson_p <- cor.test(X1,X2,method="pearson")$p.value
  t_spearman_est <- cor.test(X1,X2,method="spearman")$estimate
  t_spearman_p <- cor.test(X1,X2,method="spearman")$p.value
  X1_normal_p <- shapiro.test(X1)$p.value
  X2_normal_p <- shapiro.test(X2)$p.value
  if (X1_normal_p<0.05 | X2_normal_p<0.05){
    t_est <- t_spearman_est
    t_p <- t_spearman_p
  }else{
    t_est <- t_pearson_est
    t_p <- t_pearson_p
  }
  
  
  
  ## PIC Pearson ### 
  X1 <- tips_df[,"X1"]
  X2 <- tips_df[,"X2"]
  PIC_X1 <- pic(X1,my_tree)
  PIC_X2 <- pic(X2,my_tree)
  
  PIC_formula <- as.formula(paste0("PIC_X1~PIC_X2"))
  pearson_est <- summary(lm(PIC_X2~PIC_X1-1))$coefficients[1,1]
  pearson_p <- summary(lm(PIC_X2~PIC_X1-1))$coefficients[1,4]
  spearman_est <- cor.test(PIC_X1,PIC_X2,method="spearman")$estimate
  spearman_p <- cor.test(PIC_X1,PIC_X2,method="spearman")["p.value"]$p.value
  
  ## normality test ###
  PIC_X1_normal_p <- shapiro.test(PIC_X1)$p.value
  PIC_X2_normal_p <- shapiro.test(PIC_X2)$p.value
  if (PIC_X1_normal_p<0.05 | PIC_X2_normal_p<0.05){
    PIC_est <- spearman_est
    PIC_p <- spearman_p
  }else{
    PIC_est <- pearson_est
    PIC_p <- pearson_p
  }
  
  PIC_spearman_est <- spearman_est
  PIC_spearman_p <- spearman_p
  PIC_pearson_est <- pearson_est
  PIC_pearson_p <- pearson_p
  
  
  t_pearson_est_vec <- c(t_pearson_est_vec,t_pearson_est)
  t_pearson_p_vec <- c(t_pearson_p_vec,t_pearson_p)
  t_spearman_est_vec <- c(t_spearman_est_vec,t_spearman_est)
  t_spearman_p_vec <- c(t_spearman_p_vec,t_spearman_p)
  t_cor_est_vec <- c(t_cor_est_vec,t_est)
  t_cor_p_vec <- c(t_cor_p_vec,t_p)


  PIC_spearman_est_vec <- c(PIC_spearman_est_vec,PIC_spearman_est)
  PIC_spearman_p_vec <- c(PIC_spearman_p_vec,PIC_spearman_p)
  PIC_pearson_est_vec <- c(PIC_pearson_est_vec,PIC_pearson_est)
  PIC_pearson_p_vec <- c(PIC_pearson_p_vec,PIC_pearson_p)
  PIC_est_vec <- c(PIC_est_vec,PIC_est)
  PIC_p_vec <- c(PIC_p_vec,PIC_p)
  
}

my_results <- data.frame(
  t_pearson_est_vec,
  t_pearson_p_vec,
  t_spearman_est_vec,
  t_spearman_p_vec,
  t_cor_est_vec,
  t_cor_p_vec,
  
  PIC_spearman_est_vec,
  PIC_spearman_p_vec,
  PIC_pearson_est_vec,
  PIC_pearson_p_vec,
  PIC_est_vec,
  PIC_p_vec,
  
  final_est_vec,
  final_p_vec
)

colnames(my_results)<- c("t_spearman_est",
                         "t_spearman_p",
                         "t_pearson_est",
                         "t_pearson_p",
                         "t_est",
                         "t_p",
                         
                         "PIC_spearman_est",
                         "PIC_spearman_p",
                         "PIC_pearson_est",
                         "PIC_pearson_p",
                         "PIC_est",
                         "PIC_p",
                         
                         "final_est",
                         "final_p")

out_file <- paste0("results_1/",args,"_results.csv")
write.csv(my_results,out_file)
```
