## Supplementary material for "Improving the Robustness of Phylogenetic Independent Contrasts: Addressing Abrupt Evolutionary Shifts with Outlier-and Distribution-Guided Correlation": Codes used in this study.: 03Analysis_02_PGLS_RPR.R

```
command <- paste0("ls data | wc -l")
system(command)
args <- commandArgs(trailingOnly = TRUE)
library(ape)
library(phylolm)
library(geiger)
library(forecast)
library(ROBRT)

get_history_change <- function(all_df,relation_df,X){
  L <- length(rownames(relation_df))
  res_vec <- NULL
  for(i in c(1:L)){
    parent <- relation_df$Parent[i]
    child <- relation_df$Child[i]
    parent_trait_X <- all_df[all_df$ID==parent,X]
    child_trait_X <- all_df[all_df$ID==child,X]
    change_X <- (child_trait_X - parent_trait_X)/relation_df$distance[i]
    res_vec <- c(res_vec,change_X)
  }  
  return(res_vec)
}

calculate_M_estimator_p <- function(t_value,tips_num){
  p_value <- 2 * (1 - pt(abs(t_value), tips_num-1))
  return(p_value)
}

index_num <- 13


#### 1. define NULL vector #####
xy_est_vec <- NULL
xy_p_vec <- NULL
yx_est_vec <- NULL
yx_p_vec <- NULL
phylo_lm_est_vec <- NULL
phylo_lm_p_vec <- NULL
pc_vec <- NULL
pc_p_vec <- NULL
change_est_vec <- NULL
change_p_vec <- NULL
spearman_est_vec <- NULL
spearman_p_vec <- NULL
final_est_vec <- NULL
final_p_vec <- NULL
X1_norm_p_vec <- NULL
X2_norm_p_vec <- NULL

X1_lambda_vec <- NULL
X1_lambda_p_vec <- NULL
X2_lambda_vec <- NULL
X2_lambda_p_vec <- NULL

X1_K_vec <- NULL
X1_K_p_vec <- NULL
X2_K_vec <- NULL
X2_K_p_vec <- NULL

pvr_signal_vec <- NULL
pgls_signal_vec <- NULL
degree_vec <- NULL
OUfixed_xy_est_vec <- NULL
OUfixed_xy_p_vec <- NULL
OUfixed_xy_opt_vec <- NULL
OUfixed_xy_aic_vec <- NULL
OUfixed_xy_loglik_vec <- NULL
OUfixed_xy_rsq_vec <- NULL

OUfixed_yx_est_vec <- NULL
OUfixed_yx_p_vec <- NULL
OUfixed_yx_opt_vec <- NULL
OUfixed_yx_aic_vec <- NULL
OUfixed_yx_loglik_vec <- NULL
OUfixed_yx_rsq_vec <- NULL

lambda_xy_est_vec <- NULL
lambda_xy_p_vec <- NULL
lambda_xy_opt_vec <- NULL
lambda_xy_aic_vec <- NULL
lambda_xy_loglik_vec <- NULL
lambda_xy_rsq_vec <- NULL

lambda_yx_est_vec <- NULL
lambda_yx_p_vec <- NULL
lambda_yx_opt_vec <- NULL
lambda_yx_aic_vec <- NULL
lambda_yx_loglik_vec <- NULL
lambda_yx_rsq_vec <- NULL

EB_xy_est_vec <- NULL
EB_xy_p_vec <- NULL
EB_xy_opt_vec <- NULL
EB_xy_aic_vec <- NULL
EB_xy_loglik_vec <- NULL
EB_xy_rsq_vec <- NULL

EB_yx_est_vec <- NULL
EB_yx_p_vec <- NULL
EB_yx_opt_vec <- NULL
EB_yx_aic_vec <- NULL
EB_yx_loglik_vec <- NULL
EB_yx_rsq_vec <- NULL

OUrandom_xy_est_vec <- NULL
OUrandom_xy_p_vec <- NULL
OUrandom_xy_opt_vec <- NULL
OUrandom_xy_aic_vec <- NULL
OUrandom_xy_loglik_vec <- NULL
OUrandom_xy_rsq_vec <- NULL

OUrandom_yx_est_vec <- NULL
OUrandom_yx_p_vec <- NULL
OUrandom_yx_opt_vec <- NULL
OUrandom_yx_aic_vec <- NULL
OUrandom_yx_loglik_vec <- NULL
OUrandom_yx_rsq_vec <- NULL


BM_xy_est_vec <- NULL
BM_xy_p_vec <- NULL
BM_xy_aic_vec <- NULL
BM_xy_loglik_vec <- NULL
BM_xy_rsq_vec <- NULL

BM_yx_est_vec <- NULL
BM_yx_p_vec <- NULL
BM_yx_aic_vec <- NULL
BM_yx_loglik_vec <- NULL 
BM_yx_rsq_vec <- NULL

OLS_xy_est_vec <- NULL
OLS_xy_p_vec <- NULL
OLS_xy_aic_vec <- NULL
OLS_xy_loglik_vec <- NULL
OLS_xy_rsq_vec <- NULL

OLS_yx_est_vec <- NULL
OLS_yx_p_vec <- NULL
OLS_yx_aic_vec <- NULL
OLS_yx_loglik_vec <- NULL
OLS_yx_rsq_vec <- NULL


sig_label_vec <- NULL

PIC_MM_xy_est_vec <- NULL 
PIC_MM_xy_p_vec   <- NULL
PIC_S_xy_est_vec <- NULL
PIC_S_xy_p_vec   <- NULL
PIC_L1_xy_est_vec <- NULL
PIC_L1_xy_p_vec   <- NULL
PIC_L2_xy_est_vec <- NULL
PIC_L2_xy_p_vec   <- NULL
PIC_M_xy_est_vec <- NULL
PIC_M_xy_p_vec <- NULL


PIC_MM_yx_est_vec <- NULL 
PIC_MM_yx_p_vec   <- NULL
PIC_S_yx_est_vec <- NULL
PIC_S_yx_p_vec   <- NULL
PIC_L1_yx_est_vec <- NULL
PIC_L1_yx_p_vec   <- NULL
PIC_L2_yx_est_vec <- NULL
PIC_L2_yx_p_vec   <- NULL
PIC_M_yx_est_vec <- NULL
PIC_M_yx_p_vec <- NULL


PGLS_MM_xy_est_vec <- NULL 
PGLS_MM_xy_p_vec   <- NULL
PGLS_S_xy_est_vec <- NULL
PGLS_S_xy_p_vec   <- NULL
PGLS_L1_xy_est_vec <- NULL
PGLS_L1_xy_p_vec   <- NULL
PGLS_L2_xy_est_vec <- NULL
PGLS_L2_xy_p_vec   <- NULL
PGLS_M_xy_est_vec <- NULL 
PGLS_M_xy_p_vec   <- NULL


PGLS_MM_yx_est_vec <- NULL 
PGLS_MM_yx_p_vec   <- NULL
PGLS_S_yx_est_vec <- NULL
PGLS_S_yx_p_vec   <- NULL
PGLS_L1_yx_est_vec <- NULL
PGLS_L1_yx_p_vec   <- NULL
PGLS_L2_yx_est_vec <- NULL
PGLS_L2_yx_p_vec   <- NULL
PGLS_M_yx_est_vec <- NULL
PGLS_M_yx_p_vec <- NULL


#index<-1
#### 2. batch process ####
for (index in c(1:index_num)){
  sig_label <- "O"
  tips_df_file <- paste0("data/",index,"/tips_states.csv")
  all_df_file <- paste0("data/",index,"/all_states.csv")
  relation_file <- paste0("data/",index,"/Parent2Child.txt")
  my_tree_file <- paste0("data/",index,"/mytree.nwk")
  relation_df <- read.table(relation_file,sep = " ",header = T)
  all_df <- read.csv(all_df_file)
  tips_df <- read.csv(tips_df_file)
  my_tree <- read.tree(my_tree_file)
  
  tips_num <- length(tips_df$ID)
  relation_df$distance <- my_tree$edge.length
  #### 2.1 PGLS ####
  ##### 2.1.1 OUfixedRoot part  X1~X2####
  OUfixed_xy_fit <- phylolm(X1~X2,data=tips_df,phy=my_tree,model = "OUfixedRoot")
  OUfixed_xy_est <- summary(OUfixed_xy_fit)$coefficients[2,1]
  OUfixed_xy_p <- summary(OUfixed_xy_fit)$coefficients[2,4]
  OUfixed_xy_opt <- OUfixed_xy_fit$optpar
  OUfixed_xy_aic <- OUfixed_xy_fit$aic
  OUfixed_xy_loglik <- OUfixed_xy_fit$logLik
  OUfixed_xy_r_square <- OUfixed_xy_fit$r.squared
  
  ##### 2.1.2 OUfixedRoot part  X2~X1####
  OUfixed_yx_fit <- phylolm(X2~X1,data=tips_df,phy=my_tree,model = "OUfixedRoot")
  OUfixed_yx_est <- summary(OUfixed_yx_fit)$coefficients[2,1]
  OUfixed_yx_p <- summary(OUfixed_yx_fit)$coefficients[2,4]
  OUfixed_yx_opt <- OUfixed_yx_fit$optpar
  OUfixed_yx_aic <- OUfixed_yx_fit$aic
  OUfixed_yx_loglik <- OUfixed_yx_fit$logLik
  OUfixed_yx_r_square <- OUfixed_yx_fit$r.squared
  
  ##### 2.1.3 OUrandom part  X1~X2####
  OUrandom_xy_fit <- phylolm(X1~X2,data=tips_df,phy=my_tree,model = "OUrandomRoot")
  OUrandom_xy_est <- summary(OUrandom_xy_fit)$coefficients[2,1]
  OUrandom_xy_p <- summary(OUrandom_xy_fit)$coefficients[2,4]
  OUrandom_xy_opt <- OUrandom_xy_fit$optpar
  OUrandom_xy_aic <- OUrandom_xy_fit$aic
  OUrandom_xy_loglik <- OUrandom_xy_fit$logLik
  OUrandom_xy_r_square <- OUrandom_xy_fit$r.squared
  
  ##### 2.1.4 OUrandom part  X2~X1####
  OUrandom_yx_fit <- phylolm(X2~X1,data=tips_df,phy=my_tree,model = "OUrandomRoot")
  OUrandom_yx_est <- summary(OUrandom_yx_fit)$coefficients[2,1]
  OUrandom_yx_p <- summary(OUrandom_yx_fit)$coefficients[2,4]
  OUrandom_yx_opt <- OUrandom_yx_fit$optpar
  OUrandom_yx_aic <- OUrandom_yx_fit$aic
  OUrandom_yx_loglik <- OUrandom_yx_fit$logLik
  OUrandom_yx_r_square <- OUrandom_yx_fit$r.squared
  
  
  ##### 2.1.5 lambda part  X1~X2####
  lambda_xy_fit <- phylolm(X1~X2,data=tips_df,phy=my_tree,model = "lambda")
  lambda_xy_est <- summary(lambda_xy_fit)$coefficients[2,1]
  lambda_xy_p <- summary(lambda_xy_fit)$coefficients[2,4]
  lambda_xy_opt <- lambda_xy_fit$optpar
  lambda_xy_aic <- lambda_xy_fit$aic
  lambda_xy_loglik <- lambda_xy_fit$logLik
  lambda_xy_r_square <- lambda_xy_fit$r.squared
  
  ##### 2.1.6 lambda part  X2~X1####
  lambda_yx_fit <- phylolm(X2~X1,data=tips_df,phy=my_tree,model = "lambda")
  lambda_yx_est <- summary(lambda_yx_fit)$coefficients[2,1]
  lambda_yx_p <- summary(lambda_yx_fit)$coefficients[2,4]
  lambda_yx_opt <- lambda_yx_fit$optpar
  lambda_yx_aic <- lambda_yx_fit$aic
  lambda_yx_loglik <- lambda_yx_fit$logLik
  lambda_yx_r_square <- lambda_yx_fit$r.squared
  
  ##### 2.1.7 EB part  X1~X2####
  EB_xy_fit <- phylolm(X1~X2,data=tips_df,phy=my_tree,model = "EB")
  EB_xy_est <- summary(EB_xy_fit)$coefficients[2,1]
  EB_xy_p <- summary(EB_xy_fit)$coefficients[2,4]
  EB_xy_opt <- EB_xy_fit$optpar
  EB_xy_aic <- EB_xy_fit$aic
  EB_xy_loglik <- EB_xy_fit$logLik
  EB_xy_r_square <- EB_xy_fit$r.squared
  
  ##### 2.1.8 EB part  X2~X1####
  EB_yx_fit <- phylolm(X2~X1,data=tips_df,phy=my_tree,model = "EB")
  EB_yx_est <- summary(EB_yx_fit)$coefficients[2,1]
  EB_yx_p <- summary(EB_yx_fit)$coefficients[2,4]
  EB_yx_opt <- EB_yx_fit$optpar
  EB_yx_aic <- EB_yx_fit$aic
  EB_yx_loglik <- EB_yx_fit$logLik
  EB_yx_r_square <- EB_yx_fit$r.squared
  
  ##### 2.1.9 BM part X1~X2####
  BM_xy_fit <- phylolm(X1~X2,data=tips_df,phy=my_tree,model = "BM")
  BM_xy_est <- summary(BM_xy_fit)$coefficients[2,1]
  BM_xy_p <- summary(BM_xy_fit)$coefficients[2,4]
  BM_xy_aic <- BM_xy_fit$aic
  BM_xy_loglik <- BM_xy_fit$logLik
  BM_xy_r_square <- BM_xy_fit$r.squared
  
  ##### 2.1.10 BM part  X2~X1####
  BM_yx_fit <- phylolm(X2~X1,data=tips_df,phy=my_tree,model = "BM")
  BM_yx_est <- summary(BM_yx_fit)$coefficients[2,1]
  BM_yx_p <- summary(BM_yx_fit)$coefficients[2,4]
  BM_yx_aic <- BM_yx_fit$aic
  BM_yx_loglik <- BM_yx_fit$logLik
  BM_yx_r_square <- BM_yx_fit$r.squared
  
  #### 2.2 OLS ####
  ##### 2.2.1 OLS part  X1~X2 ####
  OLS_xy_fit <- lm(X1~X2,data=tips_df)
  OLS_xy_est <- summary(OLS_xy_fit)$coefficients[2,1]
  OLS_xy_p <- summary(OLS_xy_fit)$coefficients[2,4]
  
  ##### 2.2.2 OLS part  X2~X1####
  OLS_yx_fit <- lm(X2~X1,data=tips_df)
  OLS_yx_est <- summary(OLS_yx_fit)$coefficients[2,1]
  OLS_yx_p <- summary(OLS_yx_fit)$coefficients[2,4]
  
  
  #### 2.3 Robust Phylogenetic regression ####
  ##### 2.3.1 PIC -M,MM,S,L1,L2 ####
  ###### 2.3.1.1 PIC X1~X2 ####
  PIC_xy_model <- Conduct.Robust_PhylogeneticRegression(handle.Phylogeny = my_tree,
                                                        Y.trait = tips_df[,"X1"],
                                                        X.traits = cbind(tips_df[,"X2"]),
                                                        vector.Estimators = c("L2","L1","S","M","MM"),
                                                        string.Method = "PIC")
  
  PIC_MM_xy_est <- summary(PIC_xy_model$MM)$coefficients[1,1]
  PIC_MM_xy_p   <- summary(PIC_xy_model$MM)$coefficients[1,4]
  PIC_S_xy_est <- summary(PIC_xy_model$S)$coefficients[1,1]
  PIC_S_xy_p   <- summary(PIC_xy_model$S)$coefficients[1,4]
  PIC_L1_xy_est <- summary(PIC_xy_model$L1)$coefficients[1,1]
  PIC_L1_xy_p   <- summary(PIC_xy_model$L1)$coefficients[1,4]
  PIC_L2_xy_est <- summary(PIC_xy_model$L2)$coefficients[1,1]
  PIC_L2_xy_p   <- summary(PIC_xy_model$L2)$coefficients[1,4]
  PIC_M_xy_est <- summary(PIC_xy_model$M)$coefficients[1,1]
  PIC_M_xy_t <- summary(PIC_xy_model$M)$coefficients[1,3]
  PIC_M_xy_p <- calculate_M_estimator_p(PIC_M_xy_t,tips_num)
  
  
  ###### 2.3.1.2 PIC X2~X1 ####
  PIC_yx_model <- Conduct.Robust_PhylogeneticRegression(handle.Phylogeny = my_tree,
                                                        Y.trait = tips_df[,"X2"],
                                                        X.traits = cbind(tips_df[,"X1"]),
                                                        vector.Estimators = c("L2","L1","S","M","MM"),
                                                        string.Method = "PIC")
  
  PIC_MM_yx_est <- summary(PIC_yx_model$MM)$coefficients[1,1]
  PIC_MM_yx_p   <- summary(PIC_yx_model$MM)$coefficients[1,4]
  PIC_S_yx_est <- summary(PIC_yx_model$S)$coefficients[1,1]
  PIC_S_yx_p   <- summary(PIC_yx_model$S)$coefficients[1,4]
  PIC_L1_yx_est <- summary(PIC_yx_model$L1)$coefficients[1,1]
  PIC_L1_yx_p   <- summary(PIC_yx_model$L1)$coefficients[1,4]
  PIC_L2_yx_est <- summary(PIC_yx_model$L2)$coefficients[1,1]
  PIC_L2_yx_p   <- summary(PIC_yx_model$L2)$coefficients[1,4]
  PIC_M_yx_est <- summary(PIC_yx_model$M)$coefficients[1,1]
  PIC_M_yx_t <- summary(PIC_yx_model$M)$coefficients[1,3]
  PIC_M_yx_p <- calculate_M_estimator_p(PIC_M_yx_t,tips_num)
  
  
  ##### 2.3.1 PGLS -M,MM,S,L1,L2 ####
  ###### 2.3.1.1 PGLS X1~X2 ####
  PGLS_xy_model <- Conduct.Robust_PhylogeneticRegression(handle.Phylogeny = my_tree,
                                                         Y.trait = tips_df[,"X1"],
                                                         X.traits = tips_df[,"X2"],
                                                         vector.Estimators = c("L2","L1","S","M","MM"),
                                                         string.Method = "PGLS")
  
  PGLS_MM_xy_est <- summary(PGLS_xy_model$MM)$coefficients[2,1]
  PGLS_MM_xy_p   <- summary(PGLS_xy_model$MM)$coefficients[2,4]
  PGLS_S_xy_est <- summary(PGLS_xy_model$S)$coefficients[2,1]
  PGLS_S_xy_p   <- summary(PGLS_xy_model$S)$coefficients[2,4]
  PGLS_L1_xy_est <- summary(PGLS_xy_model$L1)$coefficients[2,1]
  PGLS_L1_xy_p   <- summary(PGLS_xy_model$L1)$coefficients[2,4]
  PGLS_L2_xy_est <- summary(PGLS_xy_model$L2)$coefficients[2,1]
  PGLS_L2_xy_p   <- summary(PGLS_xy_model$L2)$coefficients[2,4]
  PGLS_M_xy_est <- summary(PGLS_xy_model$M)$coefficients[2,1]
  PGLS_M_xy_t <- summary(PGLS_xy_model$M)$coefficients[2,3]
  PGLS_M_xy_p <- calculate_M_estimator_p(PGLS_M_xy_t,tips_num)
  
  
  
  
  ###### 2.3.1.2 PGLS X2~X1 ####
  PGLS_yx_model <- Conduct.Robust_PhylogeneticRegression(handle.Phylogeny = my_tree,
                                                         Y.trait = tips_df[,"X2"],
                                                         X.traits = tips_df[,"X1"],
                                                         vector.Estimators = c("L2","L1","S","M","MM"),
                                                         string.Method = "PGLS")
  
  PGLS_MM_yx_est <- summary(PGLS_yx_model$MM)$coefficients[2,1]
  PGLS_MM_yx_p   <- summary(PGLS_yx_model$MM)$coefficients[2,4]
  PGLS_S_yx_est <- summary(PGLS_yx_model$S)$coefficients[2,1]
  PGLS_S_yx_p   <- summary(PGLS_yx_model$S)$coefficients[2,4]
  PGLS_L1_yx_est <- summary(PGLS_yx_model$L1)$coefficients[2,1]
  PGLS_L1_yx_p   <- summary(PGLS_yx_model$L1)$coefficients[2,4]
  PGLS_L2_yx_est <- summary(PGLS_yx_model$L2)$coefficients[2,1]
  PGLS_L2_yx_p   <- summary(PGLS_yx_model$L2)$coefficients[2,4]
  PGLS_M_yx_est <- summary(PGLS_yx_model$M)$coefficients[2,1]
  PGLS_M_yx_t <- summary(PGLS_yx_model$M)$coefficients[2,3]
  PGLS_M_yx_p <- calculate_M_estimator_p(PGLS_M_yx_t,tips_num)
  
  
  
  #### 2.5 gold standard ####
  ## history PIC ###
  X1_change <- get_history_change(all_df,relation_df,"X1")
  X2_change <- get_history_change(all_df,relation_df,"X2")
  
  ##### 2.5.1 original_data ####
  change_formula <- as.formula(paste0("X1_change~X2_change"))
  change_est <- summary(lm(change_formula))$coefficients[2,1]
  change_p <- summary(lm(change_formula))$coefficients[2,4]
  spearman_est <- cor.test(X1_change,X2_change,method="spearman")$estimate
  spearman_p <- cor.test(X1_change,X2_change,method="spearman")["p.value"]$p.value
  
  ## normality test ###
  X1_normal_p <- shapiro.test(X1_change)$p.value
  X2_normal_p <- shapiro.test(X2_change)$p.value
  
  if (X1_normal_p<0.05 | X2_normal_p<0.05){
    final_est <- spearman_est
    final_p <- spearman_p
    pre_norm <- "N"
  }else{
    final_est <- change_est
    final_p <- change_p
    pre_norm <- "Y"
  }
  
  ##### 2.5.2 yeo_data ####
  boxcox_X1_lambda <- forecast::BoxCox.lambda(X1_change, lower = 0)
  boxcox_X2_lambda <- forecast::BoxCox.lambda(X2_change, lower = 0)
  global_lambda <- (boxcox_X1_lambda+boxcox_X2_lambda)/2
  
  yeo_X1_change <- forecast::BoxCox(X1_change, global_lambda)
  yeo_X2_change <- forecast::BoxCox(X2_change, global_lambda)
  
  change_formula <- as.formula(paste0("yeo_X1_change~yeo_X2_change"))
  yeo_pearson_est <- cor.test(yeo_X1_change,yeo_X2_change,method="pearson")$estimate
  yeo_pearson_p <- cor.test(yeo_X1_change,yeo_X2_change,method="pearson")["p.value"]$p.value
  yeo_spearman_est <- cor.test(yeo_X1_change,yeo_X2_change,method="spearman")$estimate
  yeo_spearman_p <- cor.test(yeo_X1_change,yeo_X2_change,method="spearman")["p.value"]$p.value
  
  ## normality test ###
  yeo_X1_normal_p <- shapiro.test(yeo_X1_change)$p.value
  yeo_X2_normal_p <- shapiro.test(yeo_X2_change)$p.value
  
  if (yeo_X1_normal_p<0.05 | yeo_X2_normal_p<0.05){
    yeo_final_est <- yeo_spearman_est
    yeo_final_p <- yeo_spearman_p
    yeo_norm <- "N"
  }else{
    yeo_final_est <- yeo_pearson_est
    yeo_final_p <- yeo_pearson_p
    yeo_norm <- "Y"
  }
  
  ##### 2.5.3 ordered quantile(QRQ) normalization ####
  QRQ_X1_change <- orderNorm(X1_change)$x.t
  QRQ_X2_change <- orderNorm(X2_change)$x.t
  
  
  change_formula <- as.formula(paste0("QRQ_X1_change~QRQ_X2_change"))
  QRQ_pearson_est <- cor.test(QRQ_X1_change,QRQ_X2_change,method="pearson")$estimate
  QRQ_pearson_p <- cor.test(QRQ_X1_change,QRQ_X2_change,method="pearson")["p.value"]$p.value
  QRQ_spearman_est <- cor.test(QRQ_X1_change,QRQ_X2_change,method="spearman")$estimate
  QRQ_spearman_p <- cor.test(QRQ_X1_change,QRQ_X2_change,method="spearman")["p.value"]$p.value
  
  ## normality test ###
  QRQ_X1_normal_p <- shapiro.test(QRQ_X1_change)$p.value
  QRQ_X2_normal_p <- shapiro.test(QRQ_X2_change)$p.value
  
  if (QRQ_X1_normal_p<0.05 | QRQ_X2_normal_p<0.05){
    QRQ_final_est <- QRQ_spearman_est
    QRQ_final_p <- QRQ_spearman_p
    QRQ_norm <- "N"
  }else{
    QRQ_final_est <- QRQ_pearson_est
    QRQ_final_p <- QRQ_pearson_p
    QRQ_norm <- "Y"
  }
  
  
  
  #### 2.6 signal part####
  X1_vec <- tips_df$X1
  X2_vec <- tips_df$X2
  names(X1_vec) <- tips_df$ID
  names(X2_vec) <- tips_df$ID
  X1.lambda <- phylosig(my_tree,X1_vec,method = "lambda",test=T)
  X1.K <- phylosig(my_tree,X1_vec,method = "K",test=T)
  X2.lambda <- phylosig(my_tree,X2_vec,method = "lambda",test=T)
  X2.K <- phylosig(my_tree,X2_vec,method = "K",test=T)
  
  X1_lambda <- X1.lambda$lambda
  X1_lambda_p <- X1.lambda$P
  X1_K <- X1.K$K
  X1_K_p <- X1.K$P
  
  X2_lambda <- X2.lambda$lambda
  X2_lambda_p <- X2.lambda$P
  X2_K <- X2.K$K
  X2_K_p <- X2.K$P
  
  
  
  #### 2.7 add results #####
  ##### 2.7.1 PGLS results add ####
  OUfixed_xy_est_vec <- c(OUfixed_xy_est_vec,OUfixed_xy_est)
  OUfixed_xy_p_vec <- c(OUfixed_xy_p_vec,OUfixed_xy_p)
  OUfixed_xy_opt_vec <- c(OUfixed_xy_opt_vec,OUfixed_xy_opt)
  OUfixed_xy_aic_vec <- c(OUfixed_xy_aic_vec,OUfixed_xy_aic)
  OUfixed_xy_loglik_vec <- c(OUfixed_xy_loglik_vec,OUfixed_xy_loglik)
  OUfixed_xy_rsq_vec <- c(OUfixed_xy_rsq_vec,OUfixed_xy_r_square)
  
  OUfixed_yx_est_vec <- c(OUfixed_yx_est_vec,OUfixed_yx_est)
  OUfixed_yx_p_vec <- c(OUfixed_yx_p_vec,OUfixed_yx_p)
  OUfixed_yx_opt_vec <- c(OUfixed_yx_opt_vec,OUfixed_yx_opt)
  OUfixed_yx_aic_vec <- c(OUfixed_yx_aic_vec,OUfixed_yx_aic)
  OUfixed_yx_loglik_vec <- c(OUfixed_yx_loglik_vec,OUfixed_yx_loglik)
  OUfixed_yx_rsq_vec <- c(OUfixed_yx_rsq_vec,OUfixed_yx_r_square)
  
  OUrandom_xy_est_vec <- c(OUrandom_xy_est_vec,OUrandom_xy_est)
  OUrandom_xy_p_vec <- c(OUrandom_xy_p_vec,OUrandom_xy_p)
  OUrandom_xy_opt_vec <- c(OUrandom_xy_opt_vec,OUrandom_xy_opt)
  OUrandom_xy_aic_vec <- c(OUrandom_xy_aic_vec,OUrandom_xy_aic)
  OUrandom_xy_loglik_vec <- c(OUrandom_xy_loglik_vec,OUrandom_xy_loglik)
  OUrandom_xy_rsq_vec <- c(OUrandom_xy_rsq_vec,OUrandom_xy_r_square)
  
  OUrandom_yx_est_vec <- c(OUrandom_yx_est_vec,OUrandom_yx_est)
  OUrandom_yx_p_vec <- c(OUrandom_yx_p_vec,OUrandom_yx_p)
  OUrandom_yx_opt_vec <- c(OUrandom_yx_opt_vec,OUrandom_yx_opt)
  OUrandom_yx_aic_vec <- c(OUrandom_yx_aic_vec,OUrandom_yx_aic)
  OUrandom_yx_loglik_vec <- c(OUrandom_yx_loglik_vec,OUrandom_yx_loglik)
  OUrandom_yx_rsq_vec <- c(OUrandom_yx_rsq_vec,OUrandom_yx_r_square)
  
  lambda_xy_est_vec <- c(lambda_xy_est_vec,lambda_xy_est)
  lambda_xy_p_vec <- c(lambda_xy_p_vec,lambda_xy_p)
  lambda_xy_opt_vec <- c(lambda_xy_opt_vec,lambda_xy_opt)
  lambda_xy_aic_vec <- c(lambda_xy_aic_vec,lambda_xy_aic)
  lambda_xy_loglik_vec <- c(lambda_xy_loglik_vec,lambda_xy_loglik)
  lambda_xy_rsq_vec <- c(lambda_xy_rsq_vec,lambda_xy_r_square)
  
  lambda_yx_est_vec <- c(lambda_yx_est_vec,lambda_yx_est)
  lambda_yx_p_vec <- c(lambda_yx_p_vec,lambda_yx_p)
  lambda_yx_opt_vec <- c(lambda_yx_opt_vec,lambda_yx_opt)
  lambda_yx_aic_vec <- c(lambda_yx_aic_vec,lambda_yx_aic)
  lambda_yx_loglik_vec <- c(lambda_yx_loglik_vec,lambda_yx_loglik)
  lambda_yx_rsq_vec <- c(lambda_yx_rsq_vec,lambda_yx_r_square)
  
  EB_xy_est_vec <- c(EB_xy_est_vec,EB_xy_est)
  EB_xy_p_vec <- c(EB_xy_p_vec,EB_xy_p)
  EB_xy_opt_vec <- c(EB_xy_opt_vec,EB_xy_opt)
  EB_xy_aic_vec <- c(EB_xy_aic_vec,EB_xy_aic)
  EB_xy_loglik_vec <- c(EB_xy_loglik_vec,EB_xy_loglik)
  EB_xy_rsq_vec <- c(EB_xy_rsq_vec,EB_xy_r_square)
  
  EB_yx_est_vec <- c(EB_yx_est_vec,EB_yx_est)
  EB_yx_p_vec <- c(EB_yx_p_vec,EB_yx_p)
  EB_yx_opt_vec <- c(EB_yx_opt_vec,EB_yx_opt)
  EB_yx_aic_vec <- c(EB_yx_aic_vec,EB_yx_aic)
  EB_yx_loglik_vec <- c(EB_yx_loglik_vec,EB_yx_loglik)
  EB_yx_rsq_vec <- c(EB_yx_rsq_vec,EB_yx_r_square)
  
  
  OLS_xy_est_vec <- c(OLS_xy_est_vec,OLS_xy_est)
  OLS_xy_p_vec <- c(OLS_xy_p_vec,OLS_xy_p)
  OLS_yx_est_vec <- c(OLS_yx_est_vec,OLS_yx_est)
  OLS_yx_p_vec <- c(OLS_yx_p_vec,OLS_yx_p)
  
  
  BM_xy_est_vec <- c(BM_xy_est_vec,BM_xy_est)
  BM_xy_p_vec <- c(BM_xy_p_vec,BM_xy_p)
  BM_xy_aic_vec <- c(BM_xy_aic_vec,BM_xy_aic)
  BM_xy_loglik_vec <- c(BM_xy_loglik_vec,BM_xy_loglik)
  BM_xy_rsq_vec <- c(BM_xy_rsq_vec,BM_xy_r_square)
  
  BM_yx_est_vec <- c(BM_yx_est_vec,BM_yx_est)
  BM_yx_p_vec <- c(BM_yx_p_vec,BM_yx_p)
  BM_yx_aic_vec <- c(BM_yx_aic_vec,BM_yx_aic)
  BM_yx_loglik_vec <- c(BM_yx_loglik_vec,BM_yx_loglik)
  BM_yx_rsq_vec <- c(BM_yx_rsq_vec,BM_yx_r_square)
  
  
  ##### 2.7.2 PVR results add. #####
  
  ##### 2.7.3 Gold standard add #### 
  change_est_vec <- c(change_est_vec,change_est)
  change_p_vec <- c(change_p_vec,change_p)
  spearman_est_vec <- c(spearman_est_vec,spearman_est)
  spearman_p_vec <- c(spearman_p_vec,spearman_p)
  final_est_vec <- c(final_est_vec,final_est)
  final_p_vec <- c(final_p_vec,final_p)
  X1_norm_p_vec <- c(X1_norm_p_vec,X1_normal_p)
  X2_norm_p_vec <- c(X2_norm_p_vec,X2_normal_p)
  pre_norm_vec <- c(pre_norm_vec,pre_norm)
  yeo_norm_vec <- c(yeo_norm_vec,yeo_norm)
  QRQ_norm_vec <- c(QRQ_norm_vec,QRQ_norm)
  
  yeo_pearson_est_vec <- c(yeo_pearson_est_vec,yeo_pearson_est)
  yeo_pearson_p_vec <- c(yeo_pearson_p_vec,yeo_pearson_p)
  yeo_spearman_est_vec <- c(yeo_spearman_est_vec,yeo_spearman_est)
  yeo_spearman_p_vec <- c(yeo_spearman_p_vec,yeo_spearman_p)
  
  QRQ_pearson_est_vec <- c(QRQ_pearson_est_vec,QRQ_pearson_est)
  QRQ_pearson_p_vec <- c(QRQ_pearson_p_vec,QRQ_pearson_p)
  QRQ_spearman_est_vec <- c(QRQ_spearman_est_vec,QRQ_spearman_est)
  QRQ_spearman_p_vec <- c(QRQ_spearman_p_vec,QRQ_spearman_p)
  
  ##### 2.7.4 PROBR add ####
  PIC_MM_xy_est_vec <- c(PIC_MM_xy_est_vec,PIC_MM_xy_est)
  PIC_MM_xy_p_vec   <- c(PIC_MM_xy_p_vec,PIC_MM_xy_p)
  PIC_S_xy_est_vec <- c(PIC_S_xy_est_vec,PIC_S_xy_est)
  PIC_S_xy_p_vec   <- c(PIC_S_xy_p_vec,PIC_S_xy_p)
  PIC_L1_xy_est_vec <- c(PIC_L1_xy_est_vec,PIC_L1_xy_est)
  PIC_L1_xy_p_vec   <- c(PIC_L1_xy_p_vec,PIC_L1_xy_p)
  PIC_L2_xy_est_vec <-c(PIC_L2_xy_est_vec,PIC_L2_xy_est)
  PIC_L2_xy_p_vec   <- c(PIC_L2_xy_p_vec,PIC_L2_xy_p)
  PIC_M_xy_est_vec <- c(PIC_M_xy_est_vec,PIC_M_xy_est)
  PIC_M_xy_p_vec <- c(PIC_M_xy_p_vec,PIC_M_xy_p)   
  
  PIC_MM_yx_est_vec <-  c(PIC_MM_yx_est_vec,PIC_MM_yx_est)
  PIC_MM_yx_p_vec   <- c(PIC_MM_yx_p_vec,PIC_MM_yx_p)
  PIC_S_yx_est_vec <- c(PIC_S_yx_est_vec,PIC_S_yx_est)
  PIC_S_yx_p_vec   <- c(PIC_S_yx_p_vec,PIC_S_yx_p)
  PIC_L1_yx_est_vec <- c(PIC_L1_yx_est_vec,PIC_L1_yx_est)
  PIC_L1_yx_p_vec   <- c(PIC_L1_yx_p_vec,PIC_L1_yx_p)
  PIC_L2_yx_est_vec <- c(PIC_L2_yx_est_vec,PIC_L2_yx_est)
  PIC_L2_yx_p_vec   <- c(PIC_L2_yx_p_vec,PIC_L2_yx_p)
  PIC_M_yx_est_vec <- c(PIC_M_yx_est_vec,PIC_M_yx_est)
  PIC_M_yx_p_vec <- c(PIC_M_yx_p_vec,PIC_M_yx_p)  
  
  
  
  PGLS_MM_xy_est_vec <- c(PGLS_MM_xy_est_vec,PGLS_MM_xy_est)
  PGLS_MM_xy_p_vec   <- c(PGLS_MM_xy_p_vec,PGLS_MM_xy_p)
  PGLS_S_xy_est_vec <- c(PGLS_S_xy_est_vec,PGLS_S_xy_est)
  PGLS_S_xy_p_vec   <- c(PGLS_S_xy_p_vec,PGLS_S_xy_p)
  PGLS_L1_xy_est_vec <- c(PGLS_L1_xy_est_vec,PGLS_L1_xy_est)
  PGLS_L1_xy_p_vec   <- c(PGLS_L1_xy_p_vec,PGLS_L1_xy_p)
  PGLS_L2_xy_est_vec <-c(PGLS_L2_xy_est_vec,PGLS_L2_xy_est)
  PGLS_L2_xy_p_vec   <- c(PGLS_L2_xy_p_vec,PGLS_L2_xy_p)
  PGLS_M_xy_est_vec <- c(PGLS_M_xy_est_vec,PGLS_M_xy_est)
  PGLS_M_xy_p_vec <- c(PGLS_M_xy_p_vec,PGLS_M_xy_p)   
  
  
  
  PGLS_MM_yx_est_vec <-  c(PGLS_MM_yx_est_vec,PGLS_MM_yx_est)
  PGLS_MM_yx_p_vec   <- c(PGLS_MM_yx_p_vec,PGLS_MM_yx_p)
  PGLS_S_yx_est_vec <- c(PGLS_S_yx_est_vec,PGLS_S_yx_est)
  PGLS_S_yx_p_vec   <- c(PGLS_S_yx_p_vec,PGLS_S_yx_p)
  PGLS_L1_yx_est_vec <- c(PGLS_L1_yx_est_vec,PGLS_L1_yx_est)
  PGLS_L1_yx_p_vec   <- c(PGLS_L1_yx_p_vec,PGLS_L1_yx_p)
  PGLS_L2_yx_est_vec <- c(PGLS_L2_yx_est_vec,PGLS_L2_yx_est)
  PGLS_L2_yx_p_vec   <- c(PGLS_L2_yx_p_vec,PGLS_L2_yx_p)
  PGLS_M_yx_est_vec <- c(PGLS_M_yx_est_vec,PGLS_M_yx_est)
  PGLS_M_yx_p_vec <- c(PGLS_M_yx_p_vec,PGLS_M_yx_p)  
  
  
  
  ##### 2.7.5 Signal add ####
  X1_lambda_vec <- c(X1_lambda_vec,X1_lambda)
  X1_lambda_p_vec <- c(X1_lambda_p_vec,X1_lambda_p)
  X2_lambda_vec <- c(X2_lambda_vec,X2_lambda)
  X2_lambda_p_vec <- c(X2_lambda_p_vec,X2_lambda_p)
  X1_K_vec <- c(X1_K_vec,X1_K)
  X1_K_p_vec <- c(X1_K_p_vec,X1_K_p)
  X2_K_vec <- c(X2_K_vec,X2_K)
  X2_K_p_vec <- c(X2_K_p_vec,X2_K_p)
}


### output dataframe #####

my_results <- data.frame(
  X1_lambda <- X1_lambda_vec,
  X1_lambda_p <- X1_lambda_p_vec,
  X2_lambda <- X2_lambda_vec,
  X2_lambda_p <- X2_lambda_p_vec,
  X1_K <- X1_K_vec,
  X1_K_p <- X1_K_p_vec,
  X2_K <- X2_K_vec,
  X2_K_p <- X2_K_p_vec,
  OUfixed_xy_est <- OUfixed_xy_est_vec,
  OUfixed_xy_p <- OUfixed_xy_p_vec,
  OUfixed_xy_opt <- OUfixed_xy_opt_vec,
  OUfixed_xy_aic <- OUfixed_xy_aic_vec,
  OUfixed_xy_loglik <- OUfixed_xy_loglik_vec,
  OUfixed_xy_rsq <- OUfixed_xy_rsq_vec,
  OUfixed_yx_est <- OUfixed_yx_est_vec,
  OUfixed_yx_p <- OUfixed_yx_p_vec,
  OUfixed_yx_opt <- OUfixed_yx_opt_vec,
  OUfixed_yx_aic <- OUfixed_yx_aic_vec,
  OUfixed_yx_loglik <- OUfixed_yx_loglik_vec,
  OUfixed_yx_rsq <- OUfixed_yx_rsq_vec,
  
  OUrandom_xy_est <- OUrandom_xy_est_vec,
  OUrandom_xy_p <- OUrandom_xy_p_vec,
  OUrandom_xy_opt <- OUrandom_xy_opt_vec,
  OUrandom_xy_aic <- OUrandom_xy_aic_vec,
  OUrandom_xy_loglik <- OUrandom_xy_loglik_vec,
  OUrandom_xy_rsq <- OUrandom_xy_rsq_vec,
  OUrandom_yx_est <- OUrandom_yx_est_vec,
  OUrandom_yx_p <- OUrandom_yx_p_vec,
  OUrandom_yx_opt <- OUrandom_yx_opt_vec,
  OUrandom_yx_aic <- OUrandom_yx_aic_vec,
  OUrandom_yx_loglik <- OUrandom_yx_loglik_vec,
  OUrandom_yx_rsq <- OUrandom_yx_rsq_vec,
  
  lambda_xy_est <- lambda_xy_est_vec,
  lambda_xy_p <- lambda_xy_p_vec,
  lambda_xy_opt <- lambda_xy_opt_vec,
  lambda_xy_aic <- lambda_xy_aic_vec,
  lambda_xy_loglik <- lambda_xy_loglik_vec,
  lambda_xy_rsq <- lambda_xy_rsq_vec,
  lambda_yx_est <- lambda_yx_est_vec,
  lambda_yx_p <- lambda_yx_p_vec,
  lambda_yx_opt <- lambda_yx_opt_vec,
  lambda_yx_aic <- lambda_yx_aic_vec,
  lambda_yx_loglik <- lambda_yx_loglik_vec,
  lambda_yx_rsq <- lambda_yx_rsq_vec,
  
  EB_xy_est <- EB_xy_est_vec,
  EB_xy_p <- EB_xy_p_vec,
  EB_xy_opt <- EB_xy_opt_vec,
  EB_xy_aic <- EB_xy_aic_vec,
  EB_xy_loglik <- EB_xy_loglik_vec,
  EB_xy_rsq <- EB_xy_rsq_vec,
  EB_yx_est <- EB_yx_est_vec,
  EB_yx_p <- EB_yx_p_vec,
  EB_yx_opt <- EB_yx_opt_vec,
  EB_yx_aic <- EB_yx_aic_vec,
  EB_yx_loglik <- EB_yx_loglik_vec,
  EB_yx_rsq <- EB_yx_rsq_vec,
  
  
  OLS_xy_est <- OLS_xy_est_vec,
  OLS_xy_p <- OLS_xy_p_vec,
  OLS_yx_est <- OLS_yx_est_vec,
  OLS_yx_p <- OLS_yx_p_vec,
  BM_xy_est <- BM_xy_est_vec,
  BM_xy_p <- BM_xy_p_vec,
  BM_xy_aic <- BM_xy_aic_vec,
  BM_xy_loglik <- BM_xy_loglik_vec,
  BM_xy_rsq <- BM_xy_rsq_vec,
  BM_yx_est <- BM_yx_est_vec,
  BM_yx_p <- BM_yx_p_vec,
  BM_yx_aic <- BM_yx_aic_vec,
  BM_yx_loglik <- BM_yx_loglik_vec,
  BM_yx_rsq <- BM_yx_rsq_vec,
  
  change_est <- change_est_vec,
  change_p <- change_p_vec,
  spearman_est <- spearman_est_vec,
  spearman_p <- spearman_p_vec,
  final_est <- final_est_vec,
  final_p <- final_p_vec,
  
  
  X1_norm_p <- X1_norm_p_vec,
  X2_norm_p <- X2_norm_p_vec,
  
  PIC_MM_xy_est <- PIC_MM_xy_est_vec,
  PIC_MM_xy_p   <- PIC_MM_xy_p_vec,
  PIC_S_xy_est <- PIC_S_xy_est_vec,
  PIC_S_xy_p  <- PIC_S_xy_p_vec,
  PIC_L1_xy_est <- PIC_L1_xy_est_vec,
  PIC_L1_xy_p   <- PIC_L1_xy_p_vec,
  PIC_L2_xy_est <- PIC_L2_xy_est_vec,
  PIC_L2_xy_p   <- PIC_L2_xy_p_vec,
  PIC_M_xy_est <- PIC_M_xy_est_vec,
  PIC_M_xy_p <- PIC_M_xy_p_vec,
  
  
  
  
  PIC_MM_yx_est <-  PIC_MM_yx_est_vec,
  PIC_MM_yx_p   <- PIC_MM_yx_p_vec,
  PIC_S_yx_est <- PIC_S_yx_est_vec,
  PIC_S_yx_p   <- PIC_S_yx_p_vec,
  PIC_L1_yx_est <- PIC_L1_yx_est_vec,
  PIC_L1_yx_p   <- PIC_L1_yx_p_vec,
  PIC_L2_yx_est <- PIC_L2_yx_est_vec,
  PIC_L2_yx_p   <- PIC_L2_yx_p_vec,
  PIC_M_yx_est <- PIC_M_yx_est_vec,
  PIC_M_yx_p <- PIC_M_yx_p_vec,
  
  
  
  PGLS_MM_xy_est <- PGLS_MM_xy_est_vec,
  PGLS_MM_xy_p   <- PGLS_MM_xy_p_vec,
  PGLS_S_xy_est <- PGLS_S_xy_est_vec,
  PGLS_S_xy_p  <- PGLS_S_xy_p_vec,
  PGLS_L1_xy_est <- PGLS_L1_xy_est_vec,
  PGLS_L1_xy_p   <- PGLS_L1_xy_p_vec,
  PGLS_L2_xy_est <- PGLS_L2_xy_est_vec,
  PGLS_L2_xy_p   <- PGLS_L2_xy_p_vec,
  PGLS_M_xy_est <- PGLS_M_xy_est_vec,
  PGLS_M_xy_p <- PGLS_M_xy_p_vec,
  
  
  
  PGLS_MM_yx_est <-  PGLS_MM_yx_est_vec,
  PGLS_MM_yx_p   <- PGLS_MM_yx_p_vec,
  PGLS_S_yx_est <- PGLS_S_yx_est_vec,
  PGLS_S_yx_p   <- PGLS_S_yx_p_vec,
  PGLS_L1_yx_est <- PGLS_L1_yx_est_vec,
  PGLS_L1_yx_p   <- PGLS_L1_yx_p_vec,
  PGLS_L2_yx_est <- PGLS_L2_yx_est_vec,
  PGLS_L2_yx_p   <- PGLS_L2_yx_p_vec,
  PGLS_M_yx_est <- PGLS_M_yx_est_vec,
  PGLS_M_yx_p <- PGLS_M_yx_p_vec
  
  
)


#round_results <- round(my_results,3)
colnames(my_results) <- c( "X1_lambda",
                           "X1_lambda_p",
                           "X2_lambda",
                           "X2_lambda_p",
                           "X1_K",
                           "X1_K_p",
                           "X2_K",
                           "X2_K_p",
                           "OUfixed_xy_est",
                           "OUfixed_xy_p",
                           "OUfixed_xy_opt",
                           "OUfixed_xy_aic",
                           "OUfixed_xy_loglik",
                           "OUfixed_xy_rsq",
                           "OUfixed_yx_est",
                           "OUfixed_yx_p",
                           "OUfixed_yx_opt",
                           "OUfixed_yx_aic",
                           "OUfixed_yx_loglik",
                           "OUfixed_yx_rsq",
                           
                           "OUrandom_xy_est",
                           "OUrandom_xy_p",
                           "OUrandom_xy_opt",
                           "OUrandom_xy_aic",
                           "OUrandom_xy_loglik",
                           "OUrandom_xy_rsq",
                           "OUrandom_yx_est",
                           "OUrandom_yx_p",
                           "OUrandom_yx_opt",
                           "OUrandom_yx_aic",
                           "OUrandom_yx_loglik",
                           "OUrandom_yx_rsq",
                           
                           "lambda_xy_est",
                           "lambda_xy_p",
                           "lambda_xy_opt",
                           "lambda_xy_aic",
                           "lambda_xy_loglik",
                           "lambda_xy_rsq",
                           "lambda_yx_est",
                           "lambda_yx_p",
                           "lambda_yx_opt",
                           "lambda_yx_aic",
                           "lambda_yx_loglik",
                           "lambda_yx_rsq",
                           
                           "EB_xy_est",
                           "EB_xy_p",
                           "EB_xy_opt",
                           "EB_xy_aic",
                           "EB_xy_loglik",
                           "EB_xy_rsq",
                           "EB_yx_est",
                           "EB_yx_p",
                           "EB_yx_opt",
                           "EB_yx_aic",
                           "EB_yx_loglik",
                           "EB_yx_rsq",
                           
                           
                           "OLS_xy_est",
                           "OLS_xy_p",
                           "OLS_yx_est",
                           "OLS_yx_p",
                           "BM_xy_est",
                           "BM_xy_p",
                           "BM_xy_aic",
                           "BM_xy_loglik",
                           "BM_xy_rsq",
                           "BM_yx_est",
                           "BM_yx_p",
                           "BM_yx_aic",
                           "BM_yx_loglik",
                           "BM_yx_rsq",
                           
                           
                           "change_est",
                           "change_p",    
                           "spearman_est",
                           "spearman_p",
                           "final_est",
                           "final_p",
    
                           
                           "X1_norm_p",
                           "X2_norm_p",
                           "PIC_MM_xy_est_vec",
                           "PIC_MM_xy_p_vec",
                           "PIC_S_xy_est_vec",
                           "PIC_S_xy_p_vec",
                           "PIC_L1_xy_est_vec",
                           "PIC_L1_xy_p_vec",
                           "PIC_L2_xy_est_vec",
                           "PIC_L2_xy_p_vec",
                           "PIC_M_xy_est_vec",
                           "PIC_M_xy_p_vec",
                           
                           "PIC_MM_yx_est_vec",
                           "PIC_MM_yx_p_vec",
                           "PIC_S_yx_est_vec",
                           "PIC_S_yx_p_vec",
                           "PIC_L1_yx_est_vec",
                           "PIC_L1_yx_p_vec",
                           "PIC_L2_yx_est_vec",
                           "PIC_L2_yx_p_vec",
                           "PIC_M_yx_est_vec",
                           "PIC_M_yx_p_vec",
                           
                           
                           "PGLS_MM_xy_est_vec",
                           "PGLS_MM_xy_p_vec",
                           "PGLS_S_xy_est_vec",
                           "PGLS_S_xy_p_vec",
                           "PGLS_L1_xy_est_vec",
                           "PGLS_L1_xy_p_vec",
                           "PGLS_L2_xy_est_vec",
                           "PGLS_L2_xy_p_vec",
                           "PGLS_M_xy_est_vec",
                           "PGLS_M_xy_p_vec",
                           
                           
                           
                           "PGLS_MM_yx_est_vec",
                           "PGLS_MM_yx_p_vec",
                           "PGLS_S_yx_est_vec",
                           "PGLS_S_yx_p_vec",
                           "PGLS_L1_yx_est_vec",
                           "PGLS_L1_yx_p_vec",
                           "PGLS_L2_yx_est_vec",
                           "PGLS_L2_yx_p_vec",
                           "PGLS_M_yx_est_vec",
                           "PGLS_M_yx_p_vec"
)
#round_results$sig_label <- sig_label_vec

out_file <- paste0("results_2/",args,"_results.csv")
write.csv(my_results,out_file)
```
