## Supplementary material for "Improving the Robustness of Phylogenetic Independent Contrasts: Addressing Abrupt Evolutionary Shifts with Outlier-and Distribution-Guided Correlation": Codes used in this study.: 03Analysis_03_Corphylo.R

```
command <- paste0("ls data | wc -l")
system(command)
args <- commandArgs(trailingOnly = TRUE)
library(ape)


get_history_change <- function(all_df,relation_df,X){
  L <- length(rownames(relation_df))
  res_vec <- NULL
  for(i in c(1:L)){
    parent <- relation_df$Parent[i]
    child <- relation_df$Child[i]
    parent_trait_X <- all_df[all_df$ID==parent,X]
    child_trait_X <- all_df[all_df$ID==child,X]
    change_X <- (child_trait_X - parent_trait_X)/relation_df$distance[i]
    res_vec <- c(res_vec,change_X)
  }  
  return(res_vec)
}

index_num <- 13


final_est_vec <- NULL
final_p_vec <- NULL
corphylo_est_vec <- NULL
corphylo_p_vec <- NULL


for (index in c(1:index_num)){
  sig_label <- "O"
  tips_df_file <- paste0("data/",index,"/tips_states.csv")
  all_df_file <- paste0("data/",index,"/all_states.csv")
  relation_file <- paste0("data/",index,"/Parent2Child.txt")
  my_tree_file <- paste0("data/",index,"/mytree.nwk")
  relation_df <- read.table(relation_file,sep = " ",header = T)
  all_df <- read.csv(all_df_file)
  tips_df <- read.csv(tips_df_file)
  my_tree <- read.tree(my_tree_file)
  
  tips_num <- length(tips_df$ID)
  relation_df$distance <- my_tree$edge.length
  final_est <- NULL
  final_p <- NULL
  
  
  ## golden standard ###
  X1_change <- get_history_change(all_df,relation_df,"X1")
  X2_change <- get_history_change(all_df,relation_df,"X2")
  
  change_formula <- as.formula(paste0("X1_change~X2_change"))
  change_est <- summary(lm(change_formula))$coefficients[2,1]
  change_p <- summary(lm(change_formula))$coefficients[2,4]
  spearman_est <- cor.test(X1_change,X2_change,method="spearman")$estimate
  spearman_p <- cor.test(X1_change,X2_change,method="spearman")["p.value"]$p.value
  
  ## normality test ###
  X1_normal_p <- shapiro.test(X1_change)$p.value
  X2_normal_p <- shapiro.test(X2_change)$p.value
  
  if (X1_normal_p<0.05 | X2_normal_p<0.05){
    final_est <- spearman_est
    final_p <- spearman_p
  }else{
    final_est <- change_est
    final_p <- change_p
  }
  
  final_est_vec <- c(final_est_vec,final_est)
  final_p_vec <- c(final_p_vec,final_p)
  
  X1 <- tips_df[,"X1"]
  X2 <- tips_df[,"X2"]
  
  newX <- data.frame(X1=X1,X2=X2)
  
  rownames(newX) <- tips_df$ID
  
  corphylo_res <- corphylo(X = newX, phy = my_tree)
  
  corphylo_est <-  corphylo_res$cor.matrix[1,2]
  
  # 计算 t 值
  t_value <- corphylo_est * sqrt(tips_num - 2) / sqrt(1 - corphylo_est^2)
  
  # 计算双尾 p 值
  corphylo_p <- 2 * pt(-abs(t_value), df = tips_num - 2)
  

  corphylo_est_vec <- c(corphylo_est_vec,corphylo_est)
  corphylo_p_vec <- c(corphylo_p_vec,corphylo_p)
}

my_results <- data.frame(
  corphylo_est_vec,
  corphylo_p_vec,
  final_est_vec,
  final_p_vec
  
)
  
colnames(my_results)<-c("corphylo_est","corphylo_p",
                        "final_est","final_p")

out_file <- paste0("results_3/",args,"_results.csv")
write.csv(my_results,out_file)
  
```
