## Supplementary material for "Improving the Robustness of Phylogenetic Independent Contrasts: Addressing Abrupt Evolutionary Shifts with Outlier-and Distribution-Guided Correlation": Codes used in this study.: 03Analysis_04_MRPMM.R

```
command <- paste0("ls data | wc -l")
system(command)
args <- commandArgs(trailingOnly = TRUE)
library(ape)
library(phylolm)
library(MCMCglmm)
library(tidyverse)
library(parallel)
library(knitr)

get_history_change <- function(all_df,relation_df,X){
  L <- length(rownames(relation_df))
  res_vec <- NULL
  for(i in c(1:L)){
    parent <- relation_df$Parent[i]
    child <- relation_df$Child[i]
    parent_trait_X <- all_df[all_df$ID==parent,X]
    child_trait_X <- all_df[all_df$ID==child,X]
    change_X <- (child_trait_X - parent_trait_X)/relation_df$distance[i]
    res_vec <- c(res_vec,change_X)
  }  
  return(res_vec)
}

index_num <- 13


calculate_MRPMM_p <- function(r,n){
  t_value <- r * sqrt((n - 2) / (1 - r^2))
  
  df <- n - 2
  
  p_value <- 2 * (1 - pt(abs(t_value), df))
  return(p_value)
}


PGLMM_est_vec <- NULL
PGLMM_p_vec <- NULL
MRPMM_est_vec <- NULL
MRPMM_p_vec <- NULL

final_est_vec <- NULL
final_p_vec <- NULL
corphylo_est_vec <- NULL
corphylo_p_vec <- NULL


for (index in c(1:index_num)){
  sig_label <- "O"
  tips_df_file <- paste0("data/",index,"/tips_states.csv")
  all_df_file <- paste0("data/",index,"/all_states.csv")
  relation_file <- paste0("data/",index,"/Parent2Child.txt")
  my_tree_file <- paste0("data/",index,"/mytree.nwk")
  relation_df <- read.table(relation_file,sep = " ",header = T)
  all_df <- read.csv(all_df_file)
  tips_df <- read.csv(tips_df_file)
  my_tree <- read.tree(my_tree_file)
  
  tips_num <- length(tips_df$ID)
  relation_df$distance <- my_tree$edge.length
  final_est <- NULL
  final_p <- NULL
  
  
  ## golden standard ###
  X1_change <- get_history_change(all_df,relation_df,"X1")
  X2_change <- get_history_change(all_df,relation_df,"X2")
  
  change_formula <- as.formula(paste0("X1_change~X2_change"))
  change_est <- summary(lm(change_formula))$coefficients[2,1]
  change_p <- summary(lm(change_formula))$coefficients[2,4]
  spearman_est <- cor.test(X1_change,X2_change,method="spearman")$estimate
  spearman_p <- cor.test(X1_change,X2_change,method="spearman")["p.value"]$p.value
  
  ## normality test ###
  X1_normal_p <- shapiro.test(X1_change)$p.value
  X2_normal_p <- shapiro.test(X2_change)$p.value
  
  if (X1_normal_p<0.05 | X2_normal_p<0.05){
    final_est <- spearman_est
    final_p <- spearman_p
  }else{
    final_est <- change_est
    final_p <- change_p
  }
  
  final_est_vec <- c(final_est_vec,final_est)
  final_p_vec <- c(final_p_vec,final_p)
  
  colnames(tips_df) <- c("animal","X1","X2")
  
  ### 1.PGLMM ####
  prior <- list(G = list(G1 = list(V = 1, nu = 0.002)), 
                R = list(V = 1, nu = 0.002))
  
  PGLMM_fit <- MCMCglmm(X2 ~ X1,                         # 固定效应：B
                    random = ~animal,           # 随机效应：系统发育关系
                    pedigree = my_tree,         # 系统发育树
                    family = "gaussian",           # 正态分布（根据 A 的分布选择）
                    data = tips_df,             # 包含 A 和 B 的数据框
                    prior = prior,                 # 先验分布
                    nitt = 13000, burnin = 3000, thin = 100)
  
  PGLMM_est <- summary(PGLMM_fit)$solutions[2,1]
  PGLMM_p <- summary(PGLMM_fit)$solutions[2,5]
  
  PGLMM_est_vec <- c(PGLMM_est_vec, PGLMM_est)
  PGLMM_p_vec <- c(PGLMM_p_vec, PGLMM_p)
  
  ### 2.MR_PMM ####
  p.m.2 <- list(G = list(G1 = list(V = diag(2), nu = 1.002)), 
                R = list(R1 = list(V = diag(2), nu = 1.002)))
  
  MRPMM_fit <- MCMCglmm(cbind(X1, X2) ~ trait-1,
                  random   = ~us(trait):animal,
                  rcov     = ~us(trait):units,
                  pedigree = my_tree,
                  family   = c("gaussian","gaussian"), 
                  data     = tips_df, 
                  prior    = p.m.2, 
                  nitt     = 13000, 
                  burnin   = 3000, 
                  thin     = 100,
                  pr = T, verbose = F
  )
  
  residual_correlation <- MRPMM_fit$VCV[,"traitX1:traitX2.units"] /
    sqrt(MRPMM_fit$VCV[,"traitX1:traitX1.units"] * MRPMM_fit$VCV[,"traitX2:traitX2.units"])
  
  MRPMM_est <- quantile(residual_correlation,0.5)
  
  MRPMM_p <- calculate_MRPMM_p(MRPMM_est,tips_num)
  
  MRPMM_est_vec <- c(MRPMM_est_vec, MRPMM_est)
  MRPMM_p_vec <- c(MRPMM_p_vec, MRPMM_p)

}

my_results <- data.frame(
  PGLMM_est_vec,
  PGLMM_p_vec,
  MRPMM_est_vec,
  MRPMM_p_vec,
  final_est_vec,
  final_p_vec
  
)

colnames(my_results)<-c("PGLMM_est","PGLMM_p",
                        "MRPMM_est","MRPMM_p",
                        "final_est","final_p")

out_file <- paste0("results_4/",args,"_results.csv")
write.csv(my_results,out_file)
```
