## Supplementary material for "Improving the Robustness of Phylogenetic Independent Contrasts: Addressing Abrupt Evolutionary Shifts with Outlier-and Distribution-Guided Correlation": Codes used in this study.: 03Analysis_05_MAD.R

```
command <- paste0("ls data | wc -l")
system(command)
args <- commandArgs(trailingOnly = TRUE)
library(ape)
index_num <- 13

change_pearson_est_vec <- NULL
change_pearson_p_vec <- NULL
change_spearman_est_vec <- NULL
change_spearman_p_vec <- NULL
change_X1_norm_p_vec <- NULL
change_X2_norm_p_vec <- NULL
change_X1_mad_max_vec <- NULL
change_X2_mad_max_vec <- NULL

PIC_spearman_est_vec <- NULL
PIC_spearman_p_vec <- NULL
PIC_pearson_est_vec <- NULL
PIC_pearson_p_vec <- NULL

PIC_X1_norm_p_vec <- NULL
PIC_X2_norm_p_vec <- NULL
PIC_X1_mad_max_vec <- NULL
PIC_X2_mad_max_vec <- NULL

count_mad_max <- function(data) {
  
  median_val <- median(data, na.rm = TRUE)
  
  mad_val <- median(abs(data - median_val), na.rm = TRUE)
  
  if (mad_val == 0) {
    mad_val <- 1e-10  
  }
  
  mad_scores <- abs(data - median_val) / mad_val
  
  max_score <- max(mad_scores)
  return(max_score)
}
get_history_change <- function(all_df,relation_df,X){
  L <- length(rownames(relation_df))
  res_vec <- NULL
  for(i in c(1:L)){
    parent <- relation_df$Parent[i]
    child <- relation_df$Child[i]
    parent_trait_X <- all_df[all_df$ID==parent,X]
    child_trait_X <- all_df[all_df$ID==child,X]
    change_X <- (child_trait_X - parent_trait_X)/relation_df$distance[i]
    res_vec <- c(res_vec,change_X)
  }  
  return(res_vec)
}

for (index in c(1:index_num)){
  sig_label <- "O"
  tips_df_file <- paste0("data/",index,"/tips_states.csv")
  all_df_file <- paste0("data/",index,"/all_states.csv")
  relation_file <- paste0("data/",index,"/Parent2Child.txt")
  my_tree_file <- paste0("data/",index,"/mytree.nwk")
  relation_df <- read.table(relation_file,sep = " ",header = T)
  all_df <- read.csv(all_df_file)
  tips_df <- read.csv(tips_df_file)
  my_tree <- read.tree(my_tree_file)
  
  tips_num <- length(tips_df$ID)
  relation_df$distance <- my_tree$edge.length
  change_final_est <- NULL
  change_final_p <- NULL
  
  #### golden standard ####
  X1_change <- get_history_change(all_df,relation_df,"X1")
  X2_change <- get_history_change(all_df,relation_df,"X2")
  
  change_formula <- as.formula(paste0("X1_change~X2_change"))
  change_pearson_est <- summary(lm(change_formula))$coefficients[2,1]
  change_pearson_p <- summary(lm(change_formula))$coefficients[2,4]
  change_spearman_est <- cor.test(X1_change,X2_change,method="spearman")$estimate
  change_spearman_p <- cor.test(X1_change,X2_change,method="spearman")["p.value"]$p.value
  
  change_X1_mad_max <- count_mad_max(X1_change)
  change_X2_mad_max <- count_mad_max(X2_change)
  
  change_X1_normal_p <- shapiro.test(X1_change)$p.value
  change_X2_normal_p <- shapiro.test(X2_change)$p.value
  
  
  #### PIC #### 
  X1 <- tips_df[,"X1"]
  X2 <- tips_df[,"X2"]
  PIC_X1 <- pic(X1,my_tree)
  PIC_X2 <- pic(X2,my_tree)
  
  PIC_formula <- as.formula(paste0("PIC_X1~PIC_X2"))
  PIC_pearson_est <- cor.test(PIC_X1,PIC_X2,method="pearson")$estimate
  PIC_pearson_p <- cor.test(PIC_X1,PIC_X2,method="pearson")["p.value"]$p.value
  PIC_spearman_est <- cor.test(PIC_X1,PIC_X2,method="spearman")$estimate
  PIC_spearman_p <- cor.test(PIC_X1,PIC_X2,method="spearman")["p.value"]$p.value
  
  ## PIC outlier and normality test ###
  PIC_X1_mad_max <- count_mad_max(PIC_X1)
  PIC_X2_mad_max <- count_mad_max(PIC_X2)
  
  PIC_X1_normal_p <- shapiro.test(PIC_X1)$p.value
  PIC_X2_normal_p <- shapiro.test(PIC_X2)$p.value
  
  ### add results ####
  PIC_spearman_est_vec <- c(PIC_spearman_est_vec,PIC_spearman_est)
  PIC_spearman_p_vec <- c(PIC_spearman_p_vec,PIC_spearman_p)
  PIC_pearson_est_vec <- c(PIC_pearson_est_vec,PIC_pearson_est)
  PIC_pearson_p_vec <- c(PIC_pearson_p_vec,PIC_pearson_p)
  
  PIC_X1_norm_p_vec <- c(PIC_X1_norm_p_vec,PIC_X1_normal_p)
  PIC_X2_norm_p_vec <- c(PIC_X2_norm_p_vec,PIC_X2_normal_p)
  PIC_X1_mad_max_vec <- c(PIC_X1_mad_max_vec,PIC_X1_mad_max)
  PIC_X2_mad_max_vec <- c(PIC_X2_mad_max_vec,PIC_X2_mad_max)
  
  change_spearman_est_vec <- c(change_spearman_est_vec,change_spearman_est)
  change_spearman_p_vec <- c(change_spearman_p_vec,change_spearman_p)
  change_pearson_est_vec <- c(change_pearson_est_vec,change_pearson_est)
  change_pearson_p_vec <- c(change_pearson_p_vec,change_pearson_p)
  
  change_X1_norm_p_vec <- c(change_X1_norm_p_vec,change_X1_normal_p)
  change_X2_norm_p_vec <- c(change_X2_norm_p_vec,change_X2_normal_p)
  change_X1_mad_max_vec <- c(change_X1_mad_max_vec,change_X1_mad_max)
  change_X2_mad_max_vec <- c(change_X2_mad_max_vec,change_X2_mad_max)
}

my_results <- data.frame(
  
  PIC_spearman_est_vec,
  PIC_spearman_p_vec,
  PIC_pearson_est_vec,
  PIC_pearson_p_vec,
  PIC_X1_norm_p_vec,
  PIC_X2_norm_p_vec,
  PIC_X1_mad_max_vec,
  PIC_X2_mad_max_vec,
  change_spearman_est_vec,
  change_spearman_p_vec,
  change_pearson_est_vec,
  change_pearson_p_vec,
  change_X1_norm_p_vec,
  change_X2_norm_p_vec,
  change_X1_mad_max_vec,
  change_X2_mad_max_vec
  
)

colnames(my_results)<- c(
  "PIC_spearman_est",
  "PIC_spearman_p",
  "PIC_pearson_est",
  "PIC_pearson_p",
  "PIC_X1_norm_p",
  "PIC_X2_norm_p",
  "PIC_X1_mad_MAX",
  "PIC_X2_mad_MAX",
  "change_spearman_est",
  "change_spearman_p",
  "change_pearson_est",
  "change_pearson_p",
  "change_X1_norm_p",
  "change_X2_norm_p",
  "change_X1_mad_MAX",
  "change_X2_mad_MAX"
)

out_file <- paste0("results_5/",args,"_results.csv")
write.csv(my_results,out_file)
```
