## Supplementary material for "Improving the Robustness of Phylogenetic Independent Contrasts: Addressing Abrupt Evolutionary Shifts with Outlier-and Distribution-Guided Correlation": Codes used in this study.: CalculateIQR.R

```
detect_outliers_binary <- function(data) {

  q1 <- quantile(data, 0.25, na.rm = TRUE)
  q3 <- quantile(data, 0.75, na.rm = TRUE)
  iqr <- q3 - q1

  lower_bound <- q1 - 1.5 * iqr
  upper_bound <- q3 + 1.5 * iqr
  

  binary_outliers <- ifelse(data < lower_bound | data > upper_bound, 1, 0)
  
  return(binary_outliers)
}
```
