## Supplementary results. for "Improving the Robustness of Phylogenetic Independent Contrasts: Addressing Abrupt Evolutionary Shifts with Outlier-and Distribution-Guided Correlation"

### *Supplementary Results S1*

### *Extended Comparisons among Robust Regression Methods*

#### S2.1 Overview

This appendix provides extended results supporting the selection of representative robust regression methods used for comparison with PIC-O(D)GC in the main text. Specifically, it presents detailed performance summaries for a comprehensive set of robust estimators evaluated under both PGLS- and PIC-based regression frameworks, including L1, S, M, and MM estimators, across the full range of simulated evolutionary scenarios.

The analyses reported here were used to identify best-performing robust implementations based on objective, simulation-based performance criteria. While the main text focuses on the final selected methods to streamline presentation, the complete set of results—including additional figures, tables, and descriptive summaries—is provided in this appendix for transparency and reproducibility. Consistent with the recent study (Adams et al. 2024), these extended comparisons confirm that PIC-MM exhibits the most stable and reliable performance among the robust estimators considered.

#### S2.2 Performance on the 128-Species Fixed-Balanced Tree under the β-Based Benchmark

Under evolutionary-shift conditions, PIC-MM consistently ranked among the best-performing methods in terms of error control. In Felsenstein's worst-case scenario with deep-root directional shifts affecting both traits (Fig. AS1a), PIC-MM ranked among the lowest false positive rates across a wide range of shift magnitudes. When only $X_1$ was subject to a deep-root shift (Fig. AS1c), PIC-MM likewise exhibited among the lowest false negative rates, indicating stable performance in detecting true trait associations under severe evolutionary perturbations. PIC-S showed comparable robustness in terms of false negative rates under univariate shifts, but its false positive rates under bivariate worst-case shifts were consistently higher than those of PIC-MM (Fig. AS1a), reflecting weaker control of type I error.

Results from simulations with randomly located trait shifts (Figs. AS1b and AS1d) exhibited qualitatively similar patterns. Under these scenarios, PIC-MM again ranked among the best-performing robust estimators, maintaining relatively low false positive rates for bivariate shifts and low false negative rates for univariate shifts across increasing shift magnitudes. Differences among robust estimators were modest under weak shifts (shift $\leq 4$), but became more pronounced as shift magnitude increased, with several PIC- and PGLS-based methods showing rapidly increasing error rates under stronger shifts (shift $> 4$). These results highlight the general sensitivity of conventional robust regressions to abrupt evolutionary changes, particularly when shifts are large.

In no-shift scenarios characterized by increasing background variance (BM–BM, BM–Norm, and Norm–BM; Figs. AS1e–g), false positive rates were uniformly near zero, and method performance was therefore primarily reflected in false negative rates. Under these conditions,

PIC-MM generally remained among the better-performing methods, although it was not uniformly optimal across all variance regimes. By contrast, PIC-S showed moderately elevated false negative rates in the BM–BM and Norm–BM scenarios, ranking among the poorest-performing methods under these background conditions.

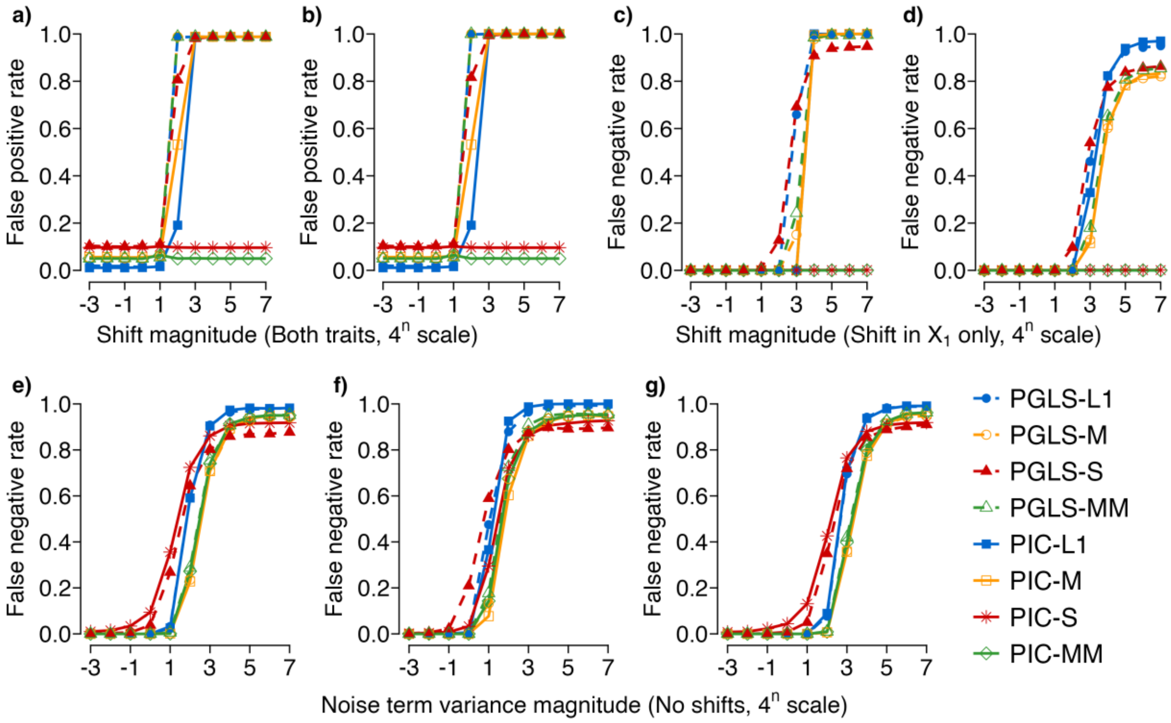


Figure AS1. Evaluation of eight robust phylogenetic regression methods under simulations on a 128-species fixed-balanced tree. Panels a and c correspond to Felsenstein's worst-case scenario, in which strong directional shifts are introduced on deep-root branches. Panels b and d show results from simulations with randomly located trait shifts. Panels a–b report false positive rates under simultaneous shifts in both $X_1$ and $X_2$, whereas panels c–d report false negative rates when only $X_1$ is shifted. Panels e–g show false negative rates under three background scenarios without abrupt shifts: (e) BM–BM, (f) BM–Norm, and (g) Norm–BM. Shift and noise variance magnitudes are expressed on a $4^n$ scale, where n corresponds to the horizontal axis label (e.g., −3 represents $4^{-3}$ and 7 represents $4^7$). False positive and false negative rates are defined relative to the generative parameter $\beta$, using a $\beta$-based benchmark. Additional details on the simulation design, variance gradient configurations, and performance metrics are provided in the Materials and Methods. The data underlying this figure are available in Supplementary Table S21.

#### S2.3 Performance on the 128-Species Fixed-Balanced Tree under the Secondary Benchmark

Under the data-pattern–based benchmark, overall performance patterns were highly consistent with those obtained under the $\beta$-based error-rate benchmark. Across both Felsenstein's worst-case deep-root shift scenarios and simulations with randomly located shifts, relative method rankings remained largely unchanged (Fig. AS2). In particular, PIC-MM again ranked among the best-performing methods across a wide range of shift magnitudes, achieving consistently high accuracy under both bivariate and univariate shift conditions. PIC-S generally followed as a strong alternative, especially under univariate shifts, but exhibited slightly lower accuracy under bivariate shifts across all shift magnitudes.

As in the error-rate analyses, differences among methods were minimal under weak shifts and became increasingly pronounced as shift magnitude increased. Under no-shift background scenarios, most methods achieved high accuracy at low variance levels, whereas performance diverged under higher variance, with qualitative patterns closely mirroring those observed under the β-based benchmark. Together, these results indicate that the main conclusions regarding relative method performance are robust to whether performance is evaluated using the first or the second benchmark.

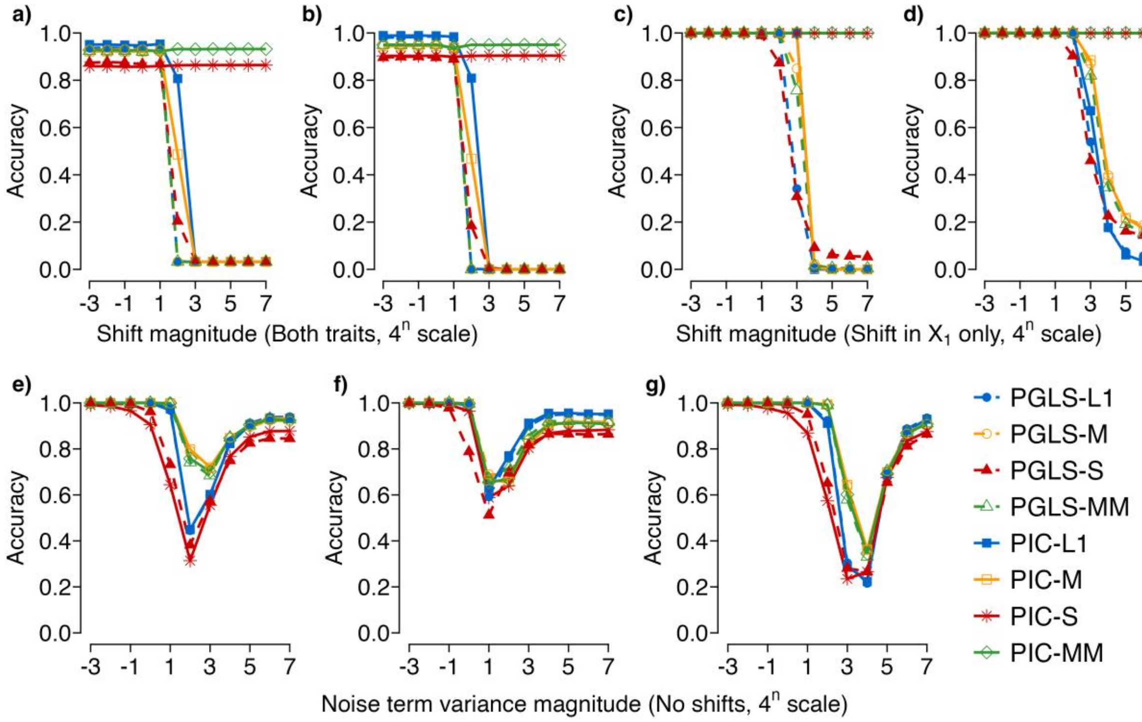


Figure AS2. Evaluation of robust phylogenetic regression methods under the data-pattern–based benchmark on the 128-species fixed-balanced trees. Panel organization and simulation settings are identical to those in Figure AS1, including Felsenstein's worst-case deep-root directional shifts (a, c), randomly located trait shifts (b, d), and no-shift background scenarios, (e) BM–BM, (f) BM–Norm, and (g) Norm–BM. Accuracy is defined as the proportion of simulations in which a method correctly identified both the direction and statistical significance of the trait correlation. The data underlying this figure are available in Supplementary Table S22.

#### S2.4 Extended Comparisons among Robust Regression Methods

To assess the robustness of the conclusions reported above, we extended performance comparisons among robust regression methods across additional phylogenetic configurations, including balanced trees with 16 and 256 species, as well as randomly generated trees with 128 species. Across these alternative settings, qualitative performance patterns closely matched those observed for the 128-species fixed-balanced tree (Fig. AS3-AS8).

Under scenarios involving evolutionary shifts, PIC-MM consistently ranked as the top-performing or among the top-performing robust estimators across all tree sizes and topologies examined (Figs. AS3–AS8). This pattern held across both balanced and randomly generated trees, indicating that its strong performance under shifts is not contingent on a particular phylogenetic configuration. Under gradual-evolution scenarios without abrupt shifts, PIC-MM likewise remained among the stronger-performing methods, although differences among robust estimators were generally smaller and no single method consistently dominated.

Notably, the relative advantages of PIC-MM—and the separation among methods more broadly—were substantially more pronounced in small-sample analyses than in large-sample settings. In particular, for 16-species balanced trees, PIC-MM exhibited clearer separation from other robust alternatives under shift scenarios (Figs. AS3–AS4), whereas performance differences diminished as sample size increased. In larger trees (128 and 256 species), methods tended to converge in performance, with PIC-MM retaining a top-tier position but with reduced contrast relative to other estimators (Figs. AS5–AS8). Together, these results indicate that while PIC-MM performs robustly across sample sizes, its comparative advantages are most evident in small-sample regimes.

Importantly, these qualitative conclusions were invariant to the benchmark used to evaluate robust regression performance. Both the $\beta$-based error-rate benchmark and the accuracy-based benchmark yielded consistent relative rankings among methods, indicating that the observed performance patterns—and the relative advantage of PIC-MM under shift conditions—are not sensitive to the particular evaluation criterion adopted.

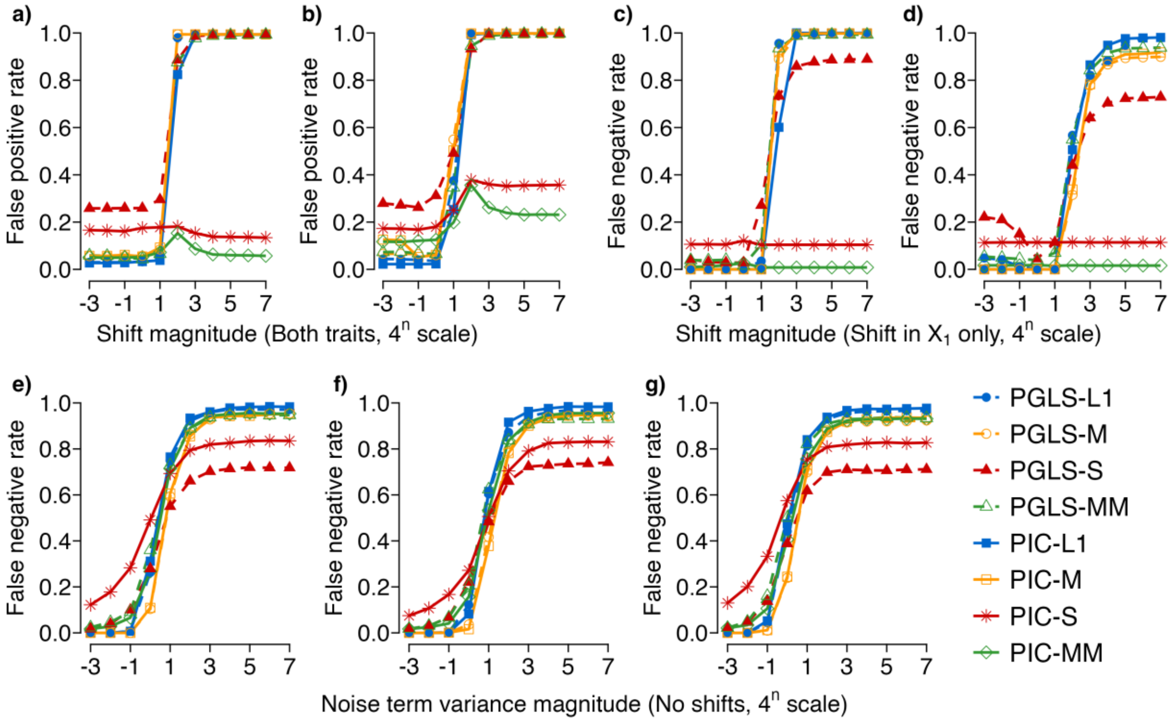


Figure AS3. Evaluation of eight robust phylogenetic regression methods under simulations on a 16-species fixed-balanced tree. Panels a and c correspond to Felsenstein's worst-case scenario, in which strong directional shifts are introduced on deep-root branches. Panels b and d show results from simulations with randomly located trait shifts. Panels a–b report false positive rates under simultaneous shifts in both $X_1$ and $X_2$, whereas panels c–d report false negative rates when only $X_1$ is shifted. Panels e–g show false negative rates under three background scenarios without abrupt shifts: (e) BM–BM, (f) BM–Norm, and (g) Norm–BM. Shift and noise variance magnitudes are expressed on a $4^n$ scale, where n corresponds to the horizontal axis label (e.g., −3 represents $4^{-3}$ and 7 represents $4^7$). False positive and false negative rates are defined relative to the generative parameter $\beta$, using a $\beta$-based benchmark. Additional details on the simulation design, variance gradient configurations, and performance metrics are provided in the Materials and Methods. The data underlying this figure are available in Supplementary Table S23.

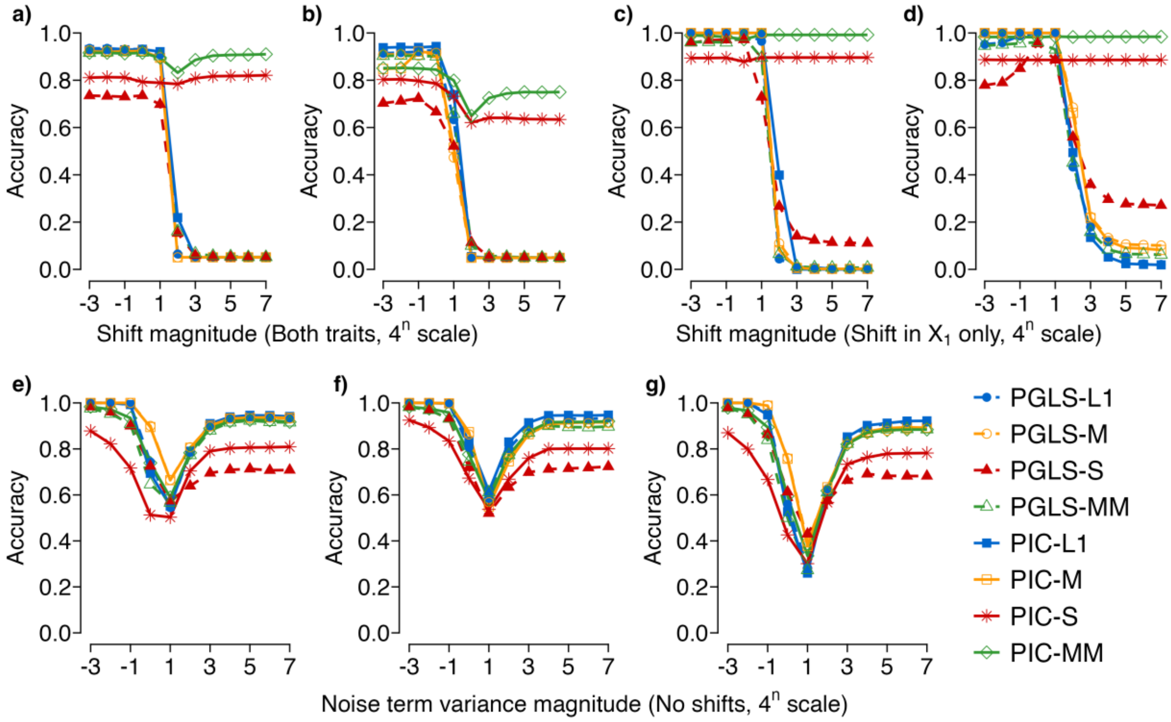


Figure AS4. Evaluation of robust phylogenetic regression methods under the data-pattern–based benchmark on the 16-species fixed-balanced trees. Panel organization and simulation settings are identical to those in Figure AS1, including Felsenstein's worst-case deep-root directional shifts (a, c), randomly located trait shifts (b, d), and no-shift background scenarios, (e) BM–BM, (f) BM–Norm, and (g) Norm–BM. Accuracy is defined as the proportion of simulations in which a method correctly identified both the direction and statistical significance of the trait correlation. The data underlying this figure are available in Supplementary Table S24.

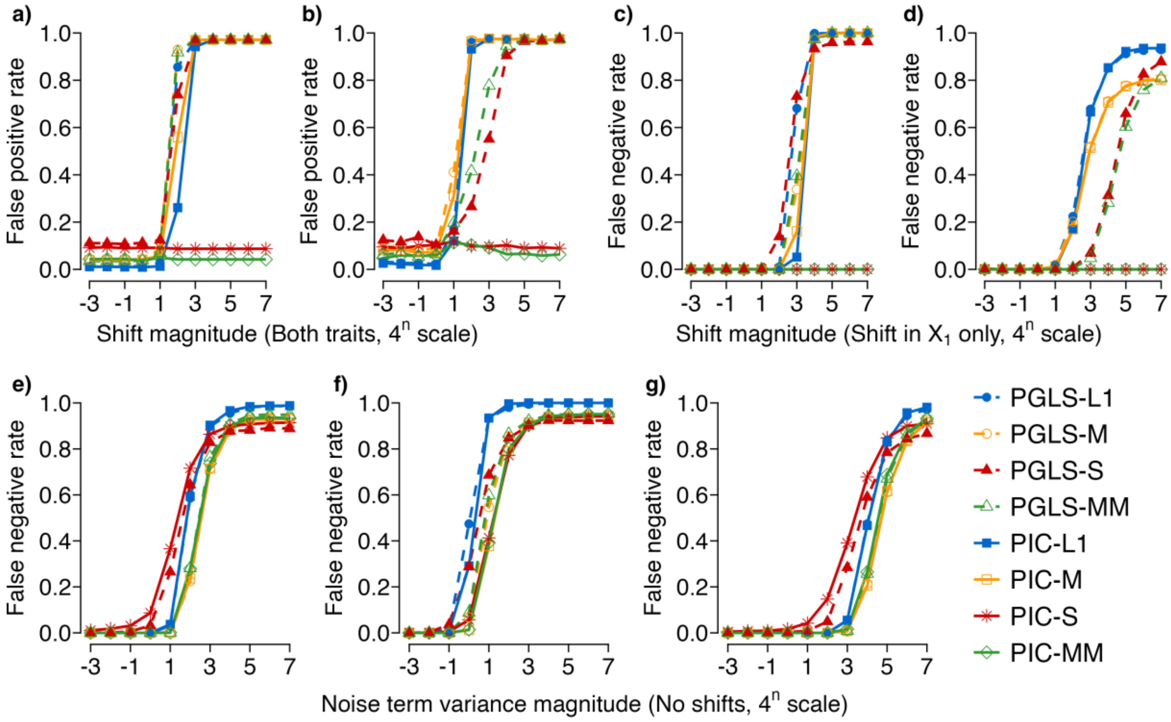


Figure AS5. Evaluation of eight robust phylogenetic regression methods under simulations on the 128-species random trees. Panels a and c correspond to Felsenstein's worst-case scenario, in which strong directional shifts are introduced on deep-root branches. Panels b and d show results from simulations with randomly located trait shifts. Panels a–b report false positive rates under simultaneous shifts in both $X_1$ and $X_2$, whereas panels c–d report false negative rates when only $X_1$ is shifted. Panels e–g show false negative rates under three background scenarios without abrupt shifts: (e) BM–BM, (f) BM–Norm, and (g) Norm–BM. Shift and noise variance magnitudes are expressed on a $4^n$ scale, where n corresponds to the horizontal axis label (e.g., −3 represents $4^{-3}$ and 7 represents $4^7$). False positive and false negative rates are defined relative to the generative parameter $\beta$, using a $\beta$-based benchmark. Additional details on the simulation design, variance gradient configurations, and performance metrics are provided in the Materials and Methods. The data underlying this figure are available in Supplementary Table S25.

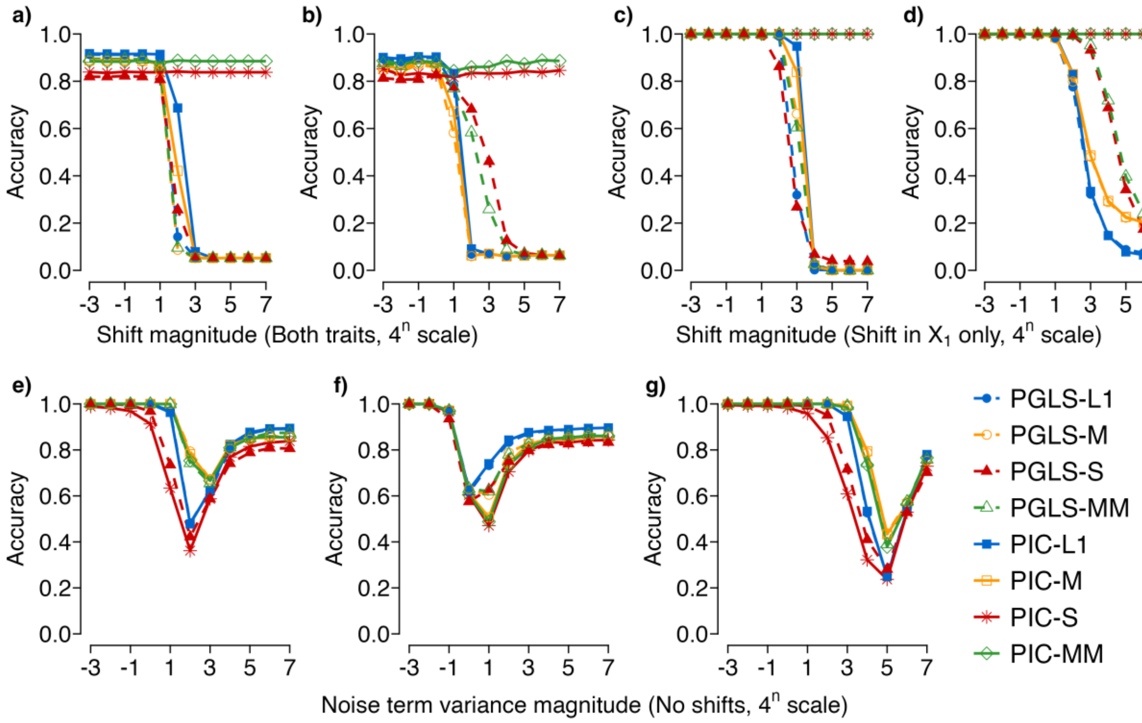


Figure AS6. Evaluation of robust phylogenetic regression methods under the data-pattern–based benchmark on the 128-species random trees. Panel organization and simulation settings are identical to those in Figure AS1, including Felsenstein's worst-case deep-root directional shifts (a, c), randomly located trait shifts (b, d), and no-shift background scenarios, (e) BM–BM, (f) BM–Norm, and (g) Norm–BM. Accuracy is defined as the proportion of simulations in which a method correctly identified both the direction and statistical significance of the trait correlation. The data underlying this figure are available in Supplementary Table S26.

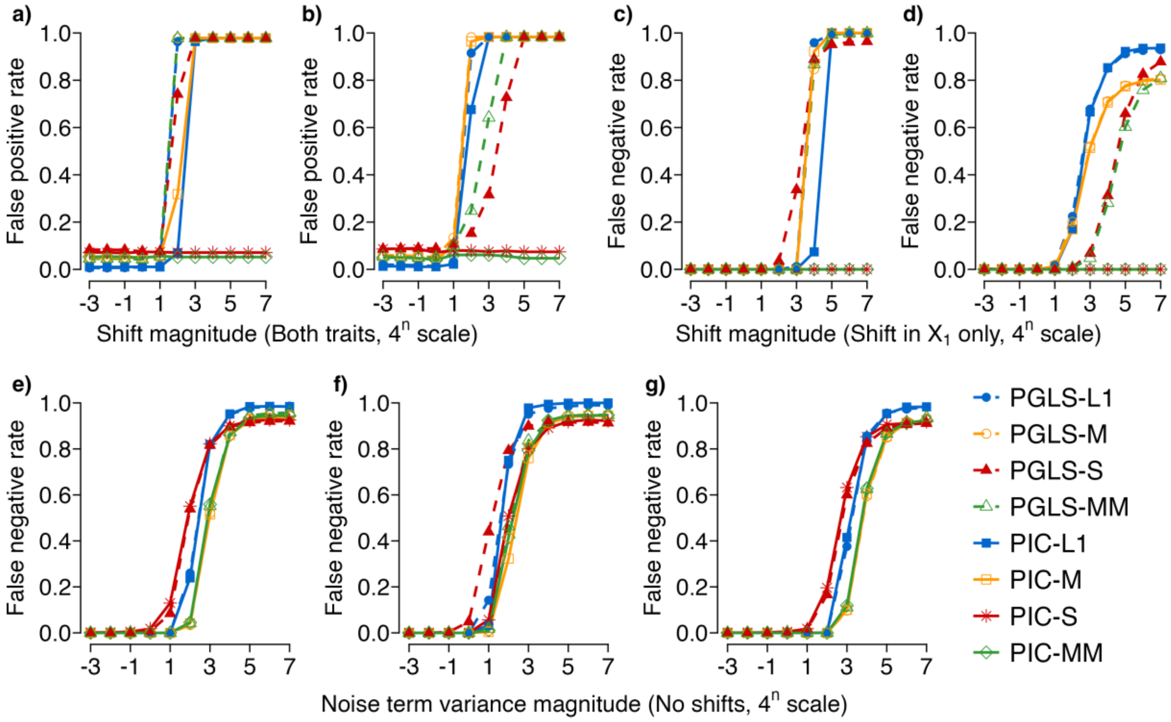


Figure AS7. Evaluation of eight robust phylogenetic regression methods under simulations on a 256-species fixed-balanced tree. Panels a and c correspond to Felsenstein's worst-case scenario, in which strong directional shifts are introduced on deep-root branches. Panels b and d show results from simulations with randomly located trait shifts. Panels a–b report false positive rates under simultaneous shifts in both $X_1$ and $X_2$, whereas panels c–d report false negative rates when only $X_1$ is shifted. Panels e–g show false negative rates under three background scenarios without abrupt shifts: (e) BM–BM, (f) BM–Norm, and (g) Norm–BM. Shift and noise variance magnitudes are expressed on a $4^n$ scale, where n corresponds to the horizontal axis label (e.g., −3 represents $4^{-3}$ and 7 represents $4^7$). False positive and false negative rates are defined relative to the generative parameter $\beta$, using a $\beta$-based benchmark. Additional details on the simulation design, variance gradient configurations, and performance metrics are provided in the Materials and Methods. The data underlying this figure are available in Supplementary Table S27.

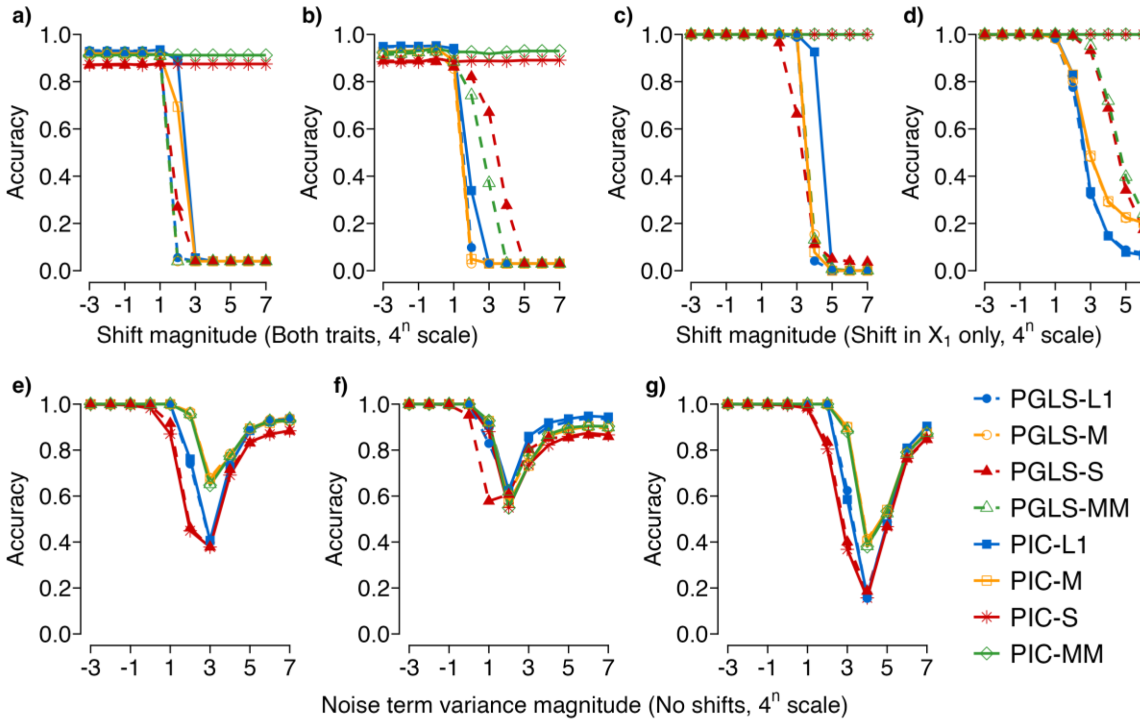


Figure AS8. Evaluation of robust phylogenetic regression methods under the data-pattern–based benchmark on the 256-species fixed-balanced trees. Panel organization and simulation settings are identical to those in Figure AS1, including Felsenstein's worst-case deep-root directional shifts (a, c), randomly located trait shifts (b, d), and no-shift background scenarios, (e) BM–BM, (f) BM–Norm, and (g) Norm–BM. Accuracy is defined as the proportion of simulations in which a method correctly identified both the direction and statistical significance of the trait correlation. The data underlying this figure are available in Supplementary Table S28.

#### S2.5 Summary

These results highlight PIC-MM's robustness and versatility, making it a reliable tool for analyzing phylogenetic data under both stable and dynamic evolutionary conditions. These findings align with the conclusions of Adams et al. (2024). Consequently, PIC-MM was selected as the representative robust regression method for comparison with PIC-O(D)GC. Through this optimization and selection process, we reduced the total number of methods compared with PIC-O(D)GC to six, facilitating a more focused and interpretable evaluation.

#### Reference

Adams R., Cain Z., Assis R., DeGiorgio M. 2024. Robust phylogenetic regression. Syst. Biol., 73:140-157.
